# Computational Tool for Ensemble Averaging of Single-Molecule Data

**DOI:** 10.1101/2020.08.10.241752

**Authors:** Thomas Blackwell, W. Tom Stump, Sarah R. Clippinger, Michael J. Greenberg

## Abstract

Molecular motors couple chemical transitions to conformational changes that perform mechanical work in a wide variety of biological processes. Disruption of this coupling can lead to diseases, and therefore there is a need to accurately measure mechanochemical coupling in motors in both health and disease. Optical tweezers, with nanometer spatial and millisecond temporal resolution, have provided valuable insights into these processes. However, fluctuations due to Brownian motion can make it difficult to precisely resolve these conformational changes. One powerful analysis technique that has improved our ability to accurately measure mechanochemical coupling in motor proteins is ensemble averaging of individual trajectories. Here, we present a user-friendly computational tool, Software for Precise Analysis of Single Molecules (SPASM), for generating ensemble averages of single-molecule data. This tool utilizes several conceptual advances, including optimized procedures for identifying single-molecule interactions and the implementation of a change point algorithm, to more precisely resolve molecular transitions. Using both simulated and experimental data, we demonstrate that these advances allow for accurate determination of the mechanics and kinetics of the myosin working stroke with a smaller set of data. Importantly, we provide our open source MATLAB-based program with a graphical user interface that enables others to readily apply these advances to the analysis of their own data.

**Statement of Significance:** Single molecule optical trapping experiments have given unprecedented insights into the mechanisms of molecular machines. Analysis of these experiments is often challenging because Brownian motion-induced fluctuations introduce noise that can obscure molecular motions. A powerful technique for analyzing these noisy traces is ensemble averaging of individual binding interactions, which can uncover information about the mechanics and kinetics of molecular motions that are typically obscured by Brownian motion. Here, we provide an open source, easy-to-use computational tool, SPASM, with a graphical user interface for ensemble averaging of single molecule data. This computational tool utilizes several conceptual advances that significantly improve the accuracy and resolution of ensemble averages, enabling the generation of high-resolution averages from a smaller number of binding interactions.

## Introduction

Molecular motors generate force and movement in a wide array of cellular processes, including muscle contraction, packaging of DNA into viral capsids, intracellular transport, DNA damage repair, and cell motility. These motors have complex mechanochemical cycles where chemical transitions are coupled to conformational changes in the protein structure that generate mechanical work. The kinetics and mechanics of these transitions are tuned to the specific molecular role of the motor in the cell, and subtle changes in these properties can lead to an array of diseases (1). Therefore, there is a need for experimental and computational techniques for probing these relationships.

Single-molecule optical trapping techniques, with nanometer spatial and millisecond temporal resolution, have proven to be powerful tools for studying the mechanochemical coupling in motors. One widely used optical trapping technique is the three-bead assay (**Fig. 1A**) (2, 3). In this assay, two beads are held in place by dual-beam optical tweezers. The motor’s track (e.g., actin) is strung between these beads and then lowered onto a third, surface-bound bead. This third bead is sparsely coated with motor molecules (e.g., myosin), such that only a single motor interacts with the track at any given time. The positions of the two optically trapped beads are monitored to study the interactions between the motor and the track (**Fig. 1B**), where motor binding to the track causes both displacement of the beads as well as a reduction in the bead variance. This assay has been applied to study several motor and non-motor systems, including dynein (4), the lac repressor (5), kinesins (6), and several myosin isoforms (7–15).

**Figure 1.**
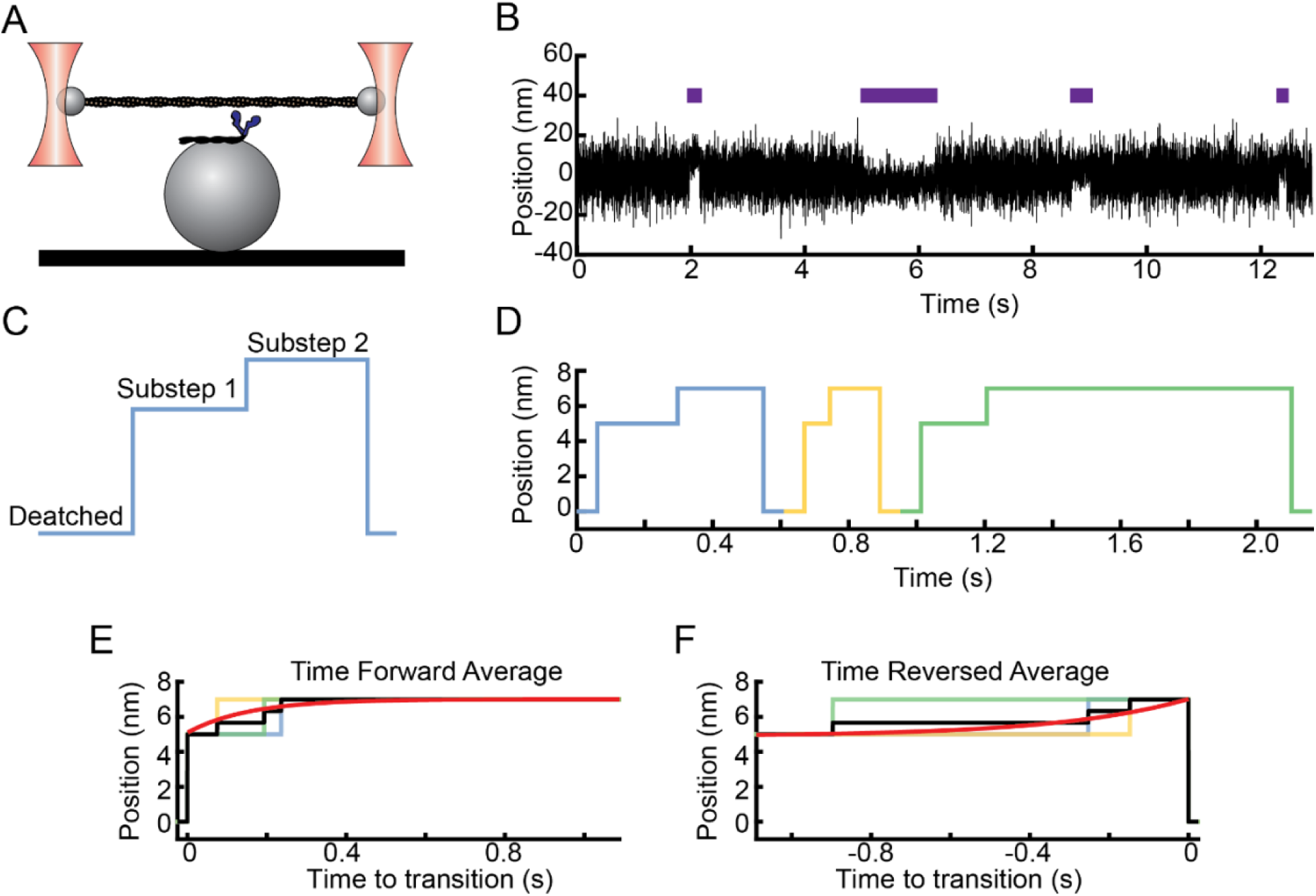
Ensemble averaging of optical trapping data enables the study of mechanochemical coupling. (**A**) Diagram of the three-bead assay, where an actin filament strung between the two optically trapped beads is lowered onto a third surface-bound bead that is sparsely coated with myosin. (**B**) Single-molecule binding interactions between cardiac myosin and actin at 1 µM ATP recorded in the optical trap. The average position between the optically trapped beads is plotted as a function of time, with blue horizontal bars indicating detected binding interactions. The mean position and variance of the beads change upon binding. Brownian motion obscures the second substep of the working stroke. (**C**) Schematic showing the two substeps of the myosin working stroke. (**D**) Idealized trace showing the position over time of a motor with a two-substep working stroke without Brownian motion. (**E**) Procedure for generating time forward ensemble averages from individual binding interactions. Individual trajectories are aligned at the initiation of binding and averaged forward in time (black line), and the average is fit with a single exponential function (red). The y-offset and amplitude of this exponential provide estimates of the average size of the first and second substeps, respectively. The rate of this exponential gives the rate of transitioning from the first substep to the second substep. (**F**) Procedure for generating time reversed ensemble averages from individual binding interactions. Individual trajectories are aligned upon dissociation and averaged backwards in time (black), and the average is fit with a single exponential function (red). The y-offset and amplitude of this exponential provide estimates of the average size of the total step and the second substep, respectively. The rate of this exponential gives the rate of transitioning from the second substep to the detached state.

Analysis of the individual time-dependent trajectories of motor-induced displacements in the bead positions can provide information about both the mechanics and the kinetics of the motor’s mechanochemical cycle. However, it can be difficult to resolve details of these trajectories, as the amplitude of Brownian motion-induced fluctuations in the bead position are frequently larger than the size of motor-induced displacements. One powerful method for extracting high spatial and temporal resolution information from noisy traces is post-synchronization ensemble averaging (13, 16). In this method, trajectories from multiple individual binding interactions are aligned and then averaged together, thereby increasing the signal-to-noise ratio. This technique has been applied to successfully identify substeps of the myosin working stroke (12, 13, 17) and transitions in the ribosome (16) that likely would have been obscured using other analysis methods. While this is a powerful tool for analyzing single-molecule data, there is no software in the public domain that is tailored to performing these calculations, and this has limited the adoption of these tools by many groups.

We have developed a MATLAB-based computational tool, Software for Precise Analysis of Single Molecules (SPASM), with a graphical user interface for the identification and ensemble averaging of single-molecule trajectories. This computational tool utilizes several conceptual advances, including an optimized method for identifying binding interactions from noisy data and improved precision in determining the exact initiation and termination times of binding interactions using a change point algorithm. Using both simulated and experimental data sets, we demonstrate that these advances permit the generation of accurate, high-resolution ensemble averages using fewer individual binding trajectories than were previously required. Our easy-to-use computational tool includes an intuitive graphical user interface and is offered both as open source code and as a standalone program which does not require full installation of MATLAB. Finally, we provide a user guide, a separate tool for simulating data, and sample data sets to help other researchers apply this tool to their own single-molecule data.

## Methods

### Implementation of the computational tool

The SPASM computational tool, which includes a graphical user interface, was written in MATLAB (MathWorks). The program uses the Signal Processing Toolbox and the Optimization Toolbox, but neither toolbox is required for analysis. The code was tested on MATLAB versions R2017b through R2020a for both Windows and macOS operating systems. Standalone versions of the program for both Windows and macOS were generated using the MATLAB Compiler. For more details, see the Supporting Materials.

### Detection of binding interactions

Binding interactions between a motor and its track in the optical trap can be identified using either a variance (18) or a covariance (2, 19) threshold, since the binding of a motor to its track causes a reduction in both the variance and covariance of the two beads (**Fig. 2**). The covariance between the beads at any time, t, is calculated by:

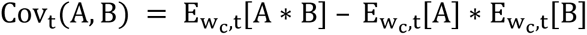

where A is the position of one bead (bead A), B is the position of the other bead (bead B), and E_wc,t_[X] denotes the mean of X over a window of size w_c_ centered at t. Before generating a histogram of covariance values, the covariance is smoothed using a second-order Savitzky-Golay filter with window size w_s_ to remove high-frequency noise. The values of w_c_ and w_s_ can be optimized using the computational tool. See the Supporting Materials for details.

**Figure 2.**
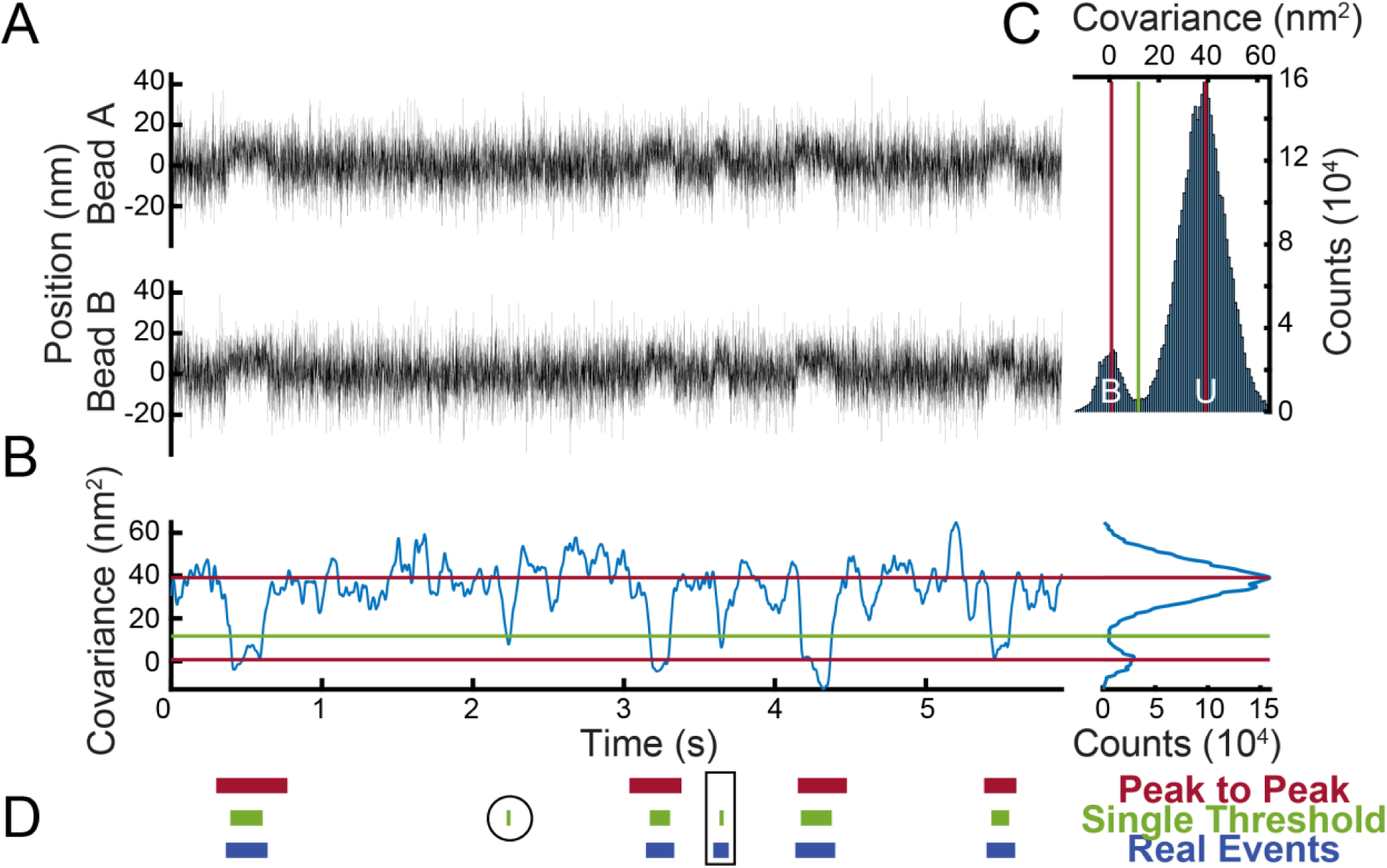
Detection of binding interactions using either the single or peak-to-peak covariance threshold method. (**A**) Simulated optical trapping data showing the position of each optically trapped bead over time. (**B**) Covariance between the position of the optically trapped beads at each time point gives rise to a bimodal distribution. (**C**) A histogram of covariance values shows two distinct populations which correspond to the bound (B) and unbound (U) states. In the single threshold method, a binding interaction is detected when the covariance crosses the value located at the minimum between the two populations (green). In the peak-to-peak method, two thresholds are placed, one at the peak of each population (red), and a binding interaction is detected when the covariance transitions from one threshold to the other threshold. (**D**) Simulated binding interactions detected by the peak-to-peak method (red), binding interactions detected by the single threshold method (green), and actual simulated binding interactions (blue). The single threshold is more susceptible to false positive interactions (circled). The peak-to-peak method is more susceptible to false negative interactions (boxed).

A histogram of the filtered covariance between the two beads shows two distinct populations corresponding to bound (B) and unbound (U) states (**Fig. 2**). This histogram can be used to determine covariance thresholds for detecting binding interactions (10). We use one of two methods to detect binding interactions from the covariance: (1) assigning a single threshold based on the minimum value between the covariance peaks or (2) using a peak-to-peak method which requires that the covariance extend between the bound peak and the unbound peak. The advantages and disadvantages of these methods are discussed in detail in the Results and Discussion.

Once potential binding interactions have been identified, temporal thresholds can be applied to filter the interactions. Any observed reductions in the covariance which are shorter than a user-defined minimum duration are ignored to lower the chance of mistakenly identifying random correlated noise as a binding interaction. Also, any two binding interactions which are separated in time by less than a user-defined minimum separation are ignored to lower the chance of mistakenly identifying random noise as premature detachment between the motor and the track. Note that this filtering takes place after the change points have been located.

### Binding interaction alignment using a change point algorithm and the generation of ensemble averages

Constructing ensemble averages requires the synchronization of individual binding interactions at transitions between the bound and unbound states. Here, we implement a change point algorithm to identify transitions. This algorithm uses maximum likelihood estimation to locate the times, or change points, where changes in both the mean and variance of each bead’s position have most likely occurred. For each binding interaction identified using covariance thresholds, the algorithm searches for the change points within a window of data. For the k^th^ binding interaction, this window spans from

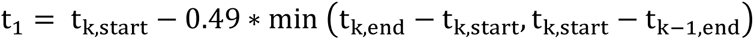

to

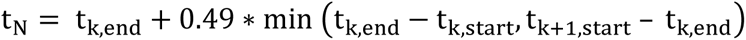

where t_k,start_ and t_k,end_ denote the beginning and end times of the k^th^ interaction as estimated by the covariance threshold method. The window must be wide enough that it includes the entirety of the k^th^ interaction but not so wide that it contains part of another interaction. The computational tool automatically searches the default window for change points, but it also allows for manual adjustment of both the search window and the identified change points.

The algorithm considers the average position between beads A and B during this window. For each pair of time points within the window, (t_i_, t_j_), the algorithm calculates the likelihood that these points coincide with changes in the mean and variance of the data. Each pair divides the window into three intervals: [t_1_, t_i_], [t_i+1_, t_j_], and [t_j+1_, t_N_], where 1 < i < j < N. The log-likelihood score, L_(ti,tj)_, assigned to (t_i_, t_j_) measures how well normal distributions can be fit to these intervals of data:

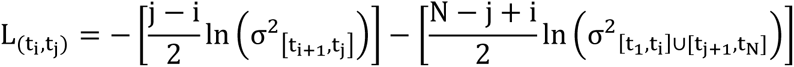

where σ^2^ is the variance of the data during the corresponding interval (see the Supporting Materials for the derivation). L is maximized where the values of t_i_ and t_j_ best divide the window into three sequences of normally distributed data, and these values of t_i_ and t_j_ are then assigned as the change points.

After synchronization at the change points, both time forward and time reversed ensemble averages of individual binding interactions are generated from the average of the two beads’ positions using well-established methods (16). Shorter-lived binding interactions are extended in time to match the duration of the longest-lived binding interaction. The value of this extension equals the average position of the beads during either the first or last 5 ms of the binding interaction for the time reversed and time forward averages, respectively.

### Generation of simulated single-molecule data

To test the accuracy of the program and to aid in the selection of proper window sizes for the analysis of experimental data, we created an additional program to simulate data that resembles single-molecule interactions with user-defined substep sizes and kinetics. The code for this program is provided alongside SPASM so that users can adapt the simulation parameters for their system of interest. Rather than explicitly solving the equations of motion for the optically trapped beads, the parameters used for simulation can be empirically varied until the simulated data matches the experimental data. Trapping data is simulated using a continuous-time Markov jump process in which the motor switches among a baseline detached state and two successive attached states, each with a unique displacement, representing a motor with a two-substep working stroke. The user can set the number of states, the rates of transitioning between the states, and the displacements of each state. High-frequency Gaussian noise is added to simulate Brownian motion. To simulate mechanical coupling between the beads (i.e., higher covariance), a fraction of the noise in each bead’s position, f, is shared between the two beads. When the motor is dissociated from its track, f is set to a larger number so that the motion of the two beads is correlated. When the motor is bound to the track, f is set to a lower number, resulting in a lower covariance. Drift in the system is simulated by the addition of low-frequency noise. For additional details, see the Supporting Materials and the provided code.

### Analysis of simulated data

To test our analysis approach, we generated simulations with well-defined characteristics. Data were simulated with a 2 kHz sampling rate. First, we generated 10 data sets (sets 1-10), each containing 100 binding interactions, to simulate beta cardiac myosin based on previous optical trapping and kinetic measurements (7, 20, 21). The rate of transitioning from the detached state to the first attached state was set to 0.5 s_-1_. The rate of transitioning from the first attached state to the second attached state was set to 70 s_-1_, matching the rate of ADP release (22). The rate of transitioning from the second attached state to the detached state was 4 s_-1_, matching the rate of ATP-induced actomyosin dissociation at 1 µM ATP. The myosin was modeled to have a two-substep working stroke with a 4.7 nm substep followed by a second substep of 1.9 nm (7).

We then generated 10 more data sets to analyze with SPASM (sets 11-20). Each of these sets of data contained 100 simulated binding interactions. The rate of transitioning from the detached state to the first attached state remained at 0.5 s_-1_. The rate of transitioning from the first attached state to the second attached state, however, was much lower at 5 s_-1_, and the rate of transitioning from the second attached state to the detached state was 2 s_-1_. As before, the myosin was modeled to have a two-substep working stroke with a 4.7 nm substep followed by a second substep of 1.9 nm.

With the simulated data, the exact locations of transition points between the bound and unbound states are known, allowing us to test the performance of different analysis methods with regards to: (1) the frequency of false positive binding interactions (i.e., when the bound state is incorrectly detected while the motor is actually unbound), (2) the number of false negative binding interactions (i.e., when the unbound state is incorrectly detected while the motor is actually bound), and (3) the error in determining the correct initiation and termination times of each binding interaction.

To determine the number of false positives, each detected binding interaction was mapped to the nearest overlapping real binding interaction. If a detected binding interaction did not overlap with any real binding interactions, it was counted as a false positive. If multiple detected binding interactions were mapped to the same real binding interaction, all but the closest were also counted as false positives. As we fixed the number of simulated binding interactions within each data set, rather than the total duration of each data set, the data sets typically varied in duration. A longer set of data is expected to result in more false positives, and so the frequency of false positives was calculated by dividing the number of false positives by the duration of the data set. To determine the number of false negatives, each real binding interaction was mapped to the nearest overlapping detected binding interaction. If a real binding interaction did not overlap with any detected binding interactions, it was counted as a false negative. If multiple real binding interactions were mapped to the same detected binding interaction, all but the closest were also counted as false negatives. The error was calculated as the difference between the computationally identified transition points and the nearest simulated transition points for which the corresponding binding interactions overlapped.

### Statistical analysis

Simulated binding interactions were detected using either the single threshold method or the peak-to-peak method, and the frequency of false positives and the number of false negatives were determined. To test for a significant difference in the mean frequency of false positives or the mean number of false negatives between the two methods, p-values were obtained from the independent two-sample t-test. To test if the median error of the detected transition points was significantly changed with the addition of the change point algorithm, p-values were obtained from the Wilcoxon rank sum test.

Ensemble averages were generated from each method of analysis, as well as from the known locations of actual simulated binding interactions. To extract parameters from the ensemble averages, exponential curves were fit to each average, yielding estimates for the substep sizes and rates of the simulated data. For each extracted parameter, a Kruskal-Wallis test was used followed by pairwise Wilcoxon rank sum tests to determine p-values.

### Design of optical trapping apparatus

Experiments were performed on a custom-built, microscope free dual beam optical trap, based on (23, 24). The optical layout is described in the Supporting Materials and Methods (**Fig. S1**). Briefly, the output from a 10 W 1064 nm laser beams (IPG Photonics) was rotated by 45 degrees and then separated into vertically and horizontally polarized components to form 2 independent traps. Optical traps were independently steerable using acoustic optical deflectors (Gooch and Housego) and frequency synthesizer boards under FPGA control (Analog Devices, AD9910 Direct Digital Synthesis evaluation boards). The displacement of the beads from the center of the optical trap was measured at the back focal plane using two quadrant photodiodes (First Sensor). Data were low pass filtered (Frequency Devices) to the Nyquist frequency and digitized on a National Instruments FPGA board (PCIe 7852) with simultaneously sampling analog to digital converters. System control was accomplished by custom software written in LabView. 3D stage control was achieved using a piezoelectric stage (Mad City Labs). Fluorescence was illuminated using the output of a 50 mW 532 laser (Crystalaser). Imaging was performed using an EMCCD camera (Andor).

### Optical trapping experiments

Porcine cardiac myosin and actin were purified from cryoground tissue (Pelfreez) as previously described (25, 26). Bead coated flow cells were assembled as previously described (2, 7, 8). All experiments were performed in KMg25 buffer (60 mM MOPS pH 7.0, 25 mM KCl, 2 mM EGTA, 4 mM MgCl_2_, 1 mM DTT). All buffers and dilutions were prepared fresh each day. Biotin-labeled actin (2 µM) was prepared using 10% biotin actin (Cytoskeleton) in KMg25 buffer. The mixture was allowed to polymerize for 20 minutes, and then the actin was stabilized using tetramethylrhodamine isothiocyanate-labeled phalloidin. Streptavidin beads (Bangs Labs) were washed in 1 mg/mL BSA in KMg25 buffer three times. Flow cells were loaded with myosin (4-20 nM in KMg25 with 200 mM KCl) for 5 minutes and then blocked with 1 mg/mL BSA for 5 minutes. Activation buffer contained KMg25 with the addition of 1 µM ATP, 192 U/mL glucose oxidase, 48 µg/mL catalase, 1 mg/mL glucose, and ∼25 pM Biotin rhodamine-phalloidin actin. A small amount (4 µL) of streptavidin beads were loaded into the flow cell, and the flow cell was sealed with vacuum grease. Trapping experiments were conducted as previously described (2). Two streptavidin beads were optically trapped, forming a bead-actin-bead dumbbell. Trap stiffness was determined by fitting of the power spectral density collected at 20 kHz. The bead-actin-bead dumbbell was pretensed to approximately 2-3 pN and then lowered onto a surface bead to search for binding interactions. Approximately 1 in 5 beads showed binding interactions. Data were acquired at 2 kHz and filtered to 1 kHz.

## Results and Discussion

### Ensemble averaging of single-molecule binding interactions

Ensemble averaging is a powerful method for analyzing single-molecule data, since it can uncover subtle molecular transitions obscured by Brownian motion (13, 16). In ensemble averaging, the time-dependent trajectories of individual binding interactions are synchronized and then averaged. While ensemble averaging techniques are broadly applicable, we will focus in this paper on their application to studying the interaction between myosin molecular motors and actin.

Using ensemble averaging of optical trapping data, it has been shown that many myosin isoforms have a two-substep working stroke, where the first substep corresponds to the release of inorganic phosphate and the second substep corresponds to a transition associated with ADP release (**Fig. 1C-D**) (7-10, 12-14, 17, 27). It is difficult to distinguish the second transition from raw data traces due to Brownian motion. However, ensemble averaging allows for easier visualization of this transition by increasing the signal-to-noise ratio.

One can collect information about both the kinetics and mechanics of the working stroke substeps from the post-synchronized ensemble averaged trajectories of individual binding interactions (13, 16). These interactions can be synchronized upon actomyosin attachment and then averaged forward in time or, alternatively, synchronized upon actomyosin detachment and then averaged backward in time (**Fig. 1E-F**). The magnitude of the initial displacement seen in the time forward averages gives the size of the first substep of the myosin working stroke, a transition which occurs within the dead time of typical optical tweezer instruments. The amplitude of the subsequent exponential rise in displacement in the time forward averages gives the size of the second substep of the working stroke. The rate of this exponential rise is the rate of transitioning from the first substep to the second substep, and it is associated with ADP release in myosins (13). For the time reversed ensemble averages, the exponential rise in displacement prior to detachment has an amplitude equal to the size of the second substep, and the rate of this exponential gives the rate of transitioning from the second substep to the detached state, a transition which corresponds to ATP-induced actomyosin dissociation (13).

### MATLAB-based computational tool for generating ensemble averages

Here, we have generated an easy-to-use MATLAB-based computational tool, SPASM, which finds binding interactions within noisy data, accurately identifies transitions between the bound and unbound states, and then generates ensemble averages. This tool includes several improvements and optimized procedures for both the identification and alignment of binding interactions, which are discussed below. The tool features a graphical user interface for ease of use and is packaged with an accompanying user guide. We provide the code for this tool as well as a compiled executable file that does not require a full installation of MATLAB. We also provide a resource for simulating single-molecule data, as well as the sample simulated data sets used in our analysis (see Supporting Materials).

### Generation of covariance histogram to identify binding interactions

The first step in generating ensemble averages is the identification of binding interactions from single-molecule data traces. When optically trapped, the two beads in the bead-actin-bead dumbbell undergo fluctuations in their position due to Brownian motion (**Fig. 2A**). The motion of these beads is mechanically coupled through the actin filament, as evidenced by the covariance between their positions (**Fig. 2B**). When the surface-bound motor binds to the actin filament, it causes several pronounced changes: (1) it reduces the positional variance of each bead’s position, (2) it reduces the coupled motion (covariance) of the two trapped beads, and (3) it displaces the mean position of each bead. The majority of analysis methods for identifying binding interactions utilize the changes in the mean position, variance, and/or covariance of the optically trapped beads upon binding to actin (11, 18, 19, 24, 28).

One popular method for selecting binding interactions is to set a threshold based on the variance or covariance of the beads. The choice of using a variance or covariance threshold for binding interaction identification will partially be dictated by the optical trap layout. For systems which only monitor the position of a single bead, one must use a variance threshold for the position of the single bead. For systems where both bead positions are monitored, a covariance threshold is preferred since it is less sensitive to noisy fluctuations in the data. While we focus on the use of our computational tool with a covariance threshold, the same approaches and conclusions will hold true for a variance threshold based on the position of one bead. A version of SPASM that uses a variance threshold is provided (see Supporting Materials).

Our computational tool identifies binding interactions from the change in the covariance between the positions of the two trapped beads that occurs upon myosin binding to actin. SPASM first calculates the covariance over a sliding window in time and then smooths the covariance over a separate window. With properly chosen window lengths, the histogram of the covariance values reveals two populations (**Fig. 2C**), where the higher covariance population corresponds to unbound states and the lower population corresponds to bound states (2). One can then select binding interactions based upon thresholds that distinguish between these two populations (see *Selection of binding interactions* below).

The success of this approach depends on the degree of separation between the two peaks in the covariance histogram. If the peaks are not well separated, the analysis is more susceptible to false and/or missed binding interactions. The ability to generate a histogram with two well separated peaks depends partly on the selection of proper window lengths for the calculation and smoothing of the covariance. Optimal values for these parameters, in turn, depend on the kinetics of the myosin’s interaction with actin, the compliance of the myosin and/or myosin-surface attachment, the pretension between the optically trapped beads, and the noise in the system. Therefore, the window lengths often need to be determined empirically. If the kinetics of the myosin’s transitions are known from other experimental measurements, one can simulate data and select window lengths which optimize analysis of the simulated data (see Supporting Materials). If kinetic information about the myosin’s transitions is unknown, it may not be possible to generate meaningful simulated data. In these cases, the window lengths can be determined empirically through the computational tool’s graphical user interface, which allows the user to vary the window lengths until a suitable bimodal covariance histogram is achieved.

### Selection of binding interactions

Once a suitable covariance histogram with two well-defined peaks has been generated, the next step is to determine proper thresholds for the covariance which will be used to detect binding interactions. One possibility for distinguishing the bound state from the unbound state is to use a single covariance threshold located at the minimum value between the two peaks of the covariance histogram (10). Here, detected interactions start when the covariance drops below this threshold value, and they end when the covariance rises back above this threshold value (**Fig. 2D**). Alternatively, one could identify the binding interactions using a set of two different covariance thresholds, located at the two peaks of the covariance histogram. In this ‘peak-to-peak’ approach, a binding interaction is considered to start when the covariance drops from the threshold defined by the unbound peak to the threshold defined by the bound peak. Likewise, a binding interaction is considered to end when the covariance rises from the threshold defined by the bound peak to the threshold defined by the unbound peak (**Fig. 2D**).

We tested the abilities of the single threshold and peak-to-peak methods to accurately detect simulated binding interactions between actin and cardiac myosin. Interactions were simulated using a continuous-time Markov jump process with kinetics and mechanics based on previously measured parameters for ventricular cardiac myosin (7, 21, 22) (see Materials and Methods for details). With simulated data, the exact locations of the binding interactions are known, allowing for easy comparison between the simulated interactions and the interactions detected by the computational tool using either method (**Fig. 2D**).

We generated 10 independent sets of simulated data, each containing 100 binding interactions (sets 1-10). For each data set, we used our computational tool to calculate the covariance histogram, locate the peaks and minimum of the histogram, and identify binding interactions using either the single threshold method or the peak-to-peak method. When we used a single threshold to identify binding interactions, we correctly detected 80 +/- 4 of the 100 binding interactions on average, and we incorrectly detected 4 +/- 1 false positive binding interactions per 100 seconds of data, on average (**Table 1**). The reported errors are standard deviations. When we used the peak-to-peak method to identify binding interactions, we correctly detected 65 +/- 5 of the 100 binding interactions on average, and we did not detect any false positive binding interactions. Although the peak-to-peak method misses a greater number of binding interactions, the false positive rate is lower for this method (p < 0.001).

**Table 1.**
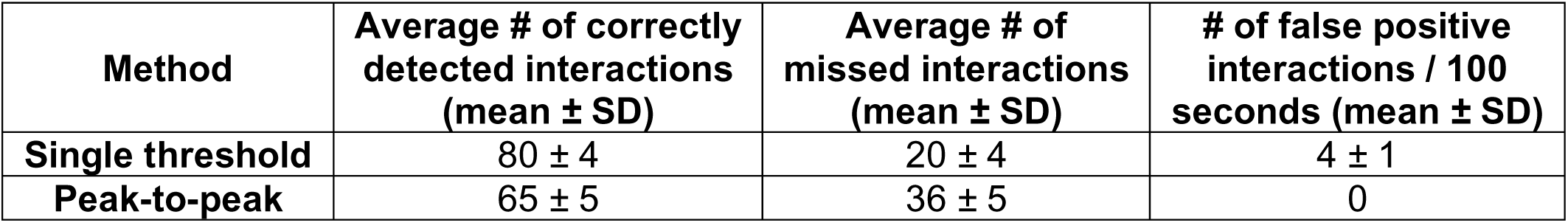
Detection of binding interactions using either the single or peak-to-peak covariance threshold method. Average number of correctly identified binding interactions and frequency of false positive binding interactions detected with the single threshold method and peak-to-peak method for 10 data sets, each containing 100 simulated binding interactions (sets 1-10). Calculated values were rounded to the nearest whole number.

| **Method** | **Average # of correctly detected interactions (mean ± SD)** | **Average # of missed interactions (mean ± SD)** | **# of false positive interactions / 100 seconds (mean ± SD)** |
| --- | --- | --- | --- |
| **Single threshold** | 80 ± 4 | 20 ± 4 | 4 ± 1 |
| **Peak-to-peak** | 65 ± 5 | 36 ± 5 | 0 |

A single threshold could work well for selecting binding interactions if the two populations of the histogram are sufficiently distinct. However, it is often not possible to obtain sufficient separation between the peaks due to factors that lower the signal-to-noise ratio (e.g., system noise, insufficient pretension between the beads, fast binding kinetics). In these cases, this single threshold approach is prone to identifying false positive interactions, where the covariance crosses the threshold even though the actomyosin has remained in an unbound state. These false positive binding interactions do not generate a net displacement in the optical trap, and so their inclusion in the ensemble averages is expected to lead to an underestimation of the true size of the working stroke. A methodology has been developed which attempts to correct for these false positive interactions through the use of normalization factors (10). Alternatively, as the vast majority of these false positive interactions arise due to either Brownian motion (or system noise) induced rapid downward spikes in the covariance (which lead to very short detected interactions) or rapid upward spikes in the covariance (which lead to multiple detected interactions in quick succession), it is possible to avoid these false positive interactions through the use of temporal filters that exclude interactions which are too short or pairs of interactions which are too close to one another. However, it is not always easy to determine appropriate values for these temporal filters. Further, the use of these temporal filters may lead to the exclusion of many correctly identified binding interactions. When we used optimized values for these filters to exclude all of the false positive interactions that were detected by the single threshold method, we were left with fewer interactions than were detected by the peak-to-peak method (**Fig. S2**).

With the peak-to-peak method, the criteria for detecting a binding interaction is much stricter than with the single threshold method, and the number of identified false positive binding interactions is expected to decrease while the number of missed, short-lived binding interactions increases. Unlike the inclusion of false positive interactions, the exclusion of these missed binding interactions does not adversely affect the size or shape of the ensemble averages. Although we demonstrate that the peak-to-peak method performs better in data traces with moderate separation between the peaks of the covariance histogram, some experimental data might have better peak separation. In this case, the single threshold method would be preferable since it maximizes the number of captured binding interactions. The computational tool allows the user to try both methods, and it automatically determines appropriate values for the thresholds.

### Alignment of binding interactions using covariance thresholds

After binding interactions are identified, they must be precisely aligned at the transitions between the bound and unbound states to generate accurate ensemble averages. The most critical step in aligning these interactions is the careful determination of when exactly a transition occurs. Inadequate determination of these transitions will lead to inaccurate measurements of the substep sizes and/or kinetics. Several methods have been applied to locate transitions in single-molecule data traces, including Hidden Markov Models (28) and step finding algorithms (29), but a frequently used method for post-synchronization is to align the binding interactions based on the same thresholds used to identify the binding interactions (2, 10, 13).

To test the abilities of the single threshold and peak-to-peak methods to accurately identify the transitions, we used the same 10 simulated data sets containing 100 transitions each, as described previously (sets 1-10). When we used a single threshold to identify transition times, we found that the detected attachment times occurred 28.2 (95% confidence intervals: +13.8, −21.7) milliseconds after the actual attachment times, on average (**Table 2**), and the detected detachment times occurred 28.6 (+11.9, −19.1) milliseconds before the actual detachment times, on average. On the other hand, when we used the peak-to-peak method to identify transitions, we found that the detected attachment times occurred 55.5 (+195.5, −69.0) milliseconds before the actual attachment times, on average, and the detected detachment times occurred 50.4 (+188.1, −64.9) milliseconds after the actual detachment times, on average. Taken together, the single threshold method has better temporal resolution when identifying transitions between the bound and unbound states.

**Table 2.**
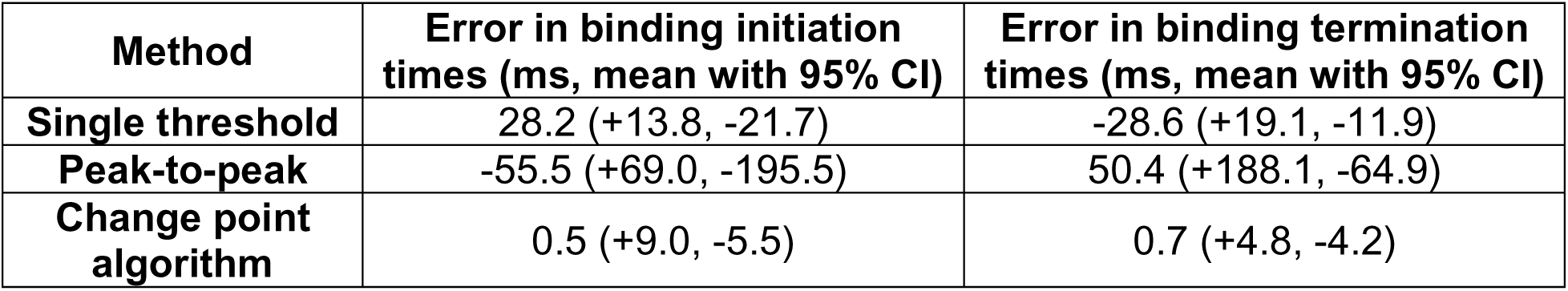
The change point algorithm minimizes the error when detecting the locations of transitions. Mean and 95% confidence intervals for the error when detecting transitions within simulated data sets 1-10 with the single threshold method, the peak-to-peak method, and the change point algorithm. When estimating the binding initiation times, 645 of 1000 transitions were detected and analyzed for the peak-to-peak method, 598 transitions were detected and analyzed for the single threshold method, and 644 transitions were detected and analyzed for the change point algorithm. The same number of transitions were detected and analyzed for each method when estimating the binding termination times. Note that a negative average error indicates that the detected transitions occurred before the actual transitions, on average.

When binding interactions are aligned based on the covariance thresholds, it is assumed that the covariance drops and rises in conjunction with transitions between the bound and unbound states. With the single threshold method, this is a fairly reasonable assumption, explaining why it outperforms the peak-to-peak method. Each true transition point separates more highly correlated bead motion (i.e., the unbound state) from less highly correlated bead motion (i.e., the bound state). The covariance is calculated over a window, so when the covariance window is centered at a transition point, the window will include equal amounts of more highly and less highly correlated data. The covariance at the transition point should then lie at some intermediate value between the two peaks of the covariance histogram. However, the single threshold method is not perfect at locating the transition points. First, while the value of the covariance at a transition point will likely be near the minimum value between the two peaks of the covariance histogram, there is no guarantee that it will lie exactly at this minimum value. Additionally, synchronized large-scale movement of both beads due to the myosin’s power stroke can produce transient spikes in the covariance value during transitions, and these spikes can potentially decrease the accuracy of the single threshold method in identifying exact transition times.

The peak-to-peak method produced poorer alignment than the single threshold method. When the peak-to-peak method is used to identify transitions, it is assumed that transitions occur when the covariance crosses the upper threshold, defined by the position of the unbound peak. This is inherently less accurate for estimating transition points than the single threshold method. A window of data which has a covariance value that is similar to the value of the unbound peak contains primarily correlated data and, therefore, it is unlikely that the center of this window is near the actual transition point. In fact, the calculated transition point using the peak-to-peak method would be expected to deviate from the actual transition point by at least half the window size.

Taken together, our data show that when binding interactions are synchronized using a single covariance threshold, the resulting ensemble averages are expected to have better alignment of binding interactions. However, as noted previously, the use of a single covariance threshold to detect binding interactions is more susceptible to false positive binding interactions which would lead to an underestimation of the true substep sizes. The peak-to-peak method is better for binding interaction detection without including false positives, but it lacks the necessary temporal resolution to accurately align the detected interactions.

### Change point algorithm for aligning interactions

Rather than relying on the covariance when estimating transition times, we tested the use of separate methods for detecting and synchronizing binding interactions. To improve our ability to locate the transition times of each binding interaction, we implemented a change point algorithm (see Materials and Methods for details). Change point algorithms have been used in step finding for transitions in biological processes, where the algorithm identifies the most likely times in which there was a change in a parameter such as motor position or rotation of the myosin lever arm (29, 30). We have adapted the change point algorithm for the three-bead assay, where we search for the most likely transition times based on changes in both the mean and the variance of the bead positions, as both of these parameters differ between the bound and unbound states (**Fig. 3A**). For each binding interaction identified by the covariance threshold method (**Fig. 3B**), our algorithm examines the positions of the trapped beads in a window surrounding that interaction and finds the two points (i.e., binding initiation and detachment) within this window that most likely represent transitions in the mean and variance of the data (**Fig. 3C**; see Methods for details).

**Figure 3.**
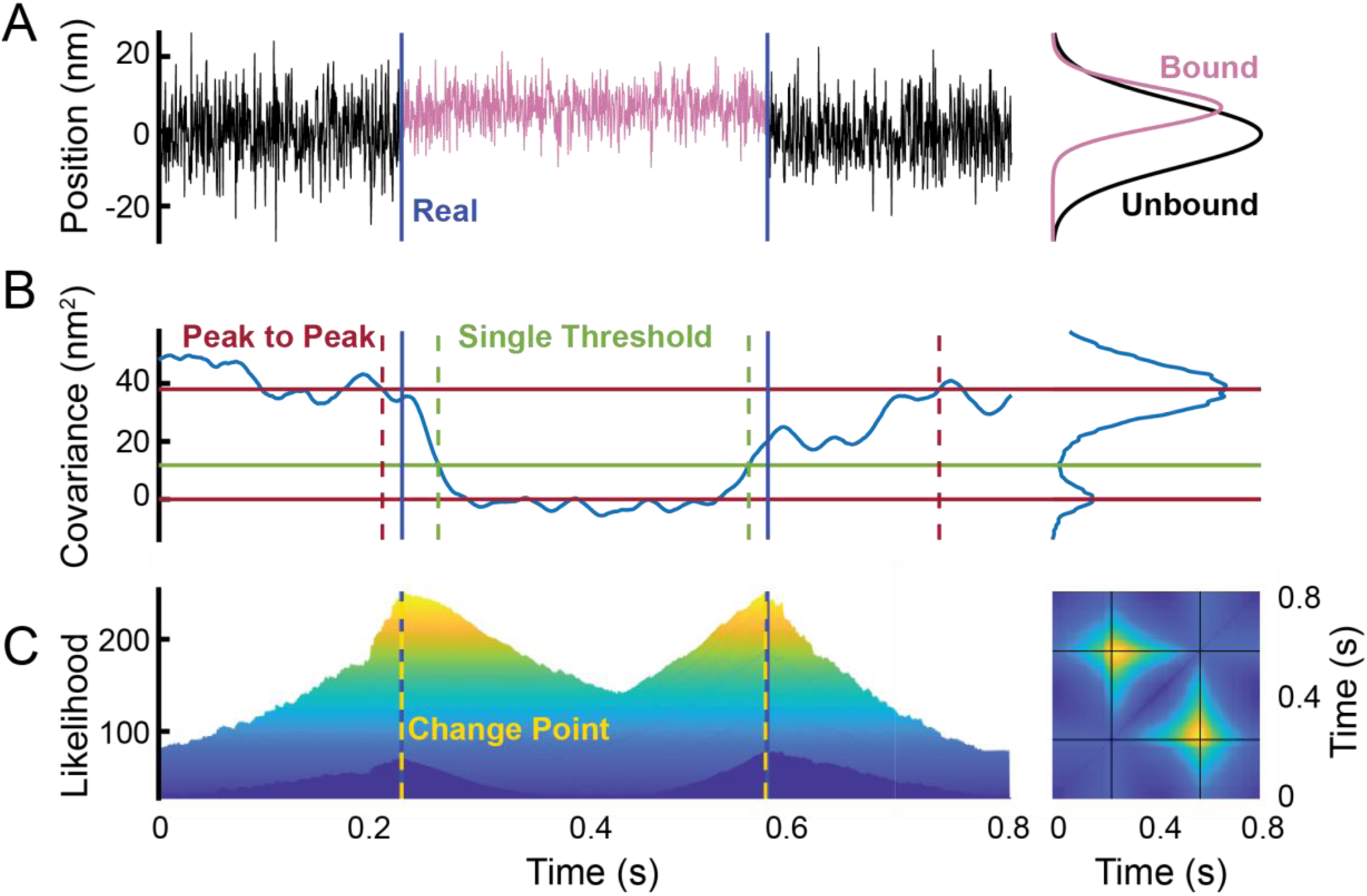
The change point algorithm more precisely identifies transitions between bound and unbound states. (**A**) Simulated optical trapping data showing the average position between the optically trapped beads over time during a binding interaction. Data obtained during the bound state (purple) are drawn from a normal distribution with a shifted mean and a lower variance when compared to data obtained during the unbound state (black). The change point algorithm seeks to find the time points which best separate the two distributions. The locations of the actual simulated transitions are marked with blue vertical lines. (**B**) Calculated covariance of the bead positions during the simulated binding interaction in (A). The attachment and detachment times identified by the single threshold (green) and the peak-to-peak (red) methods are shown with dashed vertical lines. The actual transitions are marked with solid blue vertical lines. (**C**) The change point algorithm determines the likelihood that any two points within an extended search window are the two transition points. (left) Plot of the likelihood assigned to each pair of points within the search window, viewed from the side (see Materials and Methods for details). The change points, which occur when the likelihood is maximized, are shown with dashed yellow vertical lines, while the actual transitions are marked with solid blue vertical lines. (right) The likelihood viewed from above. Regions of yellow correspond to higher likelihood, while regions of dark blue correspond to lower likelihood. The two change points are marked with solid black lines.

To test the ability of the change point algorithm to accurately identify transition times, we again analyzed the same 10 sets of simulated data described above (sets 1-10). We found that the attachment times detected by the change point algorithm occurred 0.5 (+9.0, −5.5) milliseconds after the actual attachment times, on average (**Table 2**), and the detachment times detected by the change point algorithm occurred 0.7 (+4.8, −4.2) milliseconds after the actual detachment times, on average (**Table 2**). Statistical testing demonstrates that the change point algorithm outperforms both the single threshold method (p_start_ < 0.001, p_end_ < 0.001) and the peak-to-peak method (p_start_ < 0.001, p_end_ < 0.001) in identifying transition times. As our simulated data were generated with a sampling frequency of 2 kHz, these average errors of about 0.5 ms indicate that the change point algorithm was typically correct within 1 point. It is possible that a higher sampling frequency would further increase the accuracy.

To explore the ability of these three methods to accurately identify transition points, we generated cumulative distributions of the differences between the detected transition times and the actual simulated transition times for both the initiation and termination of the binding interactions (**Fig. 4**). Here, the width of the distribution reveals the precision of the corresponding method, while the sign and magnitude of the average error reveals the systematic bias of that method. As expected, the cumulative distributions of errors generated from the peak-to-peak method are wide, indicating low precision at identifying the transitions, while the distributions generated from the single threshold method are narrower, indicating higher precision. The distributions generated from the change point algorithm are very narrow, and the mean error is close to 0. This indicates that the change point algorithm is very precise and has lower systematic bias than either the single threshold or peak-to-peak method.

**Figure 4.**
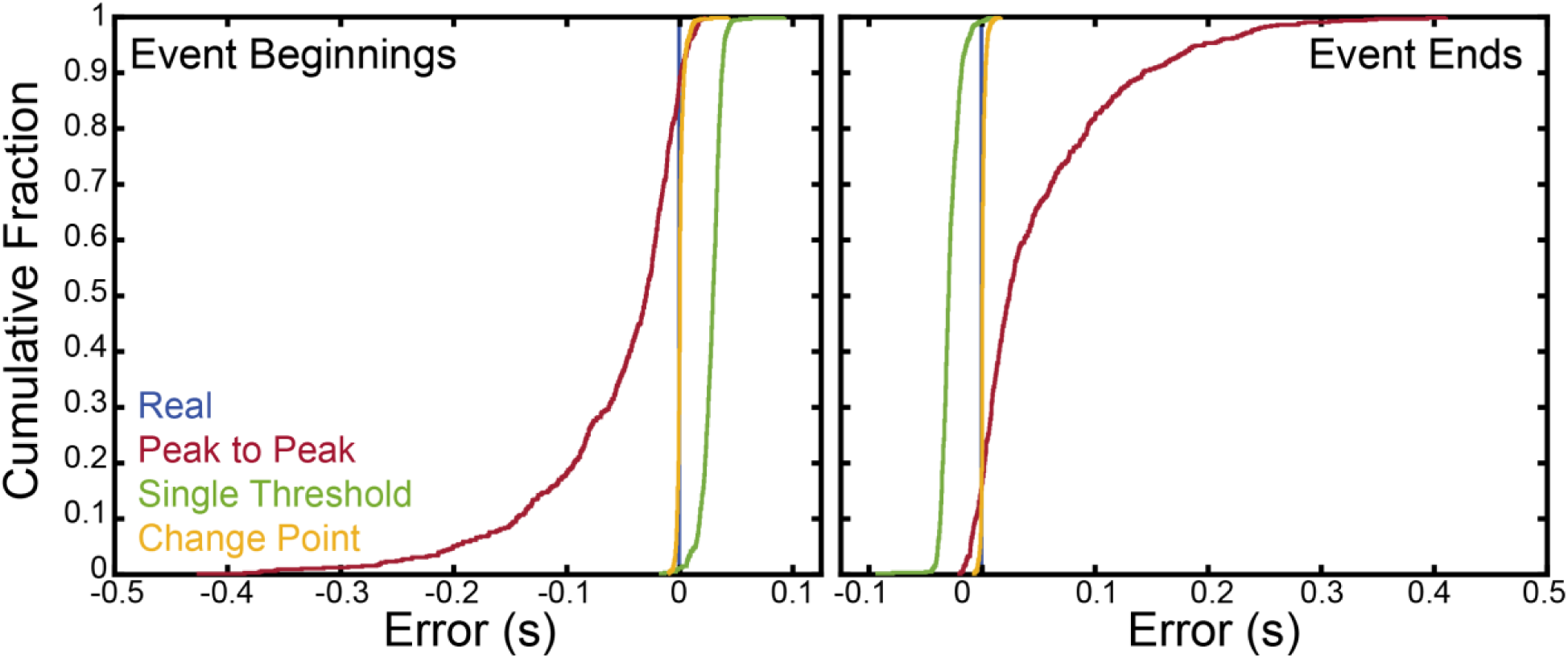
The change point algorithm minimizes the error when detecting the locations of transitions. The error was calculated as the difference between the detected binding times and the actual simulated binding times for simulated data (sets 1-10). (left) Cumulative distributions of the errors in determining the binding initiation times using the peak-to-peak method (red), the single threshold method (green), and the change point algorithm (yellow). Statistical comparisons can be found in Table 2. (right) Cumulative distributions of the errors when determining the binding termination times using the peak-to-peak method (red), the single threshold method (green), and the change point algorithm (yellow).

### Comparison of ensemble averages generated using different methods

To test our predictions about the relative accuracy of the ensemble averages when using each method of analysis, we generated ensemble averages from the 10 sets of simulated data studied previously (sets 1-10). First, we generated ensemble averages using the actual locations of all 1000 simulated binding interactions to align the binding interactions (**Fig. 5A-B**, real). We also generated ensemble averages for each of the 10 sets of data, using the actual locations of the 100 simulated binding interactions within each set. Exponential curves were fit to each of these averages to estimate the substep sizes and rates of the simulated myosin working stroke (**Fig. 5C-F**, real; **Table 3**). The magnitude of substep 1 estimated from the time forward averages was 4.7 (95% confidence intervals: +0.4, −0.4) nm, on average, while the magnitude of the total step estimated from the time forward averages was 6.4 (+0.2, −0.2) nm, on average. The magnitude of substep 1 estimated from the time reversed averages was 5.7 (+0.2, −0.3) nm, on average, while the magnitude of the total step estimated from the time reversed averages was 6.5 (+0.1, −0.2) nm, on average. The estimated rate of transitioning from the first substep to the second substep (k_f_) was 68.7 (+15.8, −20.9) s_-1_, and the estimated rate of transitioning from the second substep to the detached state (k_r_) was 4.3 (+2.2, −1.9) s_-1_.

**Figure 5.**
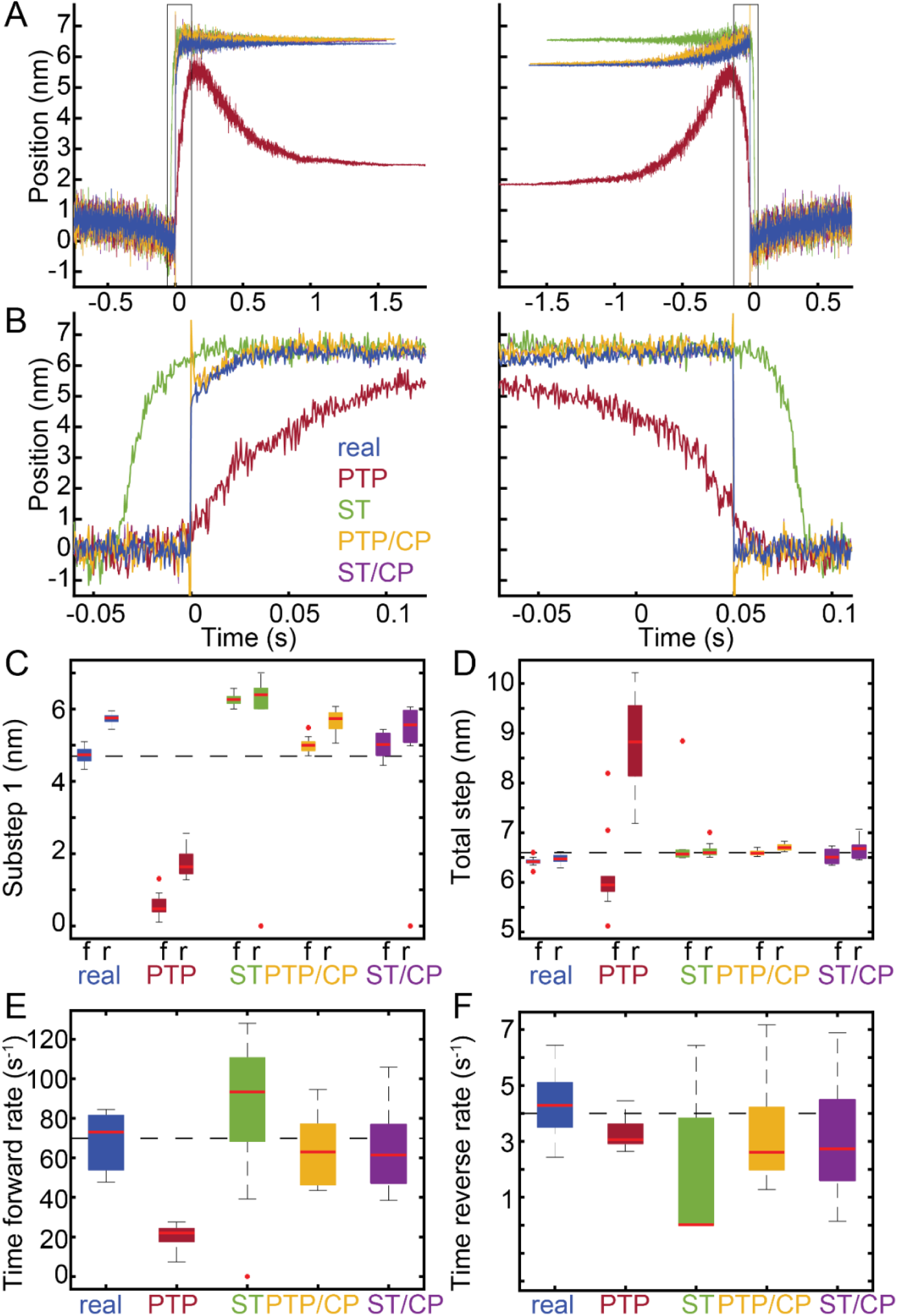
Ensemble averages of simulated binding interactions. 10 sets of data were simulated, each containing 100 binding interactions (sets 1-10). Interactions were detected using either the peak-to-peak (PTP) or the single threshold (ST) method, and interactions were aligned using either the transitions estimated by the covariance threshold method or the change points identified by the change point algorithm (CP). (**A**) (left) For each analysis method, all detected binding interactions were aligned at the estimated initiation times and averaged together to generate time forward ensemble averages. (right) For each analysis method, all detected binding interactions were aligned at the estimated termination times and averaged together to generate time reversed ensemble averages. Also shown are the time forward and time reversed ensemble averages generated from the known locations of the actual simulated binding interactions (blue, real). (**B**) A zoomed in view of the boxed segments of the ensemble averages in A highlights the misalignment in the averages when the change point algorithm is omitted. (**C-F**) For each of the 10 simulated sets of data containing 100 binding interactions, ensemble averages were generated and fit with single exponential functions. The substep sizes and rates of the simulated myosin working stroke were estimated from the exponential fits. Box plots show the estimated parameters for each analysis method. Outliers are indicated by red dots. The substep sizes were estimated from both the time forward (f) and the time reversed (r) ensemble averages. Horizontal dashed lines show the values of the simulated parameters. Statistical analysis for each parameter can be found in **Table 3**.

**Table 3.** The change point algorithm improves ensemble averages. 10 sets of data were simulated, each containing 100 binding interactions (sets 1-10). Interactions were detected using either the peak-to-peak or the single threshold method, and interactions were aligned using either the transitions estimated by the covariance threshold method or the change points identified by the change point algorithm. For each data set, ensemble averages were generated using either the known locations of actual simulated binding interactions (real) or using the binding interactions detected by each method of analysis. The averages were fit with exponential functions, and the substep sizes and rates of the simulated myosin working stroke were estimated from the rates and amplitudes of the exponential fits. (top) Mean and 95% confidence intervals for the size of substep 1, the size of substep 2, the total step size, and the rate of transitioning from the first substep to the second substep, as estimated by the time forward ensemble averages. (bottom) Mean and 95% confidence intervals for the size of substep 1, the size of substep 2, the total step size, and the rate of transitioning from the second substep to the detached state, as estimated by the time reversed ensemble averages. The p-value for a given set of parameter values estimated by a given analysis method was obtained from the Wilcoxon rank sum test between those estimated parameter values and the values estimated by using the known locations of actual simulated binding interactions (real).

| Time forward ensemble averages (mean with 95% CI) |  |  |  |  |  |
| --- | --- | --- | --- | --- | --- |
| Parameter | real | PTP | ST | PTP, CP | ST, CP |
| **Substep 1 (nm)** | 4.7<br>(+0.4, -0.4) | 0.6<br>(+0.7, -0.5)<br>p < 0.001 | 6.3<br>(+0.3, -0.3)<br>p < 0.001 | 5.0<br>(+0.5, -0.3)<br>p = 0.021 | 5.0<br>(+0.4, -0.6)<br>p = 0.045 |
| **Substep 2 (nm)** | 1.7<br>(+0.5, -0.4) | 5.6<br>(+1.7, -0.8)<br>p < 0.001 | 0.5<br>(+1.7, -0.4)<br>p = 0.003 | 1.6<br>(+0.3, -0.4)<br>p = 0.427 | 1.5<br>(+0.5, -0.6)<br>p = 0.186 |
| **Total step (nm)** | 6.4<br>(+0.2, -0.2) | 6.2<br>(+2.0, -1.1)<br>p = 0.026 | 6.8<br>(+2.1, -0.3)<br>p = 0.002 | 6.6<br>(+0.1, -0.1)<br>p = 0.001 | 6.5<br>(+0.2, -0.2)<br>p = 0.186 |
| **Rate (s<sup>-1</sup>)** | 68.7<br>(+15.8, -20.9) | 20.2<br>(+7.4, -12.8)<br>p < 0.001 | 84.3<br>(+43.8, -84.3)<br>p = 0.141 | 64.5<br>(+30.0, -20.9)<br>p = 0.473 | 63.6<br>(+42.3, -25.0)<br>p = 0.241 |
| Time reversed ensemble averages (mean with 95% CI) |  |  |  |  |  |
| Parameter | real | PTP | ST | PTP, CP | ST, CP |
| **Substep 1 (nm)** | 5.7<br>(+0.2, -0.3) | 1.7<br>(+0.8, -0.4)<br>p < 0.001 | 5.2<br>(+1.8, -5.2)<br>p = 0.026 | 5.7<br>(+0.4, -0.6)<br>p = 0.970 | 5.0<br>(+1.0, -5.0)<br>p = 0.385 |
| **Substep 2 (nm)** | 0.7<br>(+0.2, -0.2) | 7.1<br>(+1.7, -1.4)<br>p < 0.001 | 1.4<br>(+5.2, -1.4)<br>p = 0.038 | 1.0<br>(+0.5, -0.4)<br>p = 0.026 | 1.6<br>(+4.8, -1.1)<br>p = 0.045 |
| **Total step (nm)** | 6.5<br>(+0.1, -0.2) | 8.8<br>(+1.4, -1.6)<br>p < 0.001 | 6.6<br>(+0.4, -0.1)<br>p = 0.003 | 6.7<br>(+0.1, -0.1)<br>p < 0.001 | 6.7<br>(+0.4, -0.2)<br>p = 0.011 |
| **Rate (s<sup>-1</sup>)** | 4.3<br>(+2.2, -1.9) | 3.3<br>(+1.2, -0.6)<br>p = 0.054 | 1.7<br>(+4.8 -1.7)<br>p = 0.038 | 3.3<br>(+3.9, -2.0)<br>p = 0.089 | 3.0<br>(+3.8, -2.9)<br>p = 0.076 |

We then used either the single threshold method or the peak-to-peak method to detect binding interactions within each data set. When the single threshold method was used to detect binding interactions, we applied a filter to ignore any detected interactions which were shorter than 77 ms or within 63 ms of another detected interaction, to avoid including false positive interactions (**Fig. S2**; **Fig. S3** shows the effect of including these false positive binding interactions). To identify transitions between the bound and unbound states for each interaction, we either included or omitted the change point algorithm. For each of these analysis methods, we used the binding interactions and transitions detected over all 10 data sets to generate ensemble averages (**Fig. 5A-B**). As before, we also generated ensemble averages from the binding interactions detected within each of the 10 sets of data, and exponential curves were fit to each average to estimate the substep sizes and rates of the simulated myosin working stroke (**Fig. 5C-F**; **Table 3**). As expected, using the change point algorithm to align the binding interactions resulted in the most accurate estimates.

When the peak-to-peak method was used to both detect and align the binding interactions, the ensemble averages were misshapen (**Fig. 5**, PTP). The time forward average, for example, includes the characteristic increase in displacement but then drops. This drop is due to the fact that the binding interaction termination times detected by the peak-to-peak method often came after the actual termination times, leading to the inclusion of baseline data at the end of the time forward average. The time forward average also appears to start too late, as the peak-to-peak method typically guesses that binding initiation times occur before they actually do (**Fig. 4**). Exponential curves were very poorly fit to these ensemble averages.

When the single threshold method was used to both detect and align the binding interactions, the ensemble averages had better overall shape (**Fig. 5**, ST). However, similar to the averages generated with the peak-to-peak method, misalignment among the individual trajectories resulted in very gradual transitions between the bound and unbound states. The time forward average, for example, appears to start too early, as the single threshold method typically guesses that binding initiation times occur after they actually do (**Fig. 4**).

When the change point algorithm was used to align the binding interactions, the ensemble averages featured much sharper transitions (**Fig. 5**, PTP/CP and ST/CP). However, very sharp spikes in displacement occur at the transition times (**Fig. 5A-B**, PTP/CP and ST/CP). Brownian motion-driven fluctuations in the bead positions can cause changes in the data from one point to the next which are not due to transitions between the bound and unbound states. If such noise happens to occur near a real transition point, it offers an attractive candidate for the change point, and the change point algorithm may choose that point instead of the less pronounced yet correct transition time. However, we have shown that the transition times estimated by the change point algorithm are within 1 to 2 points of the actual simulated transition times, on average (**Table 2**; **Fig. 4**), and the resulting ensemble averages are very accurate. Appropriate fits can be obtained by omitting these spikes from the fitted data.

The time reversed ensemble average generated from the actual locations of the simulated binding interactions led to an overestimate of the magnitude of substep 1 (**Fig. 5B-C**; **Table 3**). To generate the time reversed ensemble average, short-lived binding interactions are extended in time to match the duration of the longest-lived binding interaction, and the value of this extension equals the average position of the beads during the first 5 ms of the binding interaction. The rate of transitioning from the first substep to the second substep in our simulated data was 70 s_-1_, matching the rate of ADP release for beta cardiac myosin (22). Because of this fast rate, a large number of transitions to the second substep occur before the 5 ms used to generate the extensions, leading to inaccurate extension values. The proportion of binding interactions which are expected to transition to the second substep within the first 5 ms is given by the integral of the probability density function:

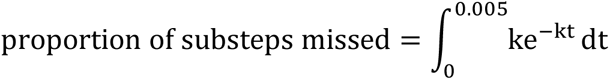

For a rate of 70 s_-1_, this proportion is equal to about 30%, and this will lead to an overestimate of the size of the first substep. A possible fix is to shorten the 5 ms window used for calculating the extensions, but it then becomes crucial that the binding initiation times are determined with high accuracy. Neither the single threshold method nor the peak-to-peak method have sufficient resolution to accurately determine the exact initiation times (**Fig. 4**). Even the change point algorithm, which we have shown to have an average error of about 0.5 ms, would be insufficient for generating the time reversed ensemble averages of interactions with very fast kinetics. It is possible that this could be improved with faster data sampling. In the case of transitions with slower kinetics, this problem is easily avoided. When we simulated 1000 binding interactions using much slower rates (k_f_ of 5 s_-1_ and k_r_ of 2 s_-1_, sets 11-20), we were able to generate time forward and time reversed ensemble averages with accurate step sizes using multiple methods (**Fig. S4**).

### Performance of the computational tool to analyze experimental data

To test the ability of the computational tool on real experimental data, we conducted optical trapping experiments using ventricular myosin at 1 µM ATP (**Fig. 6**). We intentionally collected a small data set consisting of 66 binding interactions from 5 molecules. Binding interactions were identified using the peak-to-peak method, and transition points were identified using the change point algorithm. The SPASM computational tool was used to generate cumulative distributions of individual binding interactions (**Fig. 6B**). The cumulative distributions of the attachment durations is well fit by a single exponential function. This exponential rate gives the rate of actomyosin detachment, and it has a value of 4.7 s_-1_, which is consistent with the expected rate of ATP-induced actomyosin dissociation at 1 µM ATP (22). The cumulative distribution of total working stroke displacements is well fit by a single normal distribution (indicating likely single molecule conditions), with a mean of displacement of 6.3 nm and a standard deviation of 9.2 nm. This is consistent with previous measurements of the cardiac myosin working stroke (7, 21). Ensemble averages (**Fig. 6C**) reveal that, consistent with previous measurements (7, 21), ventricular cardiac myosin has a two-substep working stroke with a first substep of 4.4 nm and a total displacement of 6.4 nm. The time forward averages have a rate of 74 s_-1_, which is consistent with the rate of ADP release, and the time reversed averages have a rate of 3.2 s_-1_, which is consistent with the rate of ATP-induced actomyosin dissociation at 1 µM ATP (22). Taken together, our computational tool can generate accurate ensemble averages with sharp transitions from a relatively small set of experimental data.

**Figure 6.**
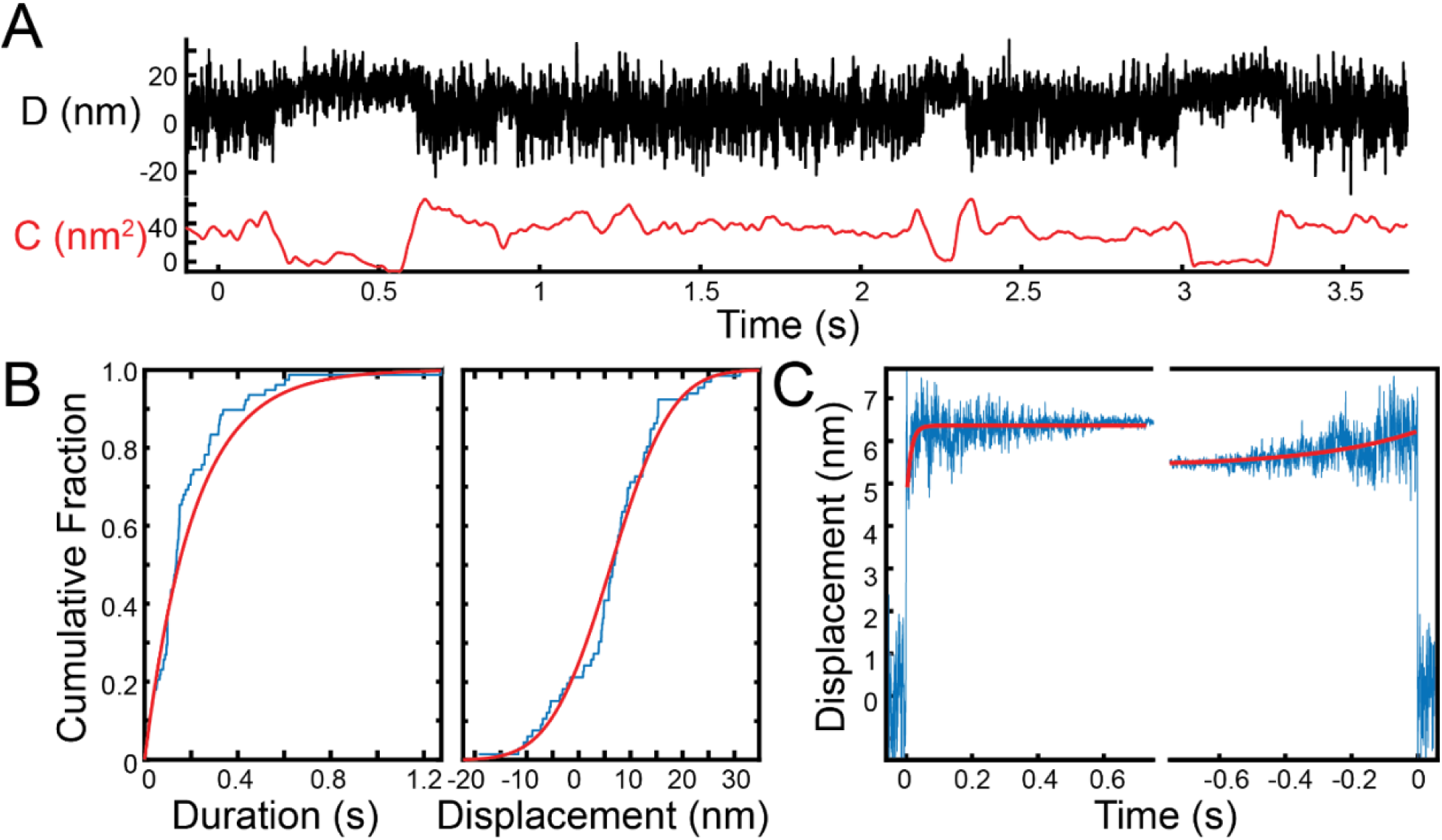
Ensemble averages of experimental optical trapping data. The kinetics and mechanics of cardiac myosin in 1 µM ATP were measured using the three-bead assay. (**A**) Experimental data trace showing the displacement (D) and covariance (C). (**B**) Cumulative distributions for the (left) binding interaction durations and (right) total working stroke displacements. The peak-to-peak method was used to detect binding interactions. Red lines show the cumulative fits based on (left) exponential and (right) normal distributions. The characteristic rate obtained from the fit to the distribution of attachment durations gives a detachment rate equal to 4.7 s_-1_, which is consistent with the expected rate of ATP-induced actomyosin dissociation at 1 µM ATP. The distribution of total step sizes has a mean of 6.3 nm and a standard deviation of 9.2 nm. (**C**) The change point algorithm was used to align the interactions identified using the peak-to-peak method. A total of 66 binding interactions from 5 molecules were analyzed. The resulting ensemble averages have estimated substep sizes of 4.4 nm and 2.0 nm. The estimated time forward rate is 74 s_-1_, and the estimated time reversed rate is 3.2 s_-1_. These values are consistent with previous measurements using a much larger data set, and they agree well with the previously measured rates of ADP release and ATP-induced dissociation 1 µM ATP.

### Broader applicability of the approach

The methods presented in this paper were applied to study actomyosin. As noted previously, the three-bead assay has been used to explore many different single-molecule systems, including dynein, the lac repressor, and kinesins. Moreover, the general ideas behind our computational tool are broadly applicable to any set of data containing well-defined populations which can be distinguished through some aspect of the data. One such possibility is data obtained from single-molecule FRET experiments. In the Supporting Materials, we describe how to adapt the change point algorithm to systems where the desired change points occur in data with different distributions.

### Limitations

There are a number of limitations accompanying our computational tool and the methods we use to analyze our data. While the covariance between the position of each trapped bead in the three-bead assay is very helpful for locating binding interactions under many circumstances, it does have drawbacks. The covariance is calculated over a window, and therefore it does not always drop enough during short-lived binding interactions to register as a genuine binding interaction. Furthermore, depending on the quality of the data, it may be difficult or even impossible to obtain a covariance histogram with two distinct populations. This could stem from system compliances. One benefit of the peak-to-peak method is that the covariance histogram populations do not need to be completely separated to avoid false positive binding interactions, but a certain degree of separation is needed to make the covariance useful. Additionally, analysis is dependent on many parameters, including the window sizes used to calculate and smooth the covariance, and it can be difficult to choose appropriate values for these parameters for a given experimental system. The computational tool includes features which allow the user to correct for these drawbacks when they are encountered. Finally, as evidenced by the ensemble averages generated from our simulated data (**Fig. 5**), ensemble averaging has limitations for estimating the rates and substep sizes for transitions with very fast kinetics.

### Summary

Here, we developed a computational tool, SPASM, for the detection and alignment of single-molecule binding interactions and for the generation of ensemble averages which can reveal characteristics about the data that are often obscured by noise. We show that it can be advantageous to use separate techniques for the detection and alignment of binding interactions. Specifically, we show that the addition of a change point algorithm to identify transition times can generate precise ensemble averages with improved alignment. We offer the computational tool, with an intuitive graphical user interface, along with a user guide so that the reader can apply these methods to their own data.

## Supporting information

User Guide

## Acknowledgements

Funding for this project was provided by the National Institutes of Health (R01HL141086 to M.J.G., T32EB018266 to S.R.C.).

## Author contributions

T.B. wrote the computational tool, simulated data, and analyzed data. W.T.S. built the optical trap and wrote software for data acquisition. S.R.C. collected optical trapping data. M.J.G. wrote code for the simulator and analyzed data. T.B. and M.J.G. wrote the first draft of the paper, and all authors contributed to the final draft.

## Code availability

The SPASM computational tool can be found at: https://github.com/GreenbergLab/SPASM

This repository includes the open source code for SPASM (**SPASM.m**), compiled versions for Windows (**SPASM_Windows.exe**) and macOS (**SPASM_macOS.app.zip**), a versions of the program which analyze only one trapped bead and uses variance thresholds rather than covariance thresholds (**SPASM_one_bead.m, SPASM_one_bead_Windows.exe, SPASM_one_bead_macOS.app.zip**), MATLAB code to generate simulated data (**simulator.m**), a user guide for the aforementioned components (**User_Guide.pdf**), and the simulated data sets analyzed in this paper (**sets 1-20**).

