## Supporting Materials for "Computational Tool for Ensemble Averaging of Single-Molecule Data"

### Derivation of the log-likelihood function for the change point algorithm

Generally, given a set of data, a change point algorithm can help determine whether any changes have occurred and, if so, where the changes most likely occurred. In our implementation, we first use the covariance to identify binding interactions and then apply the change point algorithm to precisely determine where the transitions between the bound and unbound states occurred. For each identified binding interaction, we consider windows of data surrounding the interactions. Therefore, we assume each window contains exactly two change points, and we only need to determine where they occur.

The change point algorithm aims to maximize the log-likelihood function, whose derivation requires knowledge of the distributions underlying a particular set of data. In the case of optical trapping data, the data points are typically normally distributed with unique means and variances for the bound and unbound states. Other types of data sets might have different underlying distributions and therefore different log-likelihood functions that must be maximized. For example, in the case of single molecule FRET data, the distributions of FRET efficiencies in each state typically have unique means but do not necessarily have unique variances. In the case of single photon wait times, the underlying distributions are Poisson distributed rather than normally distributed. Below, we show how to derive the log-likelihood function for normally distributed data with a change in both mean and variance, using maximum likelihood estimation. For other types of data, the log-likelihood function can be derived using similar methods.

For a window of data occurring at times $\boldsymbol{T} = \{1,2,\dots,N\}$, we know the average position of the beads $\boldsymbol{X} = \{X_1, X_2, \dots, X_N\}$. We assume there exist times $i$ and $j$ within $\{1,2,\dots,N-1\}$ such that $\{X_1, X_2, \dots, X_i\}$ and $\{X_{j+1}, X_{j+2}, \dots, X_N\}$ are drawn from some normal distribution $N(\mu_U, {\sigma_U}^2)$ and $\{X_{i+1}, X_{i+2}, \dots, X_j\}$ are drawn from a second normal distribution $N(\mu_B, {\sigma_B}^2)$. Therefore, the density function for any point $X_k$ within our data is:

$$f(X_k) = \begin{cases} \frac{1}{\sqrt{2\pi{\sigma_U}^2}} \exp\left[-\frac{1}{2}\left(\frac{X_k - \mu_U}{\sigma_U}\right)^2\right], & \text{if } 1 \le k \le i \text{ or } j < k \le N \\ \frac{1}{\sqrt{2\pi{\sigma_B}^2}} \exp\left[-\frac{1}{2}\left(\frac{X_k - \mu_B}{\sigma_B}\right)^2\right], & \text{if } i < k \le j \end{cases}$$

Assuming independence among the datapoints, the probability of obtaining the entire data set $\boldsymbol{X}$ is given by the likelihood function:

$$f(\boldsymbol{X}) = f(X_1)f(X_2)\dots f(X_N)$$

$$= \left(\prod_{k=1}^{i} \frac{1}{\sqrt{2\pi{\sigma_U}^2}} \exp\left[-\frac{1}{2}\left(\frac{X_k - \mu_U}{\sigma_U}\right)^2\right]\right) * \left(\prod_{k=i+1}^{j} \frac{1}{\sqrt{2\pi{\sigma_B}^2}} \exp\left[-\frac{1}{2}\left(\frac{X_k - \mu_B}{\sigma_B}\right)^2\right]\right)$$

$$* \left(\prod_{k=j+1}^{N} \frac{1}{\sqrt{2\pi{\sigma_U}^2}} \exp\left[-\frac{1}{2}\left(\frac{X_k - \mu_U}{\sigma_U}\right)^2\right]\right)$$

Only the values $\{X_1, X_2, \dots, X_N\}$ are known. We wish to determine values for the parameters $\mu_U$, ${\sigma_U}^2$, $\mu_B$, ${\sigma_B}^2$, $i$, and $j$ which maximize $f$. Equivalently, one could maximize the log-likelihood function. The log-likelihood function is the natural logarithm of $f$:

$$\ln f = \left[-\frac{i}{2}\ln(2\pi{\sigma_U}^2) - \frac{1}{2{\sigma_U}^2}\sum_{k=1}^{i}(X_k - \mu_U)^2\right]$$

$$+ \left[-\frac{j-i}{2}\ln(2\pi{\sigma_B}^2) - \frac{1}{2{\sigma_B}^2}\sum_{k=i+1}^{j}(X_k - \mu_B)^2\right]$$

$$+ \left[-\frac{N-j}{2}\ln(2\pi{\sigma_U}^2) - \frac{1}{2{\sigma_U}^2}\sum_{k=j+1}^{N}(X_k - \mu_U)^2\right]$$

An estimate for $\widehat{\mu_U}$, the value of $\mu_U$ which maximizes $\ln f$, is obtained by solving $\frac{\partial \ln f}{\partial \mu_U} = 0$:

$$\frac{\partial \ln f}{\partial \mu_U} = \frac{1}{{\sigma_U}^2}\left[\sum_{k=1}^{i}(X_k - \mu_U) + \sum_{k=j+1}^{N}(X_k - \mu_U)\right]$$

$$= \frac{1}{{\sigma_U}^2}\left[\left(\sum_{k=1}^{i}X_k + \sum_{k=j+1}^{N}X_k\right) - (N - j + i)\mu_U\right] = 0$$

$$\Rightarrow \sum_{k=1}^{i}X_k + \sum_{k=j+1}^{N}X_k = (N - j + i)\mu_U$$

$$\Rightarrow \widehat{\mu_U} = \frac{1}{(N - j + i)}\left[\sum_{k=1}^{i}X_k + \sum_{k=j+1}^{N}X_k\right]$$

The estimate for $\widehat{\mu_B}$ is found similarly to be:

$$\widehat{\mu_B} = \frac{1}{j - i}\sum_{k=i+1}^{j}X_k$$

An estimate for $\widehat{{\sigma_U}^2}$ is obtained by solving $\frac{\partial \ln f}{\partial {\sigma_U}^2} = 0$ and using $\widehat{\mu_U}$ as an estimate for $\mu_U$:

$$\frac{\partial \ln f}{\partial {\sigma_U}^2} = -\frac{N - j + i}{2{\sigma_U}^2} + \frac{1}{2{\sigma_U}^4}\left[\sum_{k=1}^{i}(X_k - \mu_U)^2 + \sum_{k=j+1}^{N}(X_k - \mu_U)^2\right] = 0$$

$$\Rightarrow \frac{N - j + i}{2{\sigma_U}^2} = \frac{1}{2{\sigma_U}^4}\left[\sum_{k=1}^{i}(X_k - \mu_U)^2 + \sum_{k=j+1}^{N}(X_k - \mu_U)^2\right]$$

$$\Rightarrow N - j + i = \frac{1}{{\sigma_U}^2}\left[\sum_{k=1}^{i}(X_k - \mu_U)^2 + \sum_{k=j+1}^{N}(X_k - \mu_U)^2\right]$$

$$\Rightarrow \widehat{{\sigma_U}^2} = \frac{1}{N - j + i}\left[\sum_{k=1}^{i}(X_k - \widehat{\mu_U})^2 + \sum_{k=j+1}^{N}(X_k - \widehat{\mu_U})^2\right]$$

The estimate for $\widehat{{\sigma_B}^2}$ is similarly given by:

$$\widehat{{\sigma_B}^2} = \frac{1}{j - i}\sum_{k=i+1}^{j}(X_k - \widehat{\mu_B})^2$$

The log-likelihood function simplifies after these values are substituted:

$$\ln f = \left[-\frac{j - i}{2}\ln\left(2\pi\widehat{{\sigma_B}^2}\right) - \frac{N - j + i}{2}\ln\left(2\pi\widehat{{\sigma_U}^2}\right)\right] - \frac{j - i}{2} - \frac{N - j + i}{2}$$

$$= \left[ -\frac{j-i}{2} \ln\left(\widehat{\sigma_B{}^2}\right) - \frac{N-j+i}{2} \ln\left(\widehat{\sigma_U{}^2}\right) \right] - \frac{N}{2} \ln(2\pi) - \frac{N}{2}$$

It is sufficient to maximize the following function:

$$L = -\frac{j-i}{2} \ln\left(\widehat{\sigma_B{}^2}\right) - \frac{N-j+i}{2} \ln\left(\widehat{\sigma_U{}^2}\right)$$

This function depends only on $\boldsymbol{X}$, $i$, and $j$. As $i$ and $j$ must be within the set $\{1,2,\dots,N-1\}$, $\hat{\imath}$ and $\hat{\jmath}$ may be determined empirically.

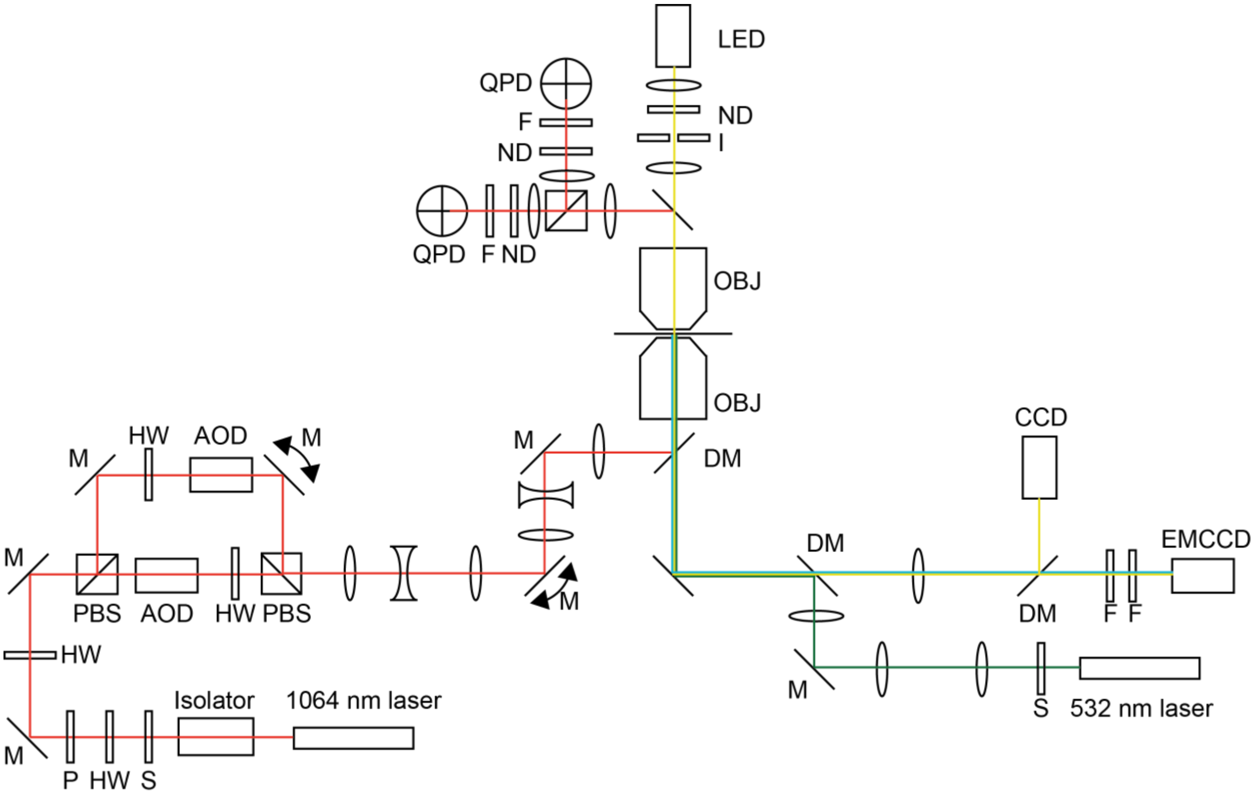


**Figure S1. Optical trap layout.** A half-wave plate (HW) / polarizer (P) combination attenuates the vertically polarized 1064 nm trapping laser beam (IPG) to the intensity required for the experiment, while a second half-wave plate adjusts the polarization angle to 45 degrees such that two traps with orthogonal polarization and equal power can be formed. The beam is split into vertical and horizontal components by the first polarizing beam splitter (PBS, Newport), and each beam passes through an acousto-optic deflector (AOD, Gooch and Housego) for computer-controlled trap steering and power control. The AODs require an input polarization parallel to their horizontal steering plane, and the output from the AODs is vertically polarized. Therefore, a half-wave plate is needed to rotate the polarization before one AOD, and a separate half-wave plate is used to rotate polarization after the other AOD. The two polarization separated beams are then recombined using a second PBS. The beams are then expanded to fill the back aperture of the microscope objective by a pair of beam expanders, each formed by two planoconvex lenses with a biconcave lens at the crossover point to limit fluctuations caused by localized heating of the air. Motorized mirrors (M, with arrows, Newport) are placed at two points for coarse positioning of the beams: the first adjusts one trap's position while the second, which is conjugate to the back aperture of the objective, repositions both traps simultaneously in the microscope field of view. The AOD's and quadrant photodiode detectors (QPDs, see below) are conjugate to the back focal plane of the objective so that beam steering is not registered as signal change on the QPDs. After the laser beams pass through the trapped beads, the resulting interference patterns are collected by the condenser (a second objective lens) and split into their respective polarizations by a PBS. These polarization-separated interference patterns are then imaged onto the two quadrant photodiodes (QPDs, First Sensor). Neutral density (ND) and 1064 nm bandpass filters (F) are used to avoid detector saturation and to remove non-trapping wavelengths of light, respectively.

Beads are visualized on an electron multiplying charge coupled device (EMCCD), or alternatively a standard CCD camera using wide field transillumination provided by a 730 nm LED (Thorlabs). An iris (I) is positioned so that its image is in focus when the condenser is set at the correct height for trapping beam imaging onto the QPDs. Before reaching the condenser, this beam passes through a dichroic mirror (DM, Semrock) that is used to reflect the trapping beams to the QPDs.

The EMCCD is also used to visualize fluorescent actin molecules in a wide field epifluorescence arrangement using an expanded 532 nm laser (CrystaLaser) introduced into the trapping beam path with a dichroic mirror. Another dichroic mirror separates the excitation light from the emission signal, which is subsequently filtered by dual filters (F). Both the 532 nm and 1064 nm lasers have shutters (S).

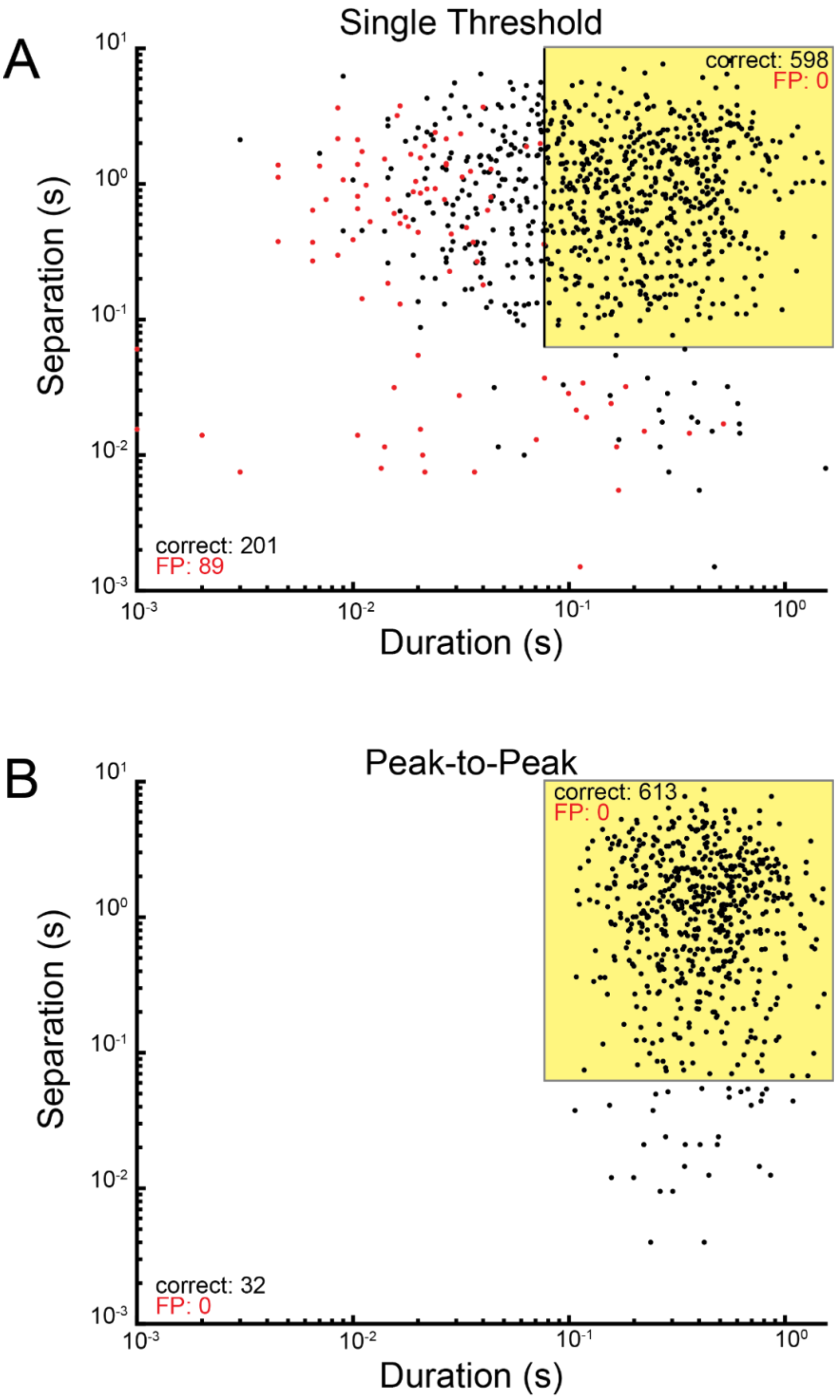


**Figure S2. The single threshold method detects more false positive binding interactions than the peak-to-peak method, and efficiently excluding these**

**interactions is difficult.** 10 sets of data, each containing 100 simulated binding interactions, were analyzed by either the single threshold method or the peak-to-peak method. (**A**) Scatter plot showing the duration and the smaller of the two separations of each binding interaction detected by the single threshold method, both in units of seconds. A binding interaction's two separations are the amounts of time separating that interaction from the preceding and subsequent interactions. Both axes are scaled logarithmically. False positive interactions identified by the method are shown in red. As can be seen, most false positive interactions have relatively short durations and/or separations. The yellow box indicates the binding interactions which remain after filtering out any interactions with a duration shorter than 77 ms or a separation shorter than 63 ms. These two values were chosen to be as small as possible while still filtering out all false positive interactions. Due to significant overlap between the false positive interactions and many of the correctly detected interactions, it is not easy to pick suitable values for filtering the interactions unless the false positive binding interactions have already been identified. This can only confidently be done using simulated data where one can know for certain whether a detected interaction is a false positive. After the filtering takes place, 598 interactions remain. (**B**) Scatter plot showing the duration and the smaller of the two separations of each binding interaction detected by the peak-to-peak method. No false positive interactions were detected by the peak-to-peak method. Applying the same filtering as with the single threshold method leaves 613 interactions. Note that the durations and separations are calculated based on the binding initiation and termination times estimated by the single threshold or peak-to-peak method. A binding interaction detected by the peak-to-peak method will always appear to be longer than the corresponding interaction detected by the single threshold method (see main text for details).

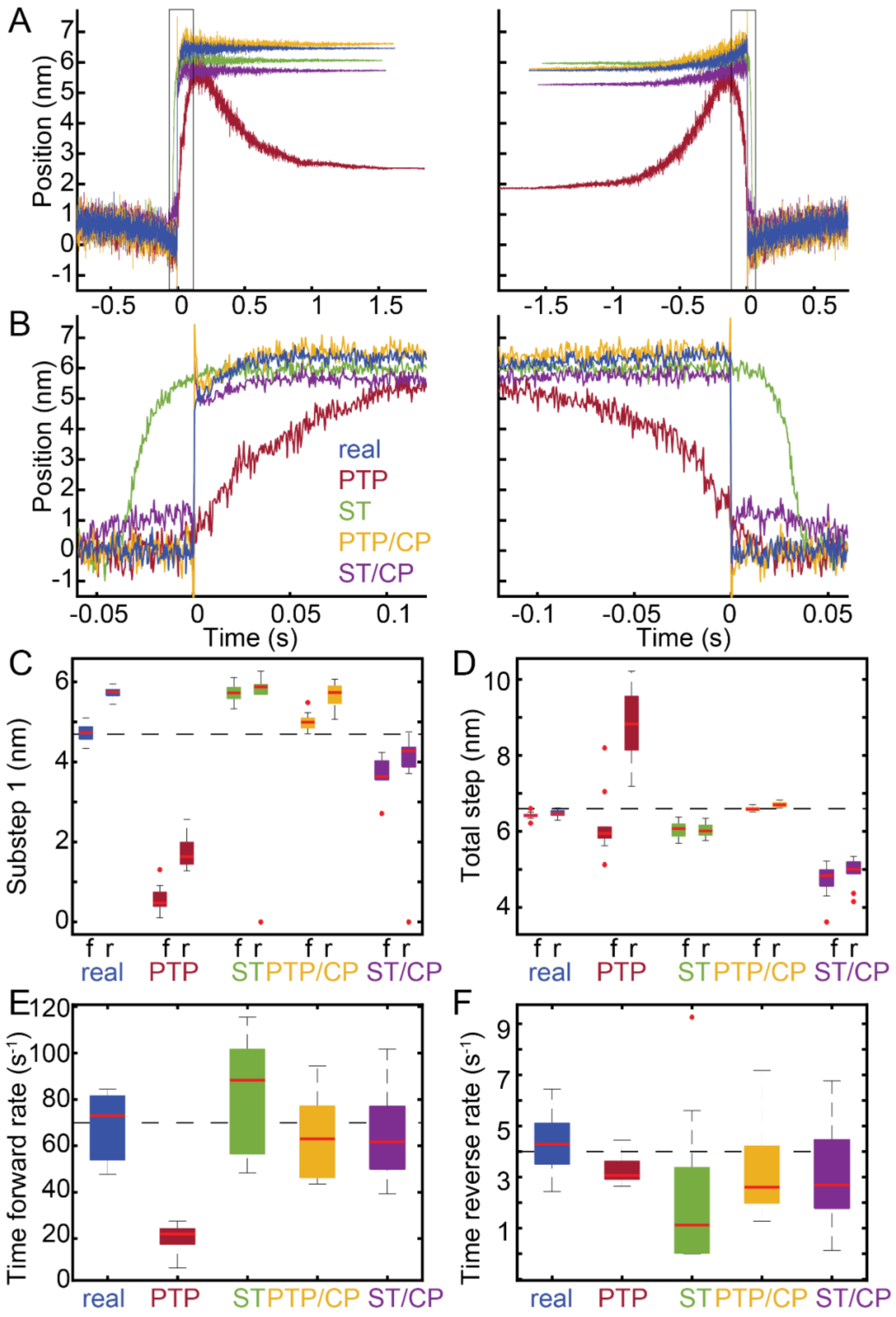


**Figure S3. Ensemble averages generated with the single threshold method without filtering events leads to an underestimate of the total step size due to the inclusion**

**of false positive binding interactions.** The same sets of simulated data were analyzed as with **Fig. 5** (sets 1-10), each containing 100 binding interactions. As with **Fig. 5**, interactions were detected using either the peak-to-peak (PTP) or the single threshold (ST) method, and interactions were aligned using either the transitions estimated by the covariance threshold method or the change points identified by the change point algorithm (CP). Here, unlike with **Fig. 5**, none of the binding interactions detected by the single threshold method were removed. Many of these binding interactions are false positives (see Figure S2). (**A-B**) Time forward (left) and time reversed (right) ensemble averages were generated from the known locations of the actual simulated binding interactions (real) and from each method of analysis. (**C-F**) Within each of the 10 sets of data, ensemble averages were generated and fit with single exponential functions. The substep sizes and rates of the simulated myosin working stroke were estimated from the exponential fits. Box plots show the estimated parameters for each analysis method. Outliers are indicated by red dots. The substep sizes were estimated from both the time forward (f) and time reversed (r) ensemble averages. Horizontal dashed lines show the values of the simulated parameters. The total step size is underestimated by the single threshold method, both with (ST/CP) and without (ST) the change point algorithm, due to the inclusion of false positive binding interactions which do not generate any net displacement in the optical trap.

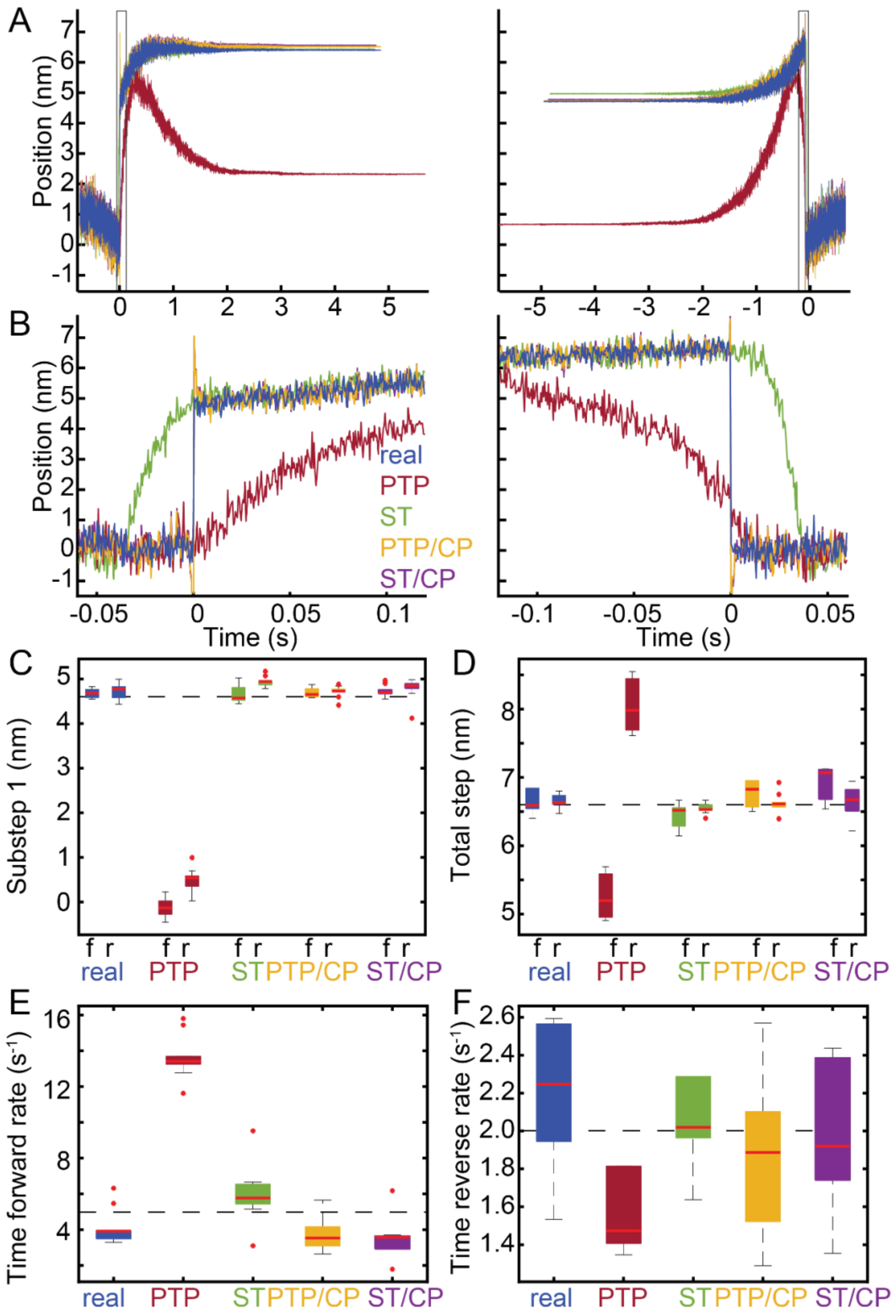


**Figure S4. Ensemble averages are able to accurately estimate the substep sizes and kinetic rates when the underlying transitions have slower kinetics.** 10 sets of data were simulated, each containing 100 binding interactions (sets 11-20). Here, the rate of transitioning from the first to second substep was set to 5 $s^{-1}$, which is much lower than the rate of 70 $s^{-1}$ used in **Fig. 5**. Additionally, the rate of transitioning from the second substep to the detached state was set to 2 $s^{-1}$. The magnitude of the two substeps were still 4.7 nm and 1.9 nm, as before. As in **Fig. 5**, interactions were detected using either the peak-to-peak (PTP) or the single threshold (ST) method, and interactions were aligned using either the transitions estimated by the covariance threshold method or the change points identified by the change point algorithm (CP). Also similar to **Fig. 5**, binding interactions detected by the single threshold method which were too short or too close to other interactions were excluded from analysis, to minimize the number of false positive interactions. (**A-B**) Time forward (left) and time reversed (right) ensemble averages were generated from the known locations of the actual simulated binding interactions (real) and from each method of analysis. (**C-F**) Within each of the 10 sets of data, ensemble averages were generated and fit with single exponential functions. The substep sizes and rates of the simulated myosin working stroke were estimated from the exponential fits. Box plots show the estimated parameters for each analysis method. Outliers are indicated by red dots. The substep sizes were estimated from both the time forward (f) and time reversed (r) ensemble averages. Horizontal dashed lines show the values of the simulated parameters. Unlike with **Fig. 5**, the time reversed averages generated from simulations with slower kinetics offer accurate estimates of the size of substep 1.
