## Supplementary material for "Computational Tool for Ensemble Averaging of Single-Molecule Data": User Guide

### SPASM User Guide

#### Overview

This is a user guide for **SPASM** (Software for Precise Analysis of Single Molecules), a MATLAB program to identify and analyze actomyosin binding interactions based on data obtained from the three-bead optical trapping assay.

The program can be found at https://github.com/GreenbergLab/SPASM.

The methods of data analysis are described in a paper published at XX.

We have provided the uncompiled code for those who have MATLAB installed on their computer. **SPASM** is compatible with MATLAB releases R2017b through at least R2020a.

- The Signal Processing Toolbox is only required for filtering the covariance (Section 3). The covariance will remain unfiltered if the Signal Processing Toolbox is not installed.
- The Optimization Toolbox is only required for fitting ensemble averages (Section 6). **SPASM** will not produce these fits if the Optimization Toolbox is not installed.

We have also provided compiled versions of **SPASM** which only require the MATLAB Compiler Runtime (MCR). The MCR will be installed on your computer upon running the installer. An internet connection is required for the installer to download the MCR.

To run the uncompiled version of **SPASM** in MATLAB, make sure the file **SPASM.m** is contained in your Current Folder or on your MATLAB path. You may need to move **SPASM.m** to your Current Folder or change your Current Folder to whatever folder contains **SPASM.m**. Then, either type *SPASM* in the Command Window and press Enter

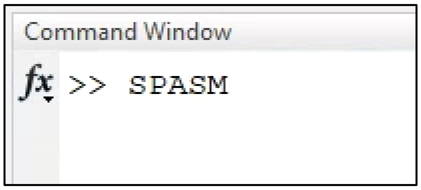

or open **SPASM.m** in the Editor and click the green Run button.

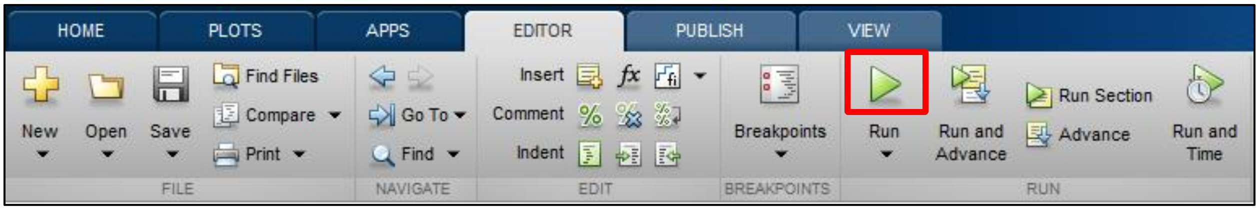

#### Analyze Single File

##### Section 1: Input Raw Data

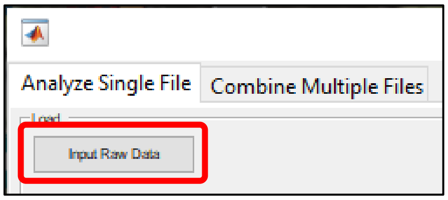

To load data, click the button labeled **Input Raw Data**. A window should appear, prompting you to select a .txt file. You must select a .txt file with the proper header and data format in order for **SPASM** to read the data correctly. Sample files with the proper format (sets 1-20) are included with **SPASM**. For a detailed description of this format, as well as a discussion of potential options for those whose data are formatted differently, see Appendix A.

Once a file is selected, the volt-to-nanometer conversion factor (CAL, units of nm/V) and stiffness (K, units of pN/nm) for each bead, the sampling frequency ($F_s$, units of Hz), and the name of the selected file will be displayed. Additionally, the positions of each bead will be plotted, with bead A shown in blue and bead B shown in orange.

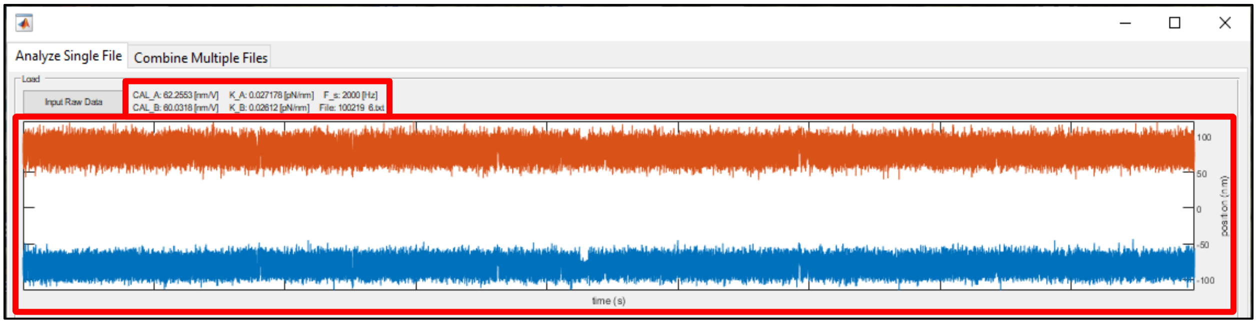

Right click anywhere within the plot to bring up a menu. The menu contains two buttons, **Save as .fig** and **Save as .eps**.

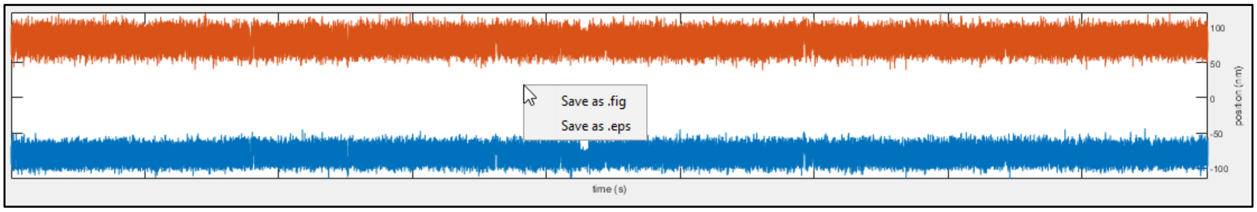

The **Save as .fig** button allows you to save the plot to a MATLAB .fig file. When you click **Save as .fig**, a window should appear, prompting you to choose a filename and location for this .fig file. Once you have saved the file, you may then open it separately in MATLAB, edit the plot however you would like, and export it as an image.

The **Save as .eps** button allows you to instead save the plot to a .eps file.

A similar menu appears when you right click any plot created by **SPASM**.

#### Section 2: Select Data for Removal (optional)

You might want to ignore some of your data - for example, if the optical table gets bumped or if pretension is lost during data collection. To do this, click **Select Data for Removal**.

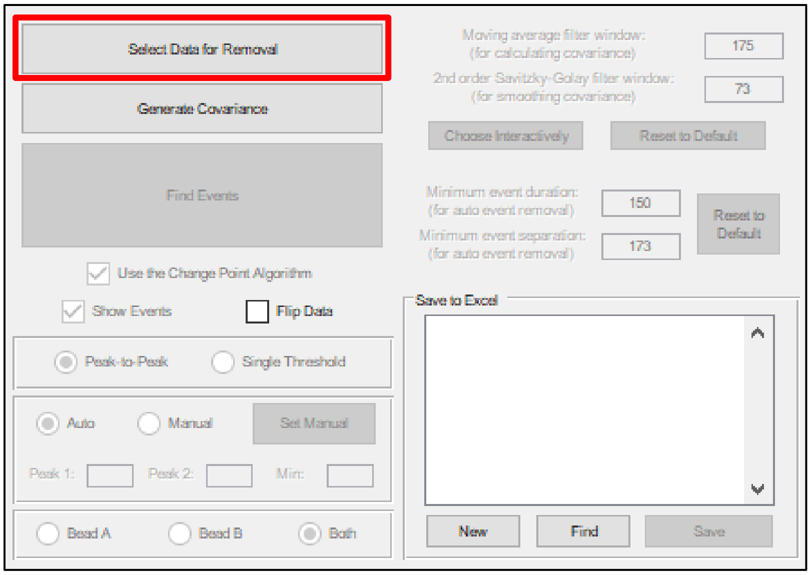

When this button is selected, a panel should appear beneath it. The panel contains three buttons, **Undo**, **Remove**, and **Save**, all of which will be disabled at first.

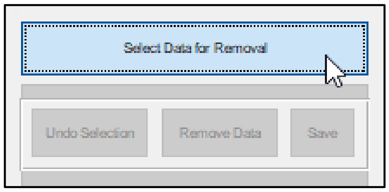

You may now click within the plot of the beads' positions to specify sections of data which should be removed. Pairs of clicks define the left and right boundaries of these sections. The first click will place a red vertical line at your cursor.

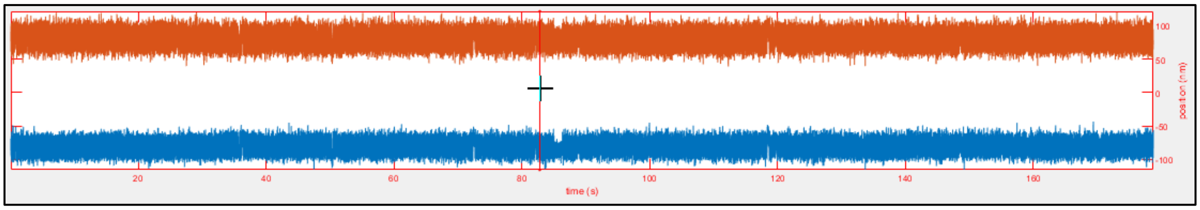

Click again to highlight a section of data in red:

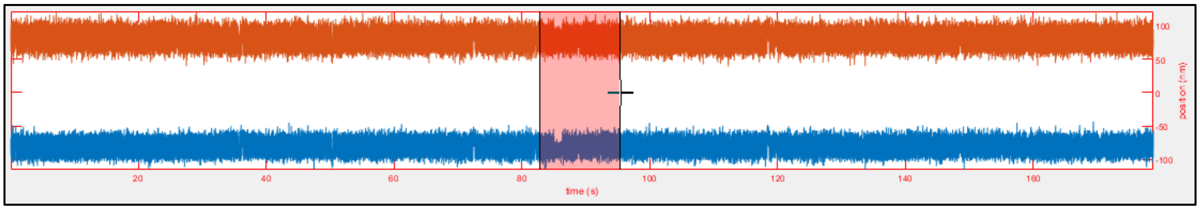

**a. Remove**

The **Remove** button becomes enabled after you have highlighted at least one section of data. By clicking **Remove**, all of the highlighted data will be removed. **SPASM** will then reset to the end of Section 1 of this user guide.

**b. Undo**

If you accidentally highlight a section of data incorrectly, you can undo it. Click on the section you would like to undo, and its color will change from red to blue.

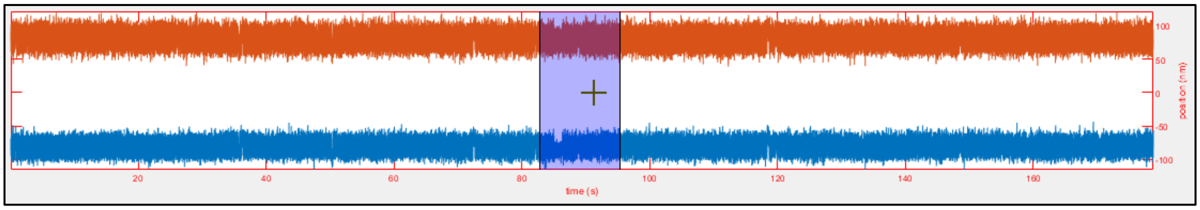

The **Undo** button becomes enabled once you have clicked a section. By clicking **Undo**, the selected patch will disappear. If you instead click again on the blue section, its color will return to red, indicating that the underlying data are once again targeted for removal.

**c. Save**

The **Save** button becomes enabled after any data has been removed. By clicking **Save**, you are able to save the newly trimmed data to a .txt file with either the same name or a new name. A window should appear, prompting you to choose a filename and location for this new file.

Click **Select Data for Removal** again to hide these three buttons and return to the main program.

#### Section 3: Generate Covariance

Our data analysis uses the covariance between the beads to detect the presence of binding events. If you choose to use the peak-to-peak method, the two peaks of the covariance histogram – or the minimum between the peaks if you decide to use the single threshold method – are used as thresholds which the covariance must cross to signal transitions between the attached and detached states. A typical workflow at this point, to prepare **SPASM** for event detection, involves

(1) deciding which method of data analysis to use (Section 3a),
(2) calculating the covariance and changing the window sizes until the covariance histogram has two distinct peaks (Section 3b), and
(3) verifying that the peaks or minimum of the histogram, depending on which method of data analysis you chose, are appropriate (Section 3c).

To calculate and display the covariance between the two beads, click **Generate Covariance**.

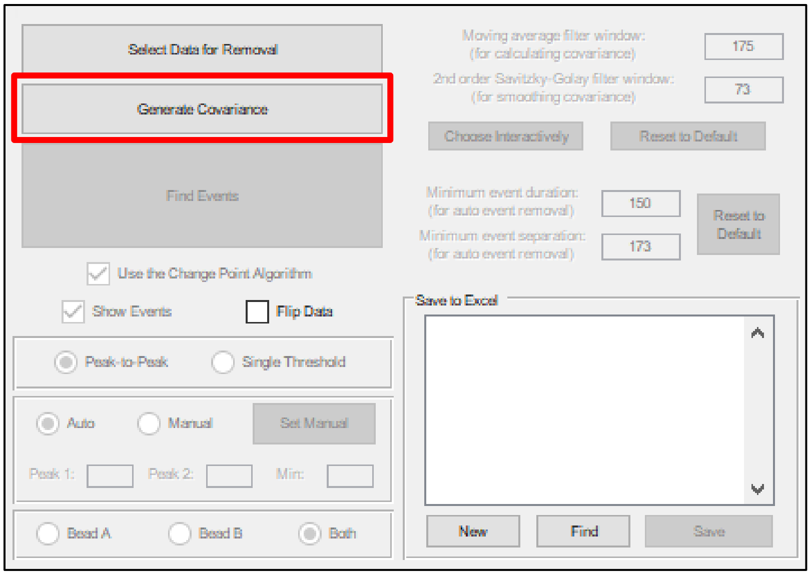

When you click this button, the covariance is plotted below the beads' positions.

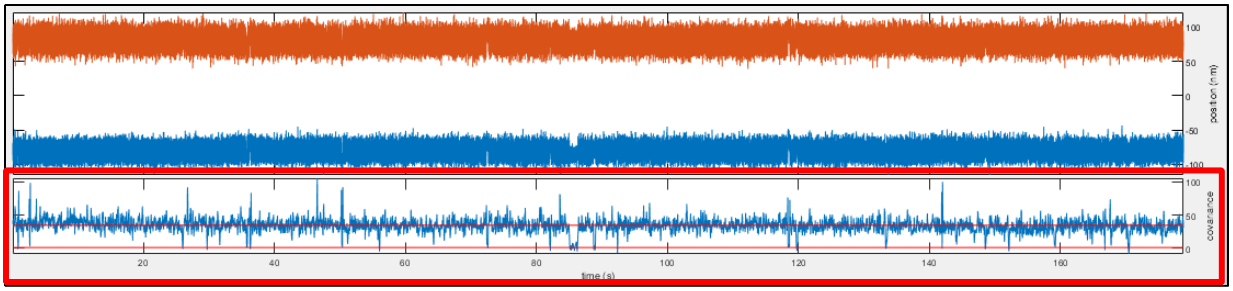

The covariance between the beads at time t is calculated according to the following equation:

$$\mathrm{Cov}_t(A, B) = E_{(w_c,t)}[A * B] - E_{(w_c,t)}[A] * E_{(w_c,t)}[B]$$

where A is the position of bead A, B is the position of bead B, and $E_{(w_c,t)}[X]$ denotes the mean of X over a window of size $w_c$ centered at t. Once the covariance has been calculated, it is filtered with a

2nd-order Savitzky-Golay filter, provided that the Signal Processing Toolbox is installed. The window sizes used in calculating and filtering the covariance are displayed on the right side of the user interface. Their values affect the covariance and therefore have a downstream effect on the data analysis. Section 3b explains how you can change their values.

A histogram of the covariance values is also plotted.

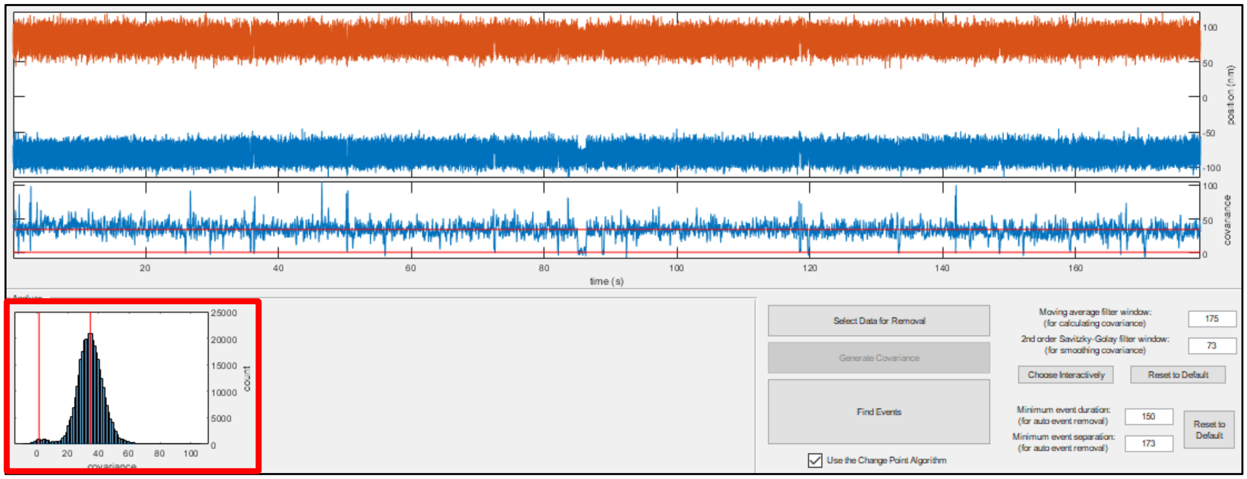

An ideal histogram will have two distinct peaks. The peak at higher covariance values corresponds to times during which actin and myosin are unbound. The peak at lower covariance values, near 0, corresponds to times during which actin and myosin are bound, as the covariance will drop upon myosin binding. If more than two peaks are found, this could be indicative of multiple bound molecules. If only one peak is found, this might indicate that there were no binding events or that there was insufficient pretension between the beads. It is important for accurate data analysis that the two peaks be sufficiently distinct, as their separation is used to detect binding events. If the histogram is suboptimal, then readjusting the aforementioned window sizes may improve its shape. Section 3b explains how to change the window sizes.

Red lines in both plots show **SPASM**'s best guess for the locations of the two peaks of the covariance histogram. These peaks are used in the peak-to-peak method to detect binding events. Section 3c explains how you can change their values.

##### Section 3a: Peak to Peak vs. Single Threshold (optional)

By default, **SPASM** will use the peak-to-peak method to detect binding events. In the peak-to-peak method, the covariance must extend from above the upper peak of the histogram to below the lower peak in order to signal an event start. As described in the paper, this criterion is stricter than using a single threshold, and it decreases the likelihood of detecting false positive events (that is, incorrectly detecting binding events during the detached state). However, you may wish to analyze your data using a single threshold. To do so, select the **Single Threshold** radio button.

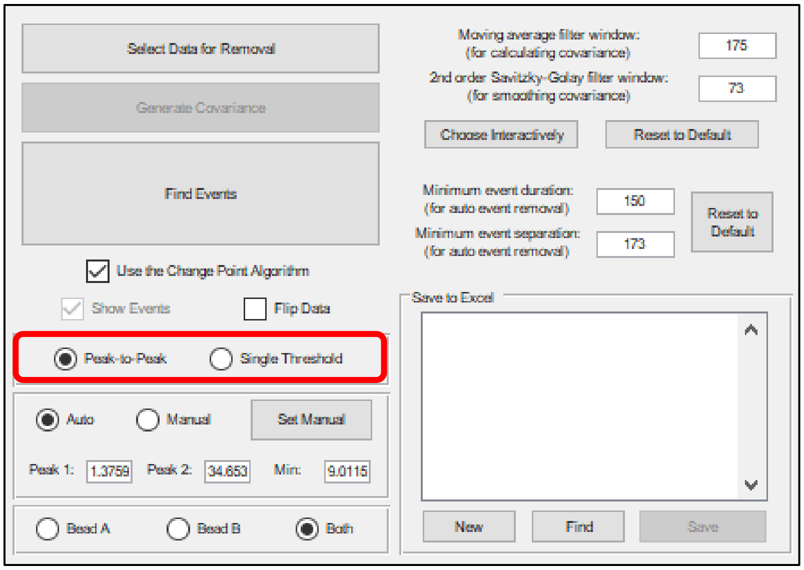

Once you select this button, the two red lines indicating the peaks of the covariance histogram will be replaced by a single red line indicating the new threshold location. By default, this location is determined by the minimum value of the histogram located between the two peaks. Section 3c explains how you can change this value.

#### Section 3b: Window Sizes Used to Generate the Covariance (optional)

As described at the beginning of Section 3, the covariance is calculated using a moving average and then smoothed using a 2nd order Savitzky-Golay filter. The default window sizes for these two filters are 175 points and 73 points, respectively; these values optimized **SPASM**'s performance on a specific set of simulated data. For an in-depth explanation of how these values were determined and guidelines for optimizing the window sizes against your own data, see Appendix C.

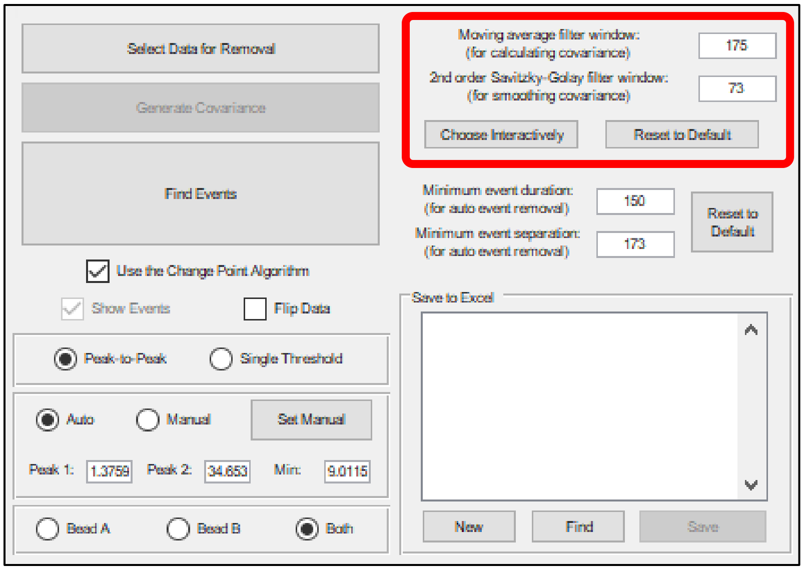

The default window sizes might not generate an appropriate covariance histogram that has two distinct peaks. There are two ways to change the window sizes from their default values.

##### i. Text Boxes

Enter a positive integer value for either filter window in the corresponding text box.

##### ii. Choose Interactively

Alternatively, click the **Choose Interactively** button. Doing so will bring up a set of axes, with the moving average filter window size along the X axis and the 2nd order Savitzky-Golay filter window size along the Y axis. A dot marks the current values of these two window sizes. By moving the cursor within these axes, the dot will relocate to the cursor's position, and the filter windows will change accordingly. The covariance and covariance histogram plots will update automatically, allowing you to visualize the effect of the new window sizes.

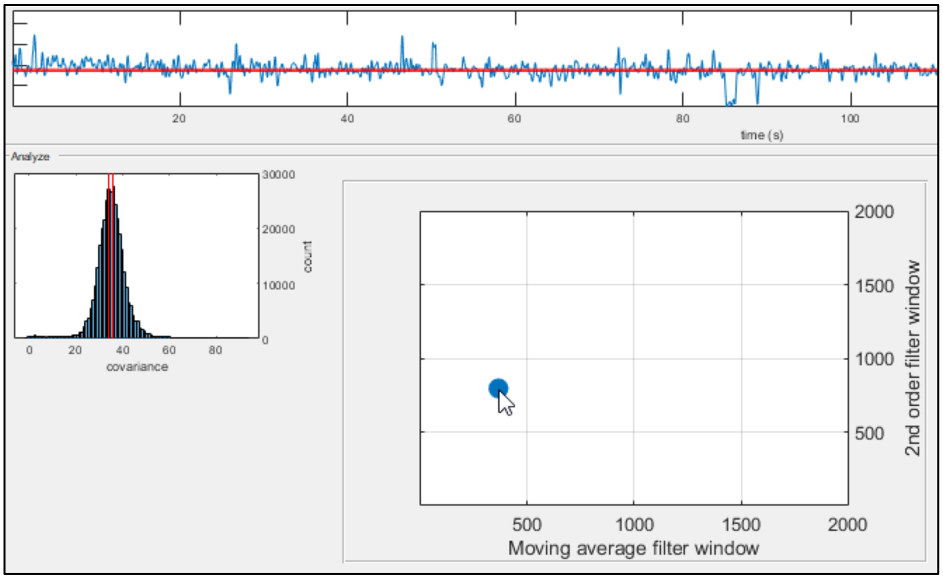

When the cursor leaves these axes, the filter windows return to the values they held before you clicked the **Choose Interactively** button.

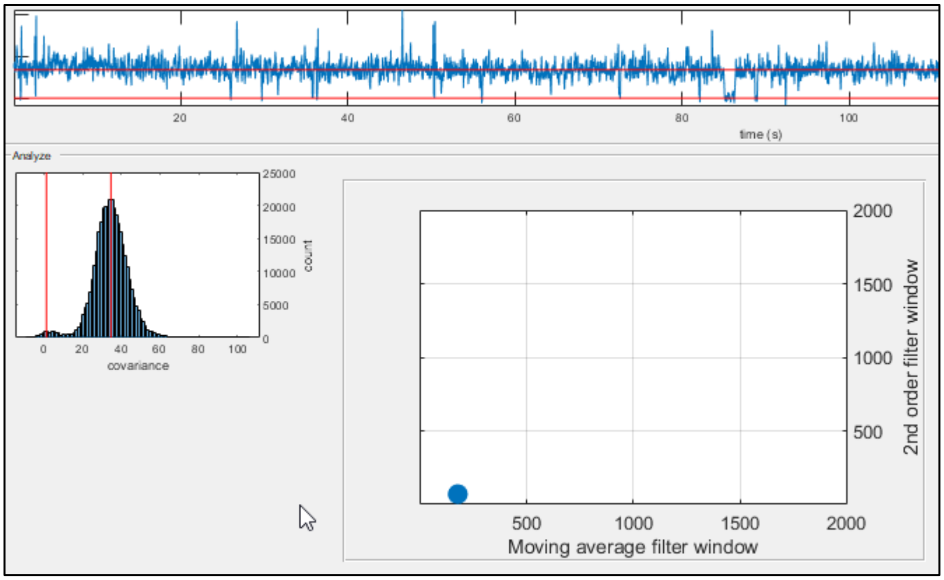

When you click the mouse, these axes will disappear. If you click within the axes, the window sizes will update to the new values.

When either window size changes, as the covariance and covariance histogram plots will update to reflect the change, the automatically calculated values for the peaks and minimum will also update. When you click the **Reset to Default** button, each window size will return to its default value.

#### Section 3c: Automatic vs Manual (optional)

You have the option to manually set the values of the histogram peaks and minimum.

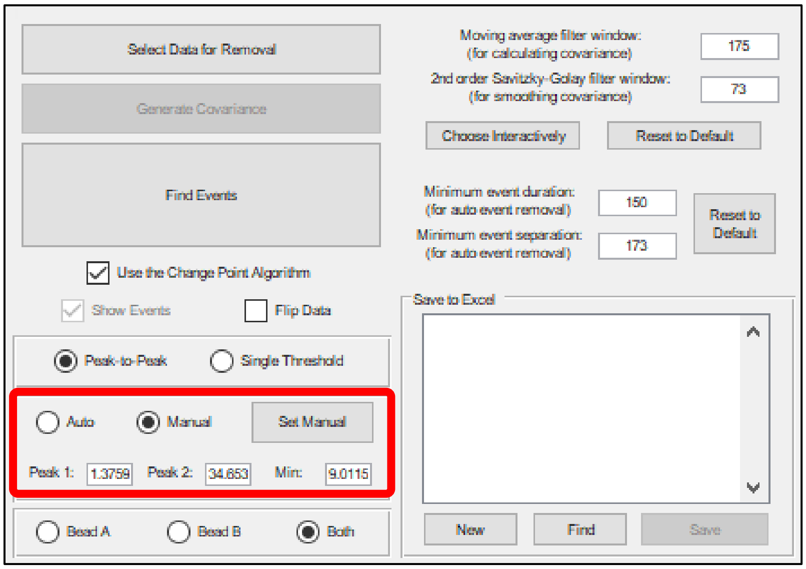

If you select the **Manual** radio button, **SPASM** will instead use the manually set values for the next steps in the analysis. By default, the manually set values will initially be equal to the automatically calculated values. There are two ways to change the manually set values:

##### i. Text Boxes

Enter numeric values for either the peak or minimum values in the corresponding text box located beneath the radio buttons.

##### ii. Set Manual

Alternatively, click the **Set Manual** button. After you do so, **SPASM** will wait for you to specify the desired values for the peaks and minimum by clicking on the covariance histogram. **SPASM** expects three clicks in total, one for each value. The order in which you set the values does not matter.

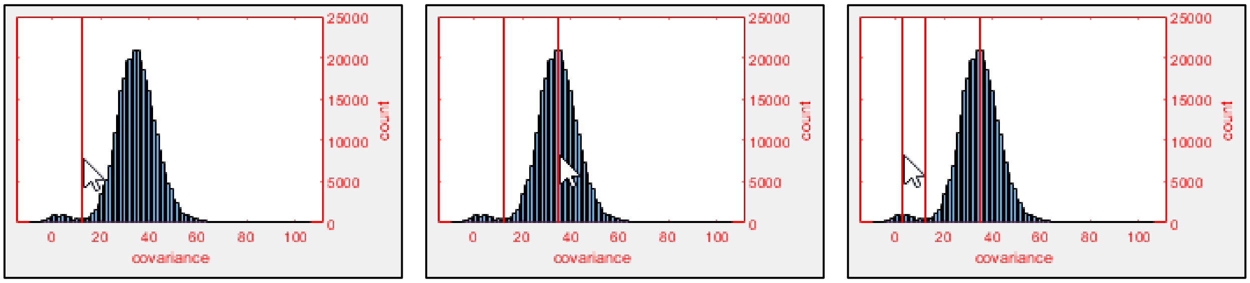

##### Section 3d: Select One Bead for Analysis (optional)

By default, **SPASM** averages the positions of both bead A and bead B together before running the change point algorithm or estimating substep sizes. However, this might not always be desirable. For example, when using an isometric optical clamp, forces are applied to one bead (the motor bead) so that the other bead (the transducer bead) is held in place. Such a clamp is useful for revealing information about the force dependence of actomyosin's attachment duration. With data obtained from an isometric optical clamp, the position of the transducer bead is necessary for calculating the covariance. However, any estimates of the force on the myosin should be derived from the position of the motor bead alone.

If you are analyzing such data, you have the option to specify which bead should be analyzed. To do so, select either the **Bead A** radio button or the **Bead B** radio button.

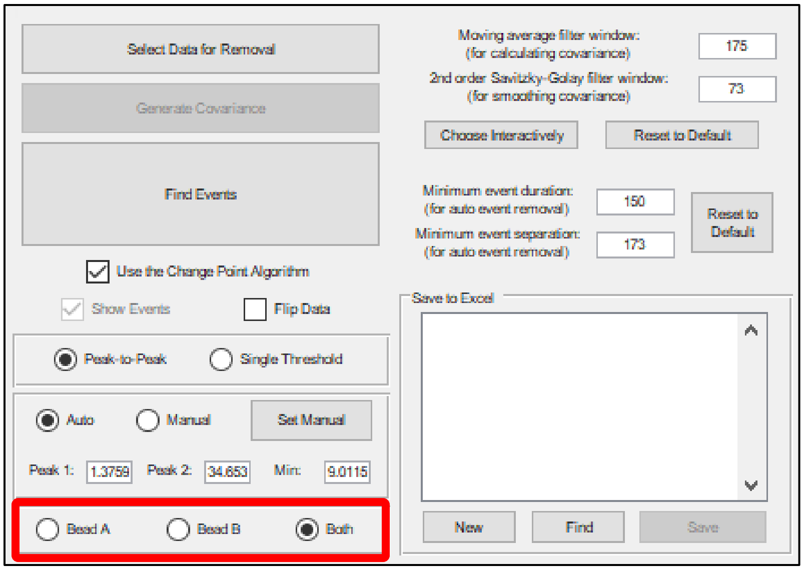

#### Section 3e: Bypass the Change Point Algorithm (optional)

You may choose to use the binding initiation and termination times estimated by the event detection method (Section 3a) rather than those estimated by the change point algorithm. To do so, uncheck the **Use the Change Point Algorithm** checkbox.

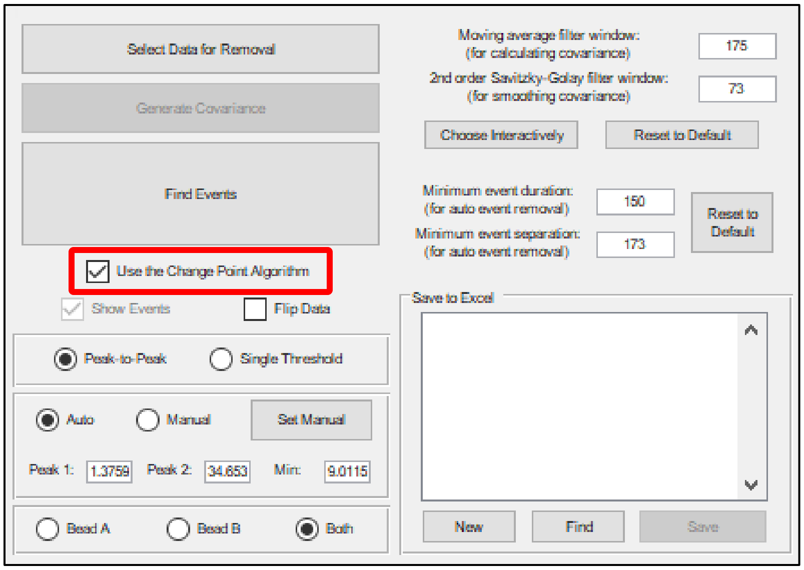

The change point algorithm is used by default.

#### Section 4: Find Events

Once the covariance and threshold(s) are established, **SPASM** is ready to find binding interactions (events). After events have been found (Section 4) and plots are generated (Sections 5, 6, and 7), a typical workflow involves

(1) viewing each event (Section 7a) and, if necessary, either
  (a) removing that event from data analysis (Sections 7b and 7c) or
  (b) correcting that event's start and end times (Section 7d), and
(2) saving the results of data analysis to an Excel workbook (Section 9).

To proceed with data analysis, click the **Find Events** button.

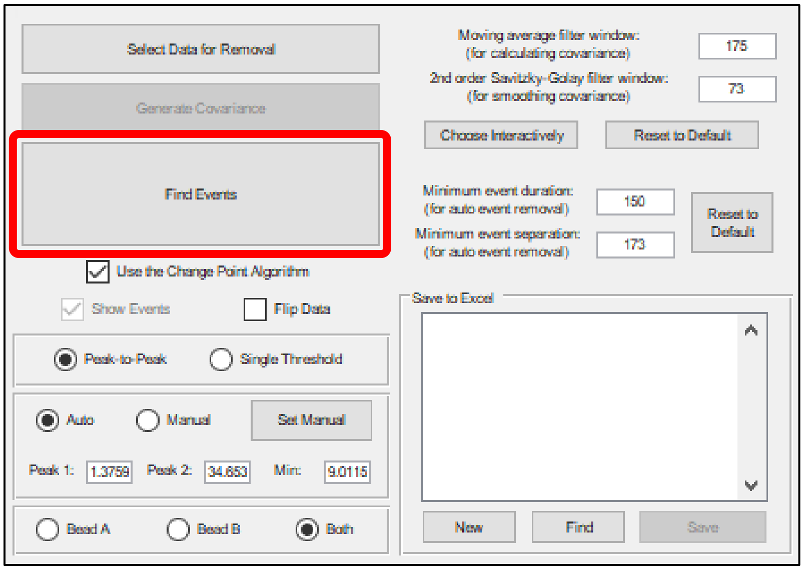

After you do so, **SPASM** will first identify events using the covariance and either the peak-to-peak method or the single threshold method, depending on which method you chose. These initial events are shown on top of the covariance in red and represent all events determined solely by the covariance. These events may include events which are later removed after factoring in the minimum event duration and minimum event separation, discussed in Section 4b.

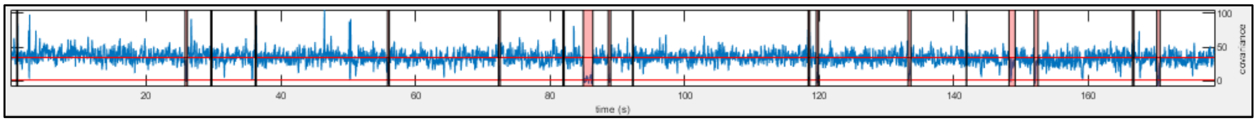

Each of these initial events is passed to the change point algorithm to better determine the event start and end times. For a given event, the change point algorithm analyzes pairs of points within a small window that surrounds the event. It finds the two points within that window which best trisect the data such that the middle section of data is likely drawn from a separate distribution than either outer section.

After the algorithm is finished, some events are automatically removed according to the minimum event duration and minimum event separation, as explained in Section 4b. The final events are shown on top of the bead positions in red.

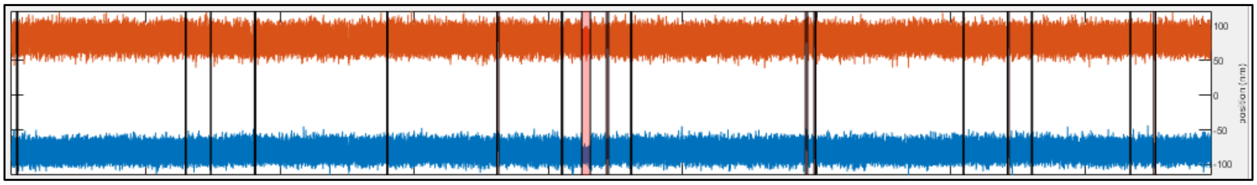

When you hover your mouse over a final event, a label should appear, displaying that event's number among all of the other events.

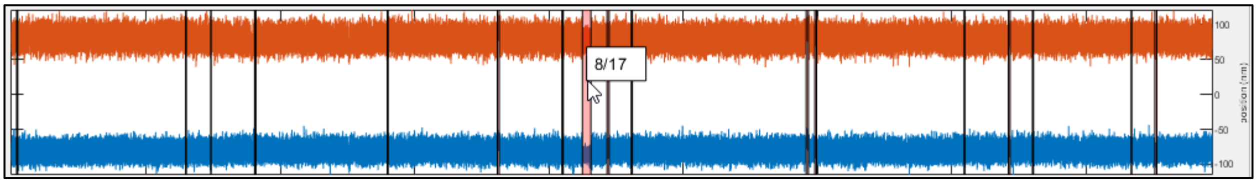

##### Section 4a: Show Events (optional)

The final events are shown in red on top of the bead positions. This can make it hard to visualize each bead's position, and you may wish to toggle the events on and off to see how well they coincide with regions of binding that are visible by eye. You can do so with the **Show Events** checkbox.

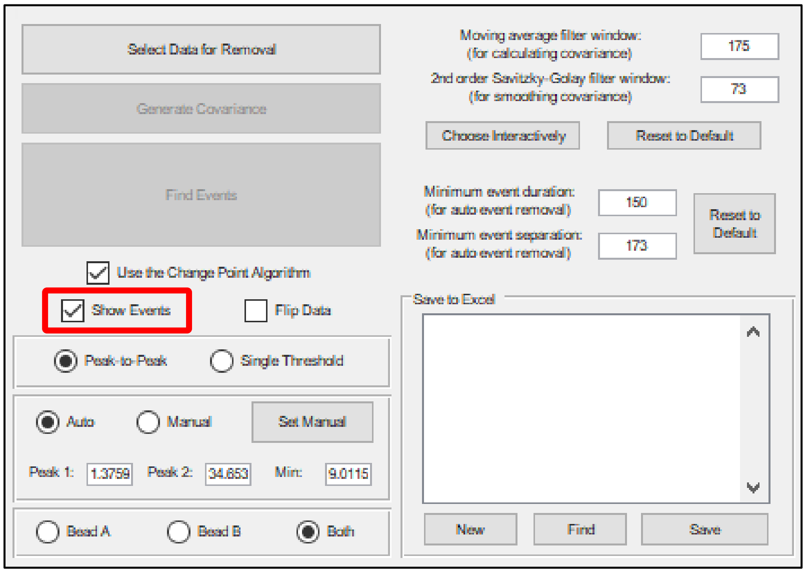

#### Section 4b: Cutoffs (optional)

As explained in the paper, events are removed if (1) they are shorter than the minimum event duration in number of points or (2) they are closer to any other event in number of points than the minimum event separation. This is done to lower the chance of detecting false positive events. The default minimum event duration is 150 points, while the default minimum event separation is 173 points; these values optimized **SPASM**'s performance on a specific set of simulated data. For an in-depth explanation of how these values were determined and guidelines for optimizing these values against your own data, see Appendix C. For more discussion about removed events, see Sections 7b and 7c.

If the default cutoff values are inadequate, you can change them using the corresponding text boxes.

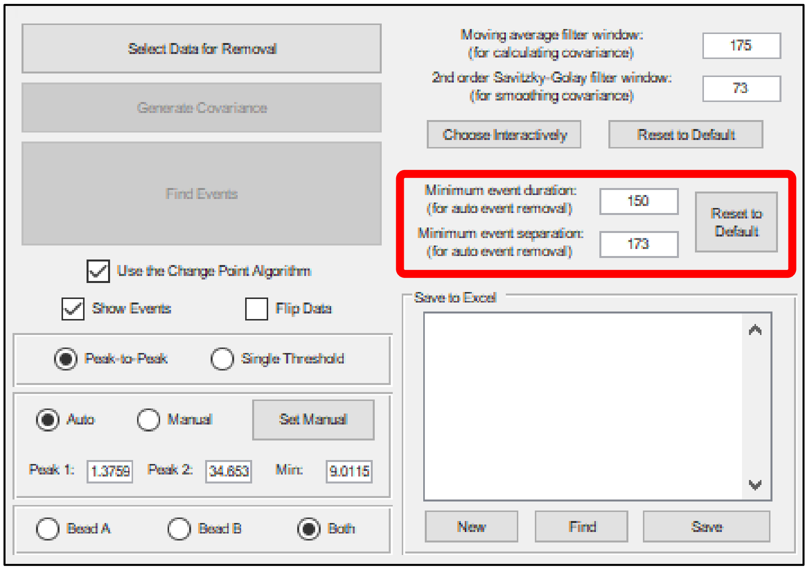

Once either value is changed, all events are automatically reexamined. **SPASM** will remove any events which are longer than the new minimum duration or closer to other events than the new minimum separation. When you click the **Reset to Default** button, each cutoff will return to its default value, and the events will again be reexamined.

#### Section 5: Cumulative Distribution of Event Durations

Additionally, after you click **Find Events**, a cumulative distribution of event durations is plotted, along with a single exponential fit which gives the detachment rate.

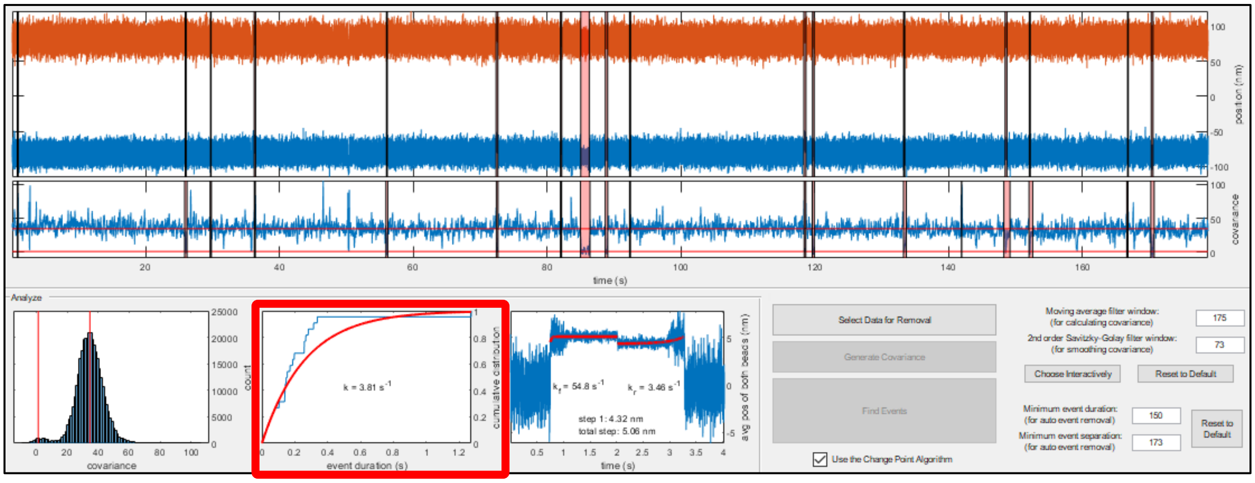

A single exponential distribution is fit to the data (red), and a label gives the fitted rate.

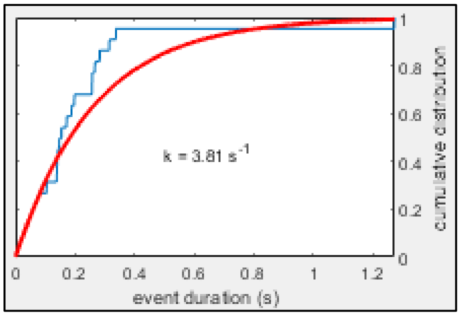

When you right click this plot, a menu appears.

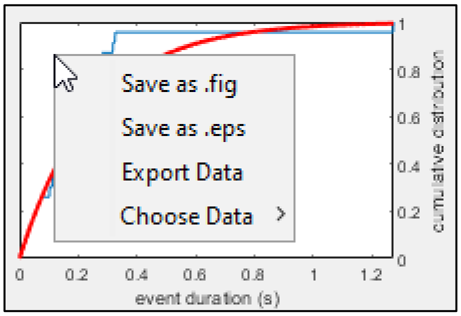

This menu contains **Save as .fig** and **Save as .eps**. These buttons are described in Section 1. The menu also contains **Export Data**. This button allows you to save data from the plot to an Excel workbook. When you click **Export Data**, a window should appear, prompting you to choose a filename and location for this workbook. The created Excel file will contain the plotted data and the fit data, as well as values for the fitted parameter(s).

Finally, the menu contains **Choose Data**. When you hover over **Choose Data**, a submenu will appear. You may then specify which set of data should be plotted.

If you choose **Step 1 Size**, **Step 2 Size**, or **Total Step Size**, the plot will be replaced by a cumulative distribution of the indicated substep sizes during each event. Substep 1 sizes are estimated by averaging 10 ms before and 10 ms after each event start and taking the difference. Total step sizes are estimated by averaging 10 ms before and 10 ms after each event end and taking the difference. Substep 2 sizes are estimated by taking the difference between the total step sizes and the substep 1 sizes.

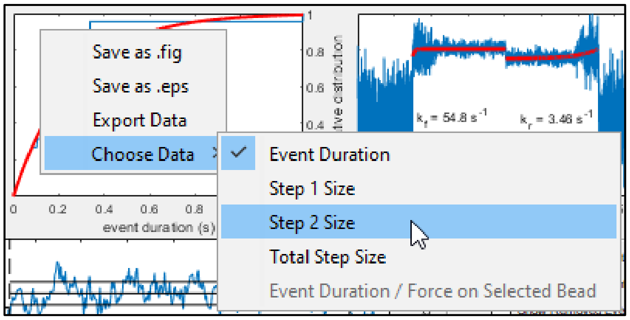

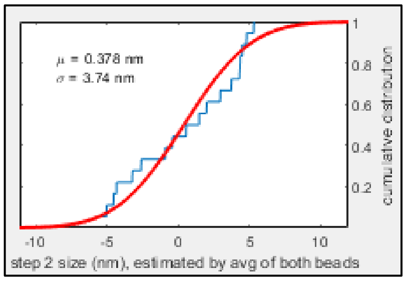

The distribution of step sizes is fit with a normal distribution. The fit will be shown in red, and labels will show the mean and standard deviation of the distribution.

If you previously chose to analyze just one of the beads (see Section 3d), you may also select **Event Duration / Force on Selected Bead**. The plot will then be replaced by a scatter plot, with the duration of each event on the Y axis (with logarithmic scale) and the selected bead's force during each event on the X axis. The force for a given event is calculated by subtracting the force just after detachment, averaged over a 10 ms window, from the average force during the event.

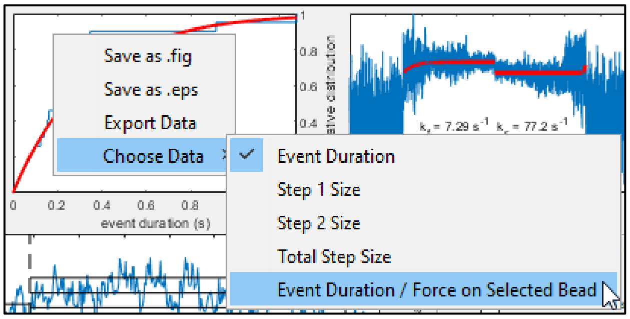

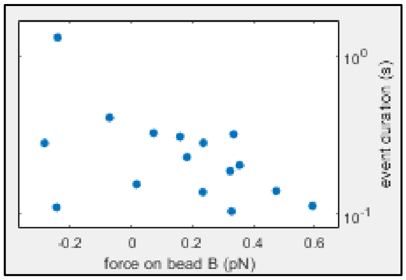

#### Section 6: Ensemble Averages

Next, time forward and time reversed ensemble averages are calculated and plotted.

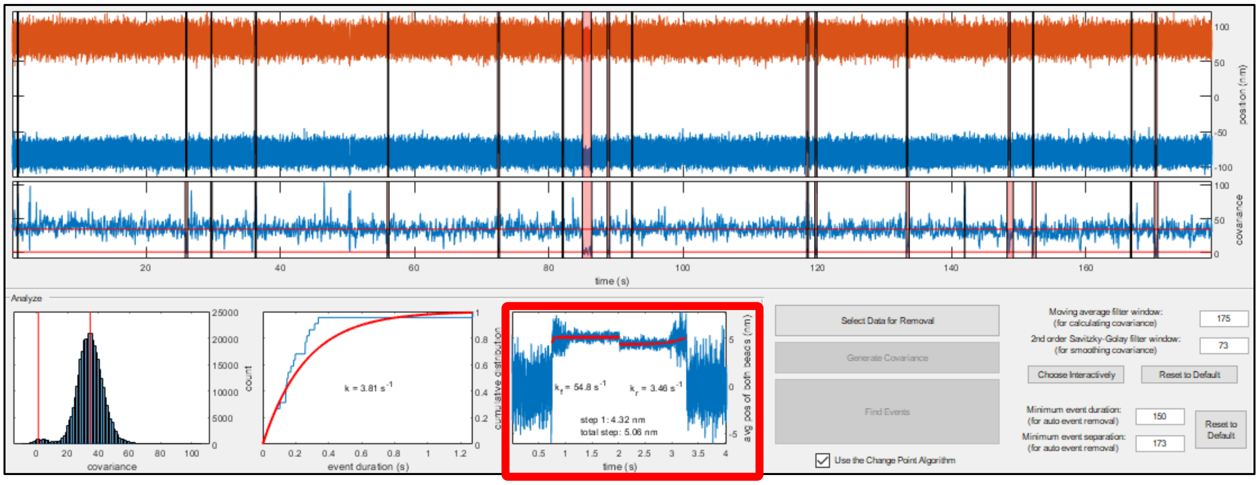

Note that while the X axis scale is correct, the two ensemble averages are plotted one after the other, and so the actual times displayed along the X axis are not meaningful. These times depend on the duration of the longest event contained within the averages.

If the Optimization Toolbox is installed, single exponential curves are fit to each average and are shown in red. Labels give the rates of the fits and estimated substep sizes.

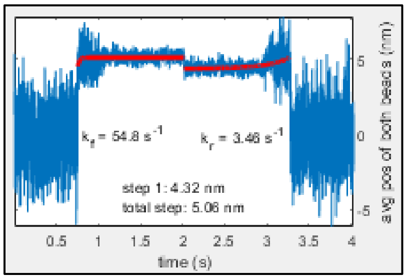

Depending on the shape of each average, MATLAB may take a while to calculate the corresponding fit. If MATLAB takes more than three seconds to calculate a fit, a **Skip** button will appear along the upper edge of the user interface. If you click **Skip**, **SPASM** will continue without fitting the average.

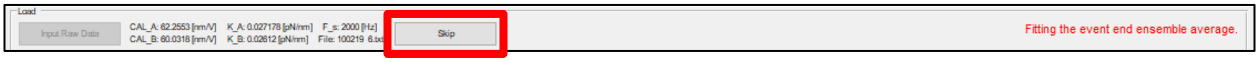

**SPASM** will also continue without the fit if the MATLAB function *lsqcurvefit( )* is unable to solve for the fit after 100,000 attempts.

When you right click the ensemble average plot, a menu will appear.

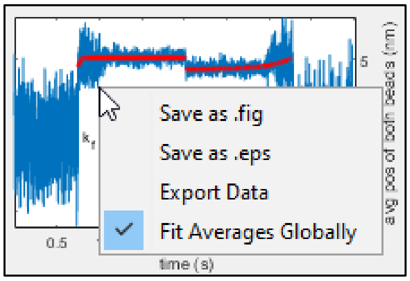

This menu contains **Save as .fig** and **Save as .eps**. These buttons are described in Section 1. The menu also contains **Export Data**, which allows you to save data from the averages to an Excel workbook. When you click **Export Data**, a window should appear, prompting you to choose a filename and location for this workbook. The created Excel file will contain data for the averages and the fits, as well as the fitted rates and the estimated substep sizes. **The fits calculated by SPASM may be suboptimal – different averages may warrant different fitting routines which SPASM cannot anticipate**. You may therefore prefer to export the ensemble averages and fit them using other software or using other algorithms in MATLAB.

Finally, the menu contains **Fit Averages Globally**, which is checked by default. When checked, **SPASM** attempts to fit the time forward and time reversed averages so that they share substep estimates. The time forward and time reversed exponential rates will likely differ. If you uncheck **Fit Averages Globally**, **SPASM** will calculate new fits for the ensemble averages which are independent of one another.

#### Section 7: Individual Event and Likelihood Plots

For each initial event, the change point algorithm analyzes a window of data surrounding that event. By default, these data will be the average position of both beads, but if a certain bead has been selected (Section 3d), these data will instead be that bead's position. The analyzed window of data is shown at the bottom of the user interface. Overlying this plot, vertical dotted lines show the chosen change points, and horizontal solid lines show the mean and standard deviation of the data during the event and before and after the event.

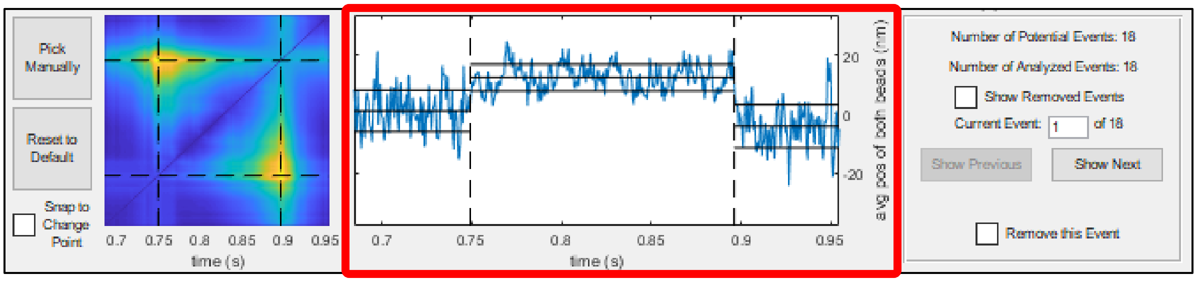

To the left of this plot, a 3D surface plot shows the score given to each pair of points within the window when determining the change points for this event. Both the X axis and the Y axis span the duration of the window, and any given point on this plot, ($t_i$, $t_j$), corresponds to a pair of points within the window. The Z axis – and color – gives the score of each of these pairs. The plot is mirrored across the line X = Y because the point ($t_i$, $t_j$) has the same score as the point ($t_j$, $t_i$). Points in this plot which are closer to the line X = Y correspond to pairs of points in the individual event plot which are closer to each other. Dotted lines show the locations of the chosen change points. These lines intersect at both ($\tau_1$, $\tau_2$) and ($\tau_2$, $\tau_1$), where $\tau_1$ is the estimated start time of the event and $\tau_2$ is the estimated end time of the event.

**Note that this program considers a change point to be the last point before a change occurs, not the first point after the change occurs.**

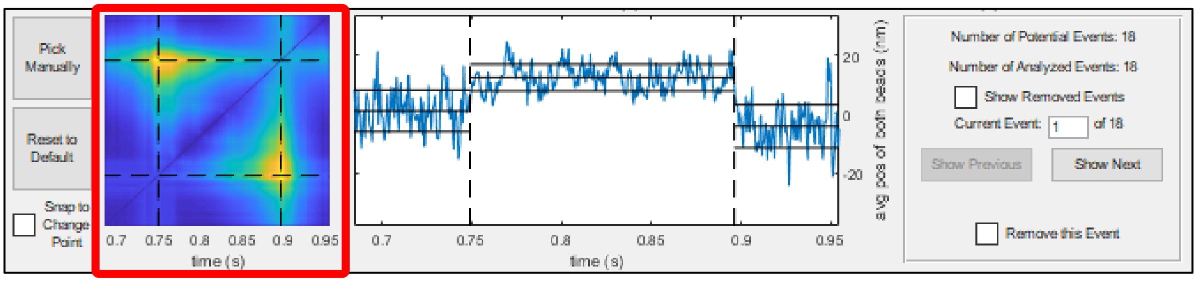

To reduce computational complexity for windows of data longer than 1500 points, **SPASM** only calculates the likelihood for pairs of 1500 data points spaced evenly throughout the window. It uses this likelihood to estimate the change points, and it then searches near each estimate to better determine the optimal values. **This means that for long windows of data, the likelihood surface plot will have a lower resolution than the individual event plot. The dotted lines in the likelihood plot may be slightly inaccurate – each line will be located at one of the 1500 evenly-spaced points, rather than the actual change points. The dotted lines in the individual event plot will be accurate.**

For MATLAB releases R2018b and later: After hovering over this surface plot, a toolbar should appear along the upper edge. You can better visualize the surface by selecting the **Rotate 3D** button in the toolbar and then clicking and dragging the plot with the mouse. Restore to a view from above by clicking **Restore View** in the toolbar.

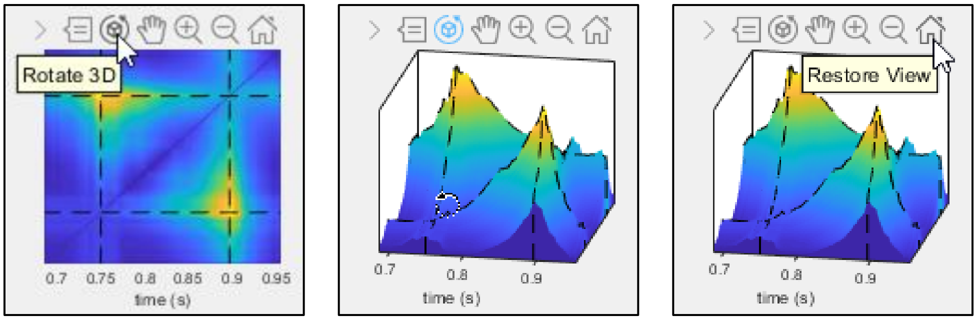

For MATLAB releases prior to R2018b: At the very top of the user interface, there should be a button labeled **Tools**. When you click this button, a submenu should appear, showing several more buttons for interacting with plots. You can better visualize the surface by selecting the **Rotate 3D** button and then clicking and dragging the plot with the mouse. Restore to a view from above by clicking **Restore View**.

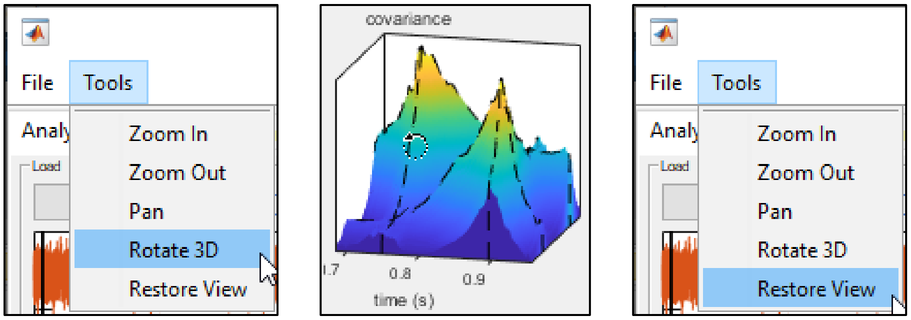

Often, the change points will correspond to the highest peak in the likelihood plot. However, **SPASM** requires that the variance of the data during the event be less than the variance of the data before or after the event. **It is possible that the chosen change points will correspond to a lower peak in the likelihood plot in order to satisfy this requirement.**

A panel to the right shows which event is currently being displayed in these plots. This panel also shows the total number of events that were initially identified by the covariance (**Number of Potential Events**), as well as the number of events after automatic removal due to the user-defined minimum event duration and minimum event separation values (**Number of Analyzed Events**).

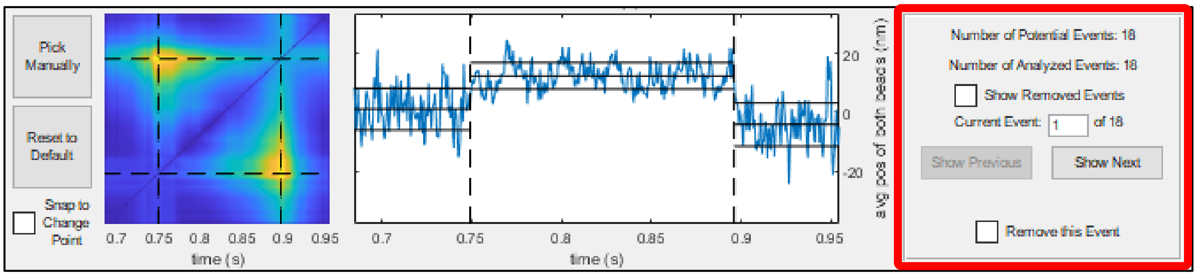

#### Section 7a: Change the Current Event (optional)

There are two ways to change which event is displayed in the individual event and likelihood plots.

##### i. Text Box

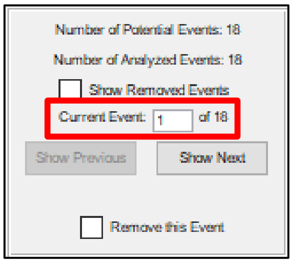

Enter an integer value in the text box. To determine the number for a specific event, hover over that event in the plot of the bead positions, as described at the beginning of Section 4.

##### ii. Show Previous / Show Next

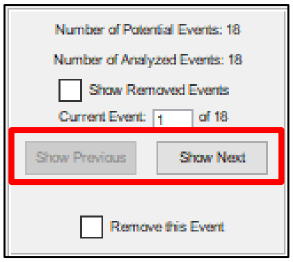

Alternatively, click the **Show Previous** and **Show Next** buttons to navigate through the events one by one.

#### Section 7b: Remove an Event (optional)

As explained in the paper and in Section 4b of this user guide, some events are automatically removed according to the minimum event duration and minimum event separation. Removed events are excluded from the analysis. They do not contribute to the event duration plot, substep size plots, duration per force plot, or the ensemble average plots. They will no longer be shown in red on top of the bead positions (although they will still be shown in red on top of the covariance). Finally, they will not be saved to Excel (described in Section 9).

You have the option to manually remove an event if you do not think it should be included in the data analysis. To do so, the event must currently be showing in the individual event and likelihood plots. Refer to Section 7a for instructions on how to set which event is showing in these plots. To remove the current event, check the **Remove this Event** checkbox.

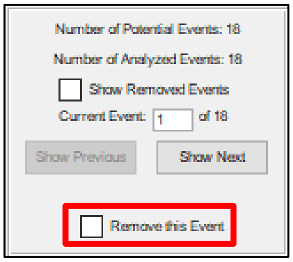

When you remove an event, plots will be updated.

#### Section 7c: Show Removed Events (optional)

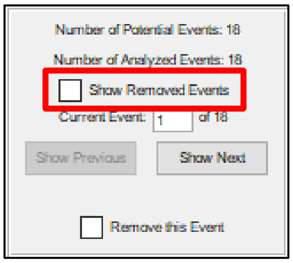

By default, the individual event and likelihood plots will only display events that are used in the analysis. When **Show Removed Events** is checked, you may also view events which have been removed. Note that the numbering of certain events might change after you check **Show Removed Events**. You can determine any event's number by hovering over it in the plot of the bead positions, as described at the beginning of Section 4. Although removed events do not show in the plot of the bead positions, they will appear if you hover over them.

When a removed event is displayed in the individual event and likelihood plots, a red label will give the reason for its removal:

- **dur**: The event was removed automatically due to the minimum event duration.
- **sep**: The event was removed automatically due to the minimum event separation.
- **user**: The event was removed manually.

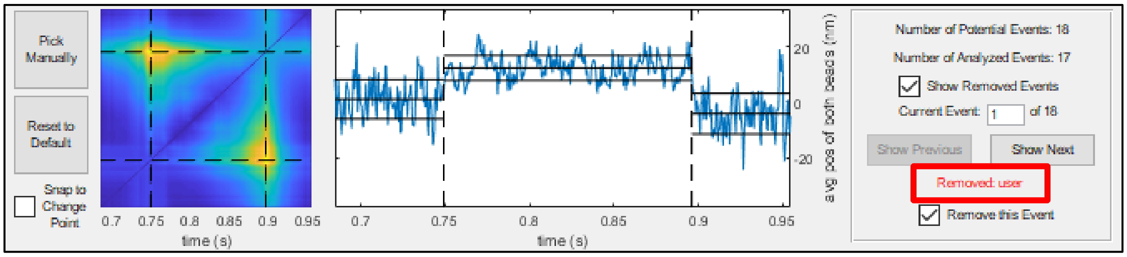

The label will indicate if an event has been removed for multiple reasons.

You may opt to include an event which has been previously removed, either automatically (by the minimum event duration or minimum event separation) or manually. To do so, uncheck the **Remove this Event** checkbox. A green label will indicate if you include an event which would otherwise be automatically removed.

#### Section 7d: Pick Change Points Manually (optional)

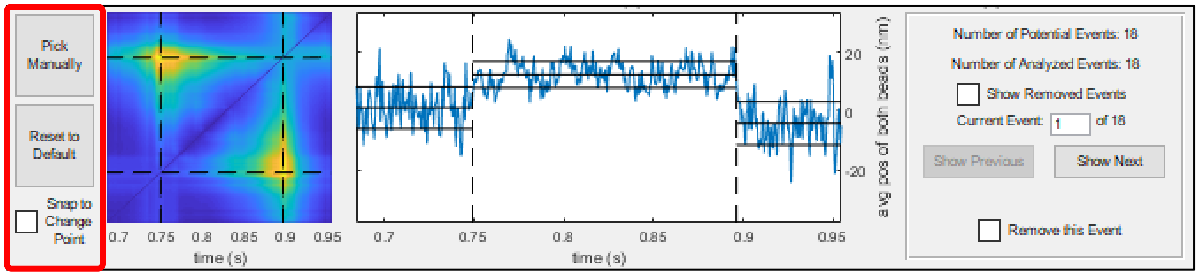

Finally, to the left of the likelihood plot, two buttons and a checkbox will appear. When you click the first button, **Pick Manually**, **SPASM** will wait for you to specify new change points for the current event by either

(1) clicking on the likelihood plot once or

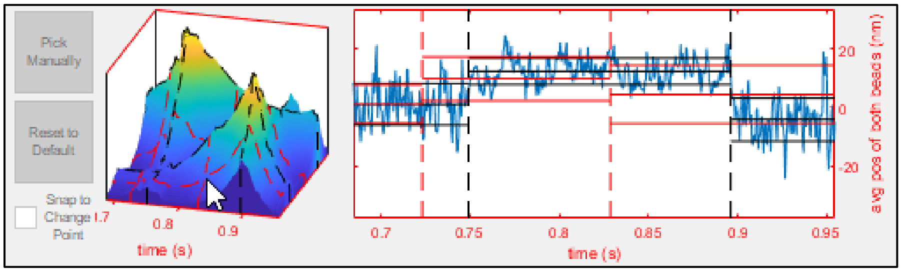

(2) clicking on the individual event plot twice.

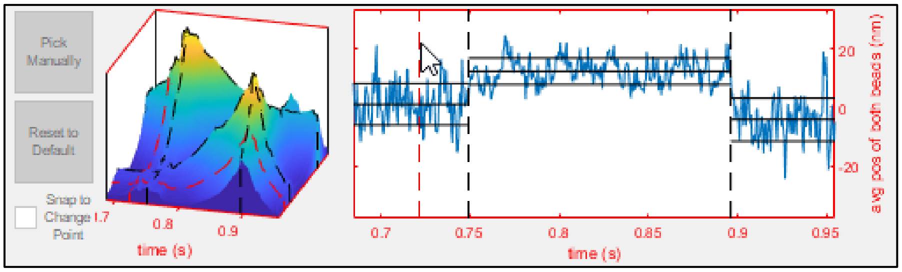

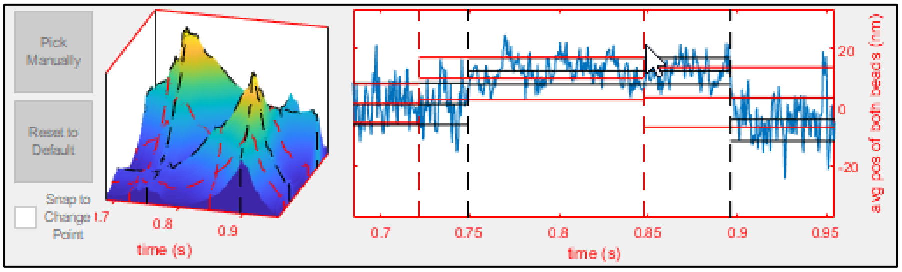

When **Snap to Change Point** is unchecked, each new change point will appear exactly where your cursor is. When **Snap to Change Point** is checked, **SPASM** will instead analyze a 50 point window of data centered at your cursor. The new change point will be placed at the point with the highest likelihood, as determined by the change point algorithm. This feature is available to help you easily pick obvious change points.

As the new change points are chosen by clicking within either plot, you are limited to only those points which are shown in the plots. For example, in the three pictures shown previously, you could not choose to set one of the change points at 1.0 seconds because the time spans from about 0.68 seconds to about 0.95 seconds. If you want to set a change point at a time which is not showing in the plots, you must first pan the individual event plot as needed (or alternatively zoom out) to show the new data. Panning the likelihood plot will not show new data.

For MATLAB releases R2018b and later: After hovering over the individual event plot, a toolbar should appear along the upper edge. You can enter pan mode by selecting the **Pan** button.

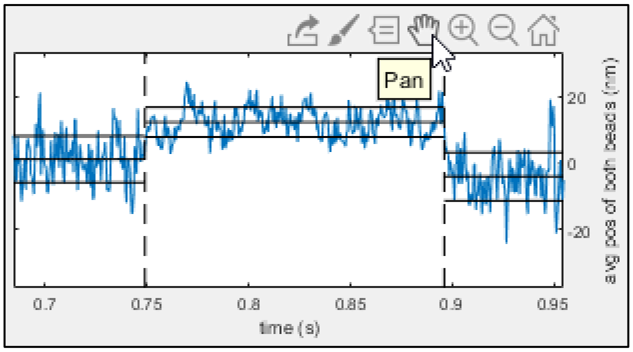

For MATLAB releases prior to R2018b: At the very top of the user interface, there should be a button labeled **Tools**. When you click this button, a submenu should appear, showing several more buttons for interacting with plots. You can enter pan mode by selecting the **Pan** button.

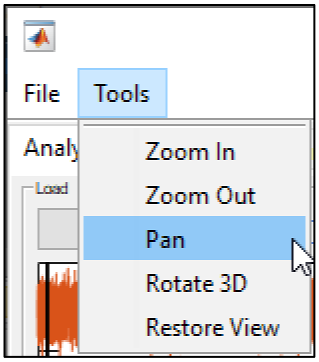

You can then use the mouse to click and drag the individual event plot. The likelihood plot will update accordingly.

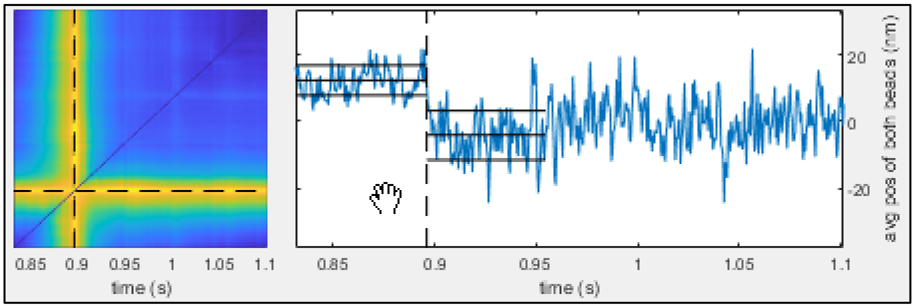

Note that you must click the **Pan** button again to deactivate pan mode and return to analysis. You may need to zoom out to show both desired change points before you click **Pick Manually**.

The likelihood plot updates because it always shows the likelihood for whatever window of data is plotted in the individual event plot. If this window is longer than 1500 points, then the likelihood plot will only show the likelihood for pairs of 1500 points spaced evenly throughout this window (see Section 7 for more explanation).

After you set an event's change points, the event is reevaluated – if it should be removed due to the minimum event duration or minimum event separation, it will be. Plots, such as the ensemble averages plot, are then updated.

If you click the **Reset to Default** button, the change points will return to their original values, the event will be reevaluated, and plots will update accordingly.

#### Section 8: Recalculate Events and Update Plots (optional)

By default, calculating the events involves running the change point algorithm on each potential event. If all of the events need to be recalculated, **SPASM** will not automatically rerun the change point algorithm this many times, because doing so can take a while. (Other updates, such as recalculating the covariance histogram peaks after the window sizes are changed, do occur automatically.)

After the events have been found, changing anything about how the events were initially identified or how the change points were calculated will cause the **Find Events** button to again become enabled. It will now read **Recalculate Events and Update Plots**. This will happen if you

(1) change the method of event detection (Section 3a),

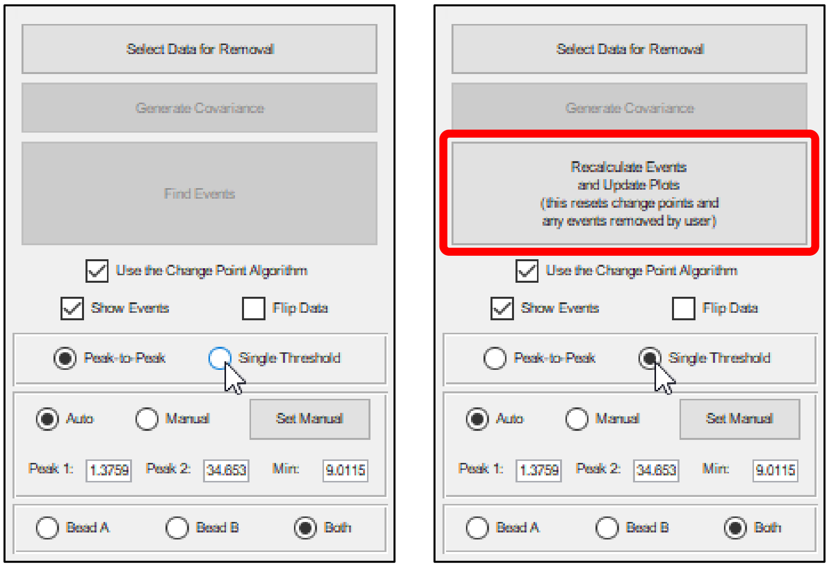

(2) change either filter window size (Section 3b),

(3) change the covariance histogram peaks or minimum, depending on which method of event detection was used (Section 3c),

(4) change which bead(s) should be analyzed (Section 3d), or

(5) change whether the change point algorithm should be used (Section 3e).

When any of the above changes take place, **many plots, including the ensemble averages, will not update**. You can tell **SPASM** to recalculate the events and update these plots by clicking **Recalculate Events and Update Plots**. If you do so, however, you will lose any changes you had made by manually removing events (Section 7b) or manually picking change points (Section 7d).

As long as the method of event detection, covariance histogram peaks or minimum, filter window sizes, analyzed bead(s), and method of transition detection match those which were used to determine the current set of events, then the **Find Events** button will stay disabled, all plots will remain up to date, and you will be able to save your data to an Excel workbook (Section 9).

#### Section 9: Save to Excel

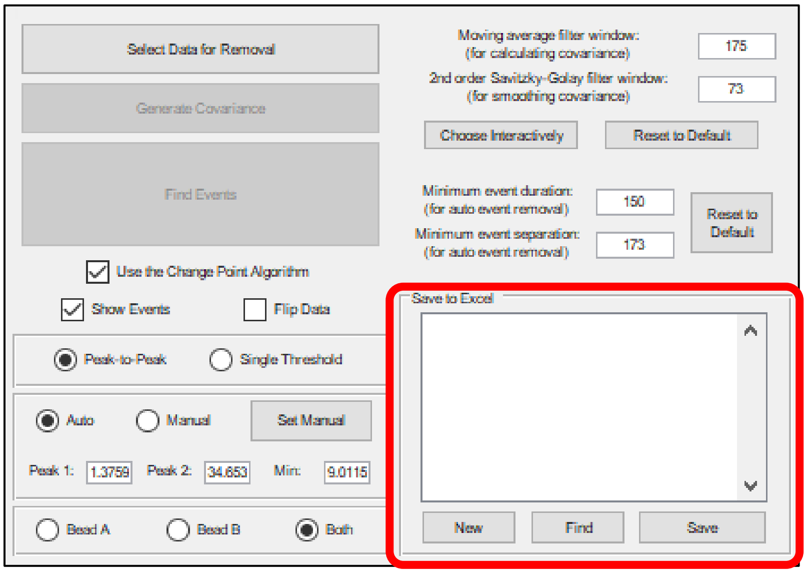

A panel labeled **Save to Excel** is located in the right lower corner of the user interface. This panel contains an empty list and three buttons.

##### a. New

At any point, you may click **New** to create a new Excel workbook. When you click **New**, a window should appear, prompting you to choose a filename and location for the workbook.

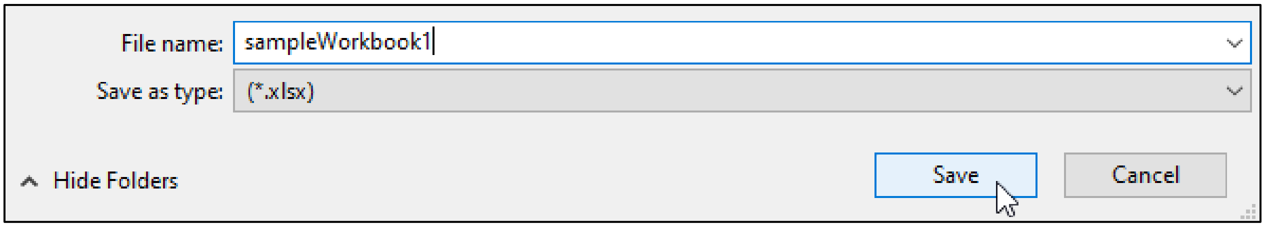

**SPASM** will create the blank file, and its filename should be added to the list.

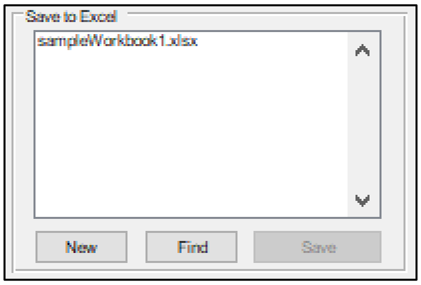

##### b. Find

Alternatively, you may click **Find** to locate an Excel workbook which already exists. When you click **Find**, a window should appear, prompting you to select a .xlsx file.

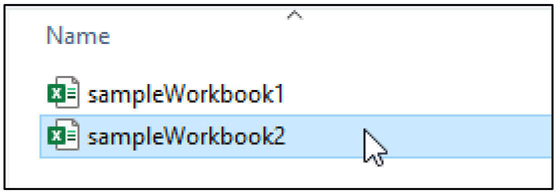

After you select a file, its filename should be added to the list.

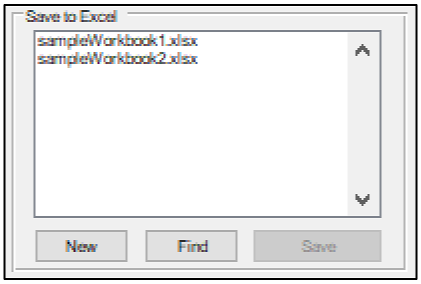

While only the filenames are displayed in the list, the entire path of each file is stored. You may select any of the files in the list by clicking on them. Select multiple files by holding the Control key or Command key (depending on your operating system) while clicking.

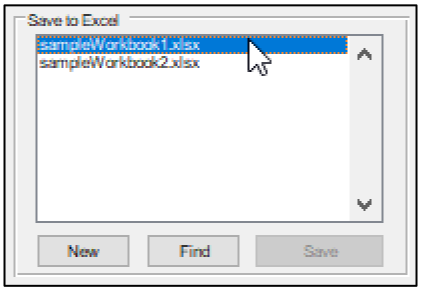

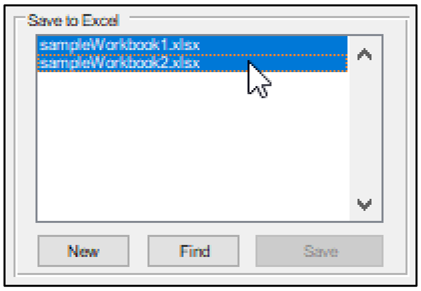

**c. Save**

The **Save** button becomes enabled after events have been found. When you click **Save**, the results of data analysis will be added to the next blank sheet of each selected Excel workbook in the list. This newly created Excel sheet will include

I. general information about the analyzed data, such as the full path of the .txt file, the sampling frequency, and the number of analyzed events;

II. all of the parameters used in data analysis, such as the window sizes or the covariance histogram peak locations, so that the data analysis can be repeated; and

III. a table which contains, for each event,

- the event start and end times, in both points and seconds;
- the duration of the event, in both points and seconds;
- the amount of time separating the event from its neighboring events, in both points and seconds;

- the position of each bead just before and just after the event start, averaged over 10 ms windows;
- the position of each bead just before and just after the event end, averaged over 10 ms windows;
- the estimated substep 1, substep 2, and total step sizes based on each bead's position;
- the force on each bead during the event, averaged over the entire event;
- the force on each bead before and after the event, averaged over the entire time separating the event from its neighboring events; and
- the force on each bead just before and just after the event end, averaged over 10 ms windows.

By using the **Save to Excel** panel, you can save the analysis results for multiple .txt files to a single Excel workbook. The results for each .txt file will be saved to a unique sheet. That workbook can then be used as the input in the second tab of **SPASM**, **Combine Multiple Files** (see Section 11). There, **SPASM** will go through each sheet in the workbook and locate the .txt file referenced in that sheet. It will then perform data analysis on all of the .txt files combined. **If you save the results from a .txt file to a workbook and then move or rename that .txt file, SPASM will not be able to locate it. You will either need to manually change the path name inside the corresponding sheet of the Excel file or keep a copy of the .txt file in its original location.**

#### Section 9a: Flip Data (optional)

When preparing a set of .txt files to be analyzed together, some of these files may contain data which is oriented in the opposite direction as data in other files. However, when the analysis is performed, it is important that each set of data is oriented in the same way. As an example, it would be okay combine these,

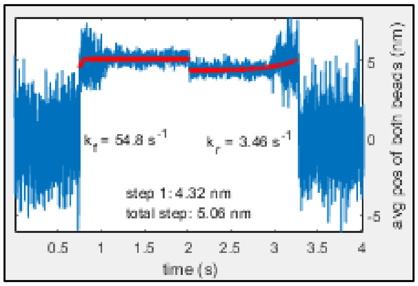

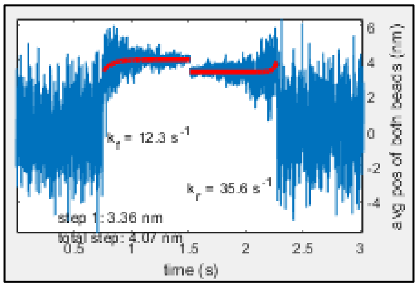

while these should not be combined.

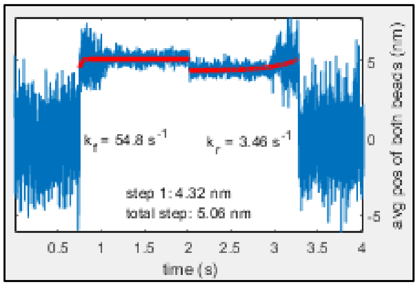

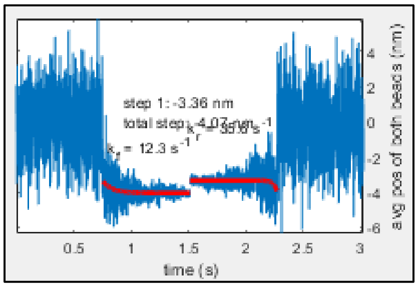

Use the **Flip Data** checkbox to flip a set of data vertically.

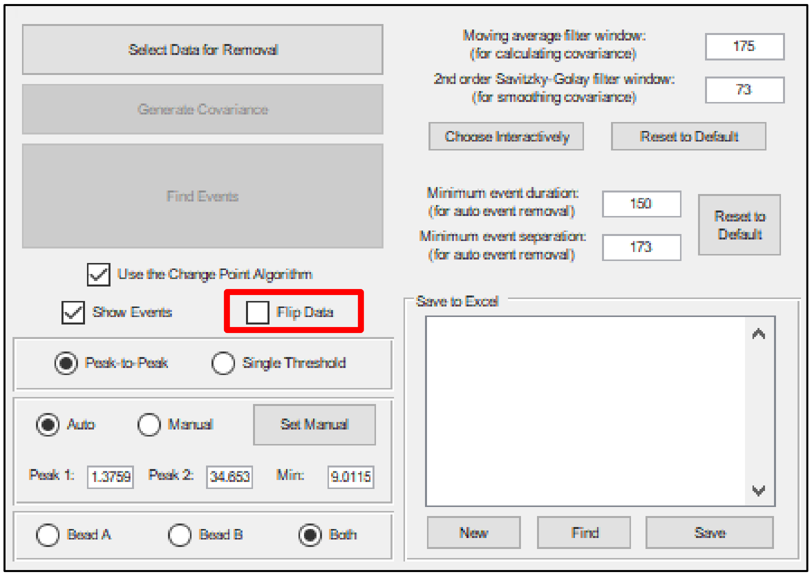

When you save that set of data to Excel, the Excel sheet will indicate whether or not it has been flipped. Then, in the **Combine Multiple Files** tab, **SPASM** will know whether a particular data set needs to be flipped in order to orient it in the same way as the other data sets.

#### Section 10: Error Messages

If **SPASM** encounters incorrect or unexpected user input, it may display an error dialog box

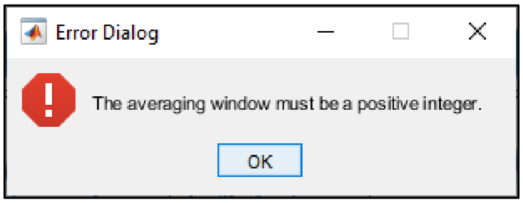

or a warning dialog box.

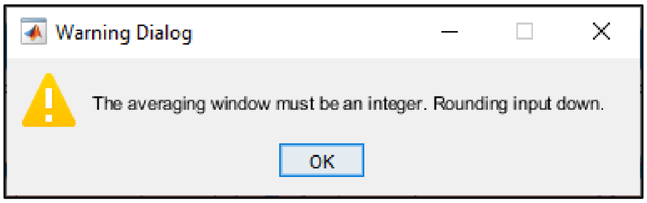

Alternatively, it is possible that **SPASM** may encounter an unexpected error which it is not able to handle.

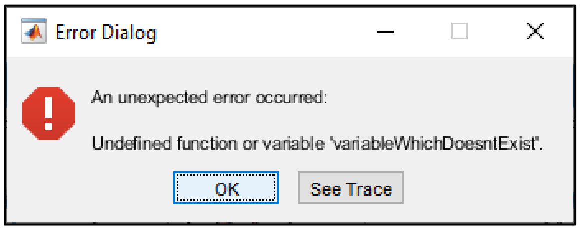

Click **See Trace** if you would like to see and/or copy the full error message.

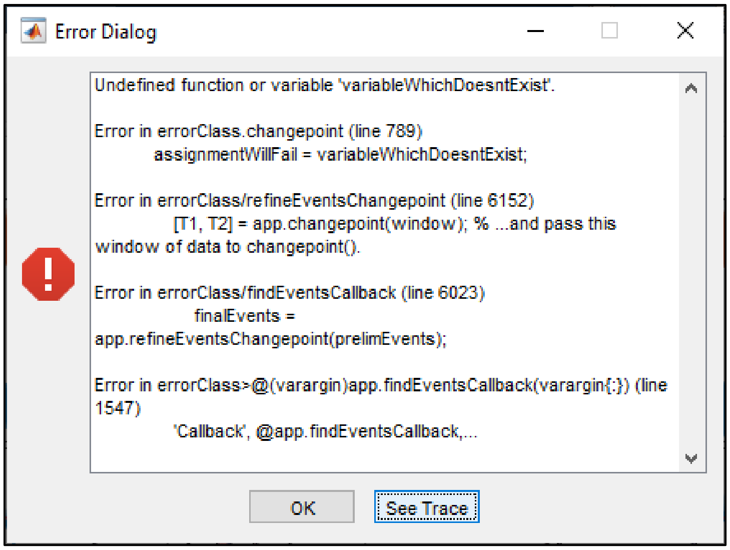

**SPASM** will then attempt to reset to a previous state. It may need to reset completely or exit.

#### Combine Multiple Files

##### Section 11: Choose Files

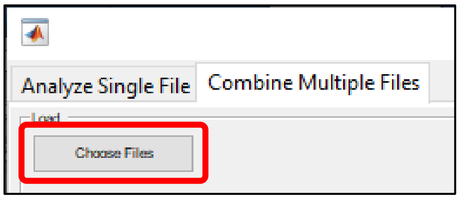

To load multiple files, go to the second tab of **SPASM**, labeled **Combine Multiple Files**, and click the button labeled **Choose Files**. A window should appear, prompting you to select a .xlsx file. Choose a .xlsx file that was created by the first tab, **Analyze Single File**.

Once the selected .xlsx file is read, a table will appear. This table will show each individual file's name, the total number of events found in that file, and a checkbox indicating whether that file should be included in the combined data analysis (see Section 14).

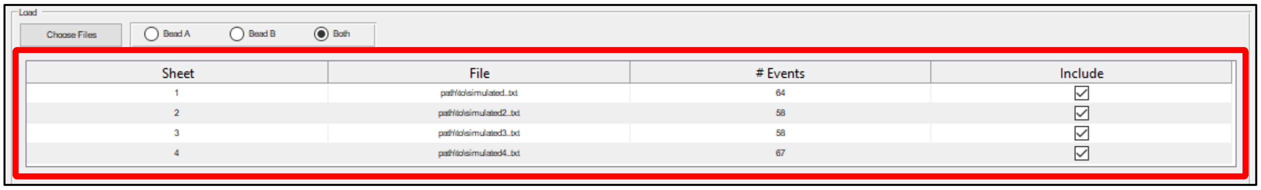

#### Section 12: Cumulative Distribution of Event Durations

Additionally, a cumulative distribution of event durations is plotted by default. Other plots, including cumulative distributions of step and substep sizes, can be selected (see below).

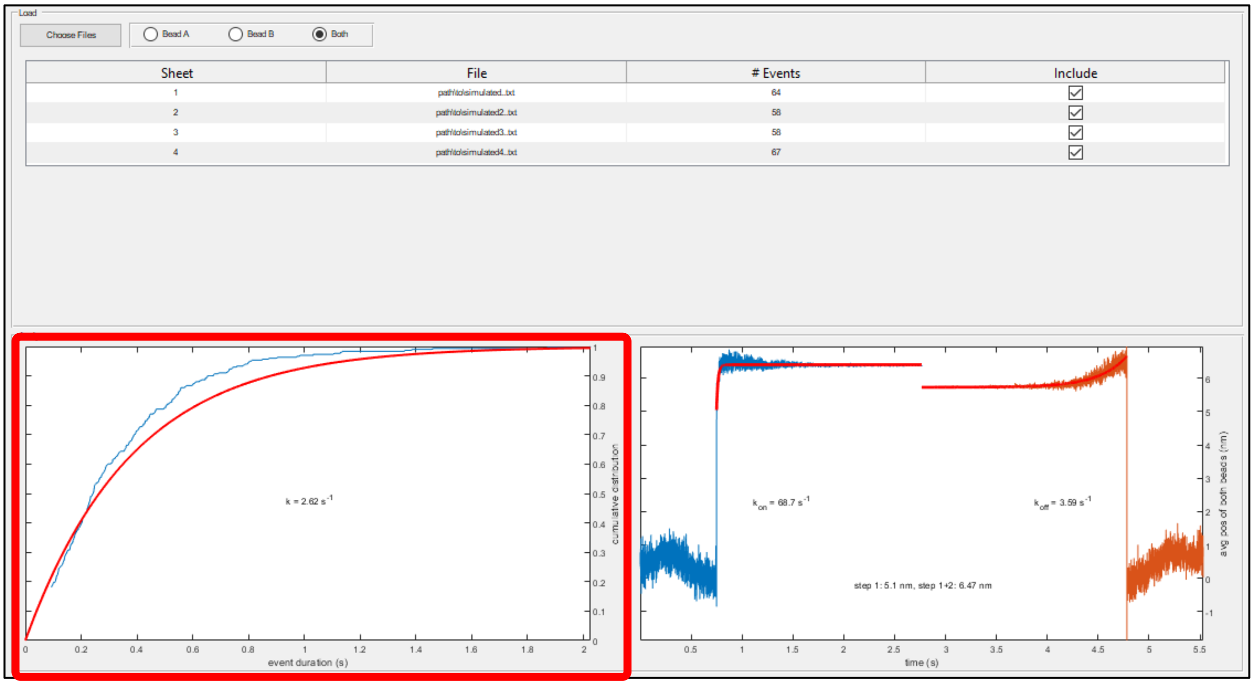

An exponential distribution is fit to the data. The fit is shown in red, and a label gives the fitted rate (i.e. the detachment rate).

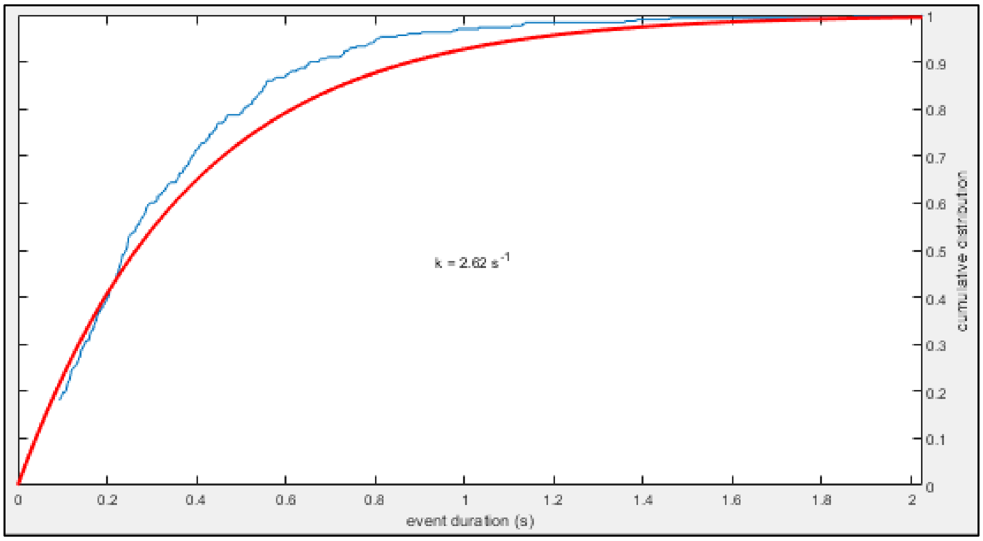

As with previous plots, when you right click within this plot, a menu appears. You may click **Save as .fig** or **Save as .eps**, both described in Section 1. For this plot, as before (see Section 5), you may also click **Export Data**, which will save data from the plot to an Excel file. Finally, hovering over **Choose**

**Data** will again bring up a submenu, and you may then specify which set of data should be plotted (e.g. substep 1 size). Refer to Section 5 for detailed descriptions of each option.

#### Section 13: Ensemble Averages

Ensemble averages are plotted to the right of the cumulative distribution plot.

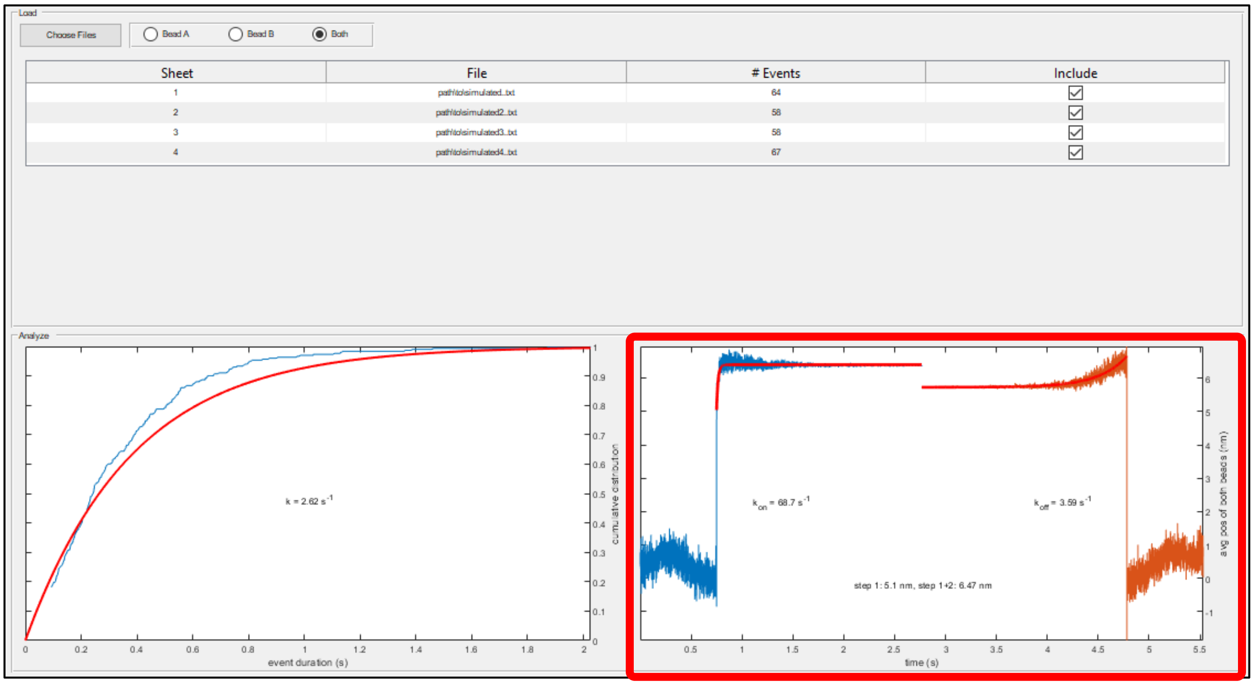

If the Optimization Toolbox is installed, exponential curves are fit to each average and are shown in red. Labels give the rates of the fits and estimated step sizes.

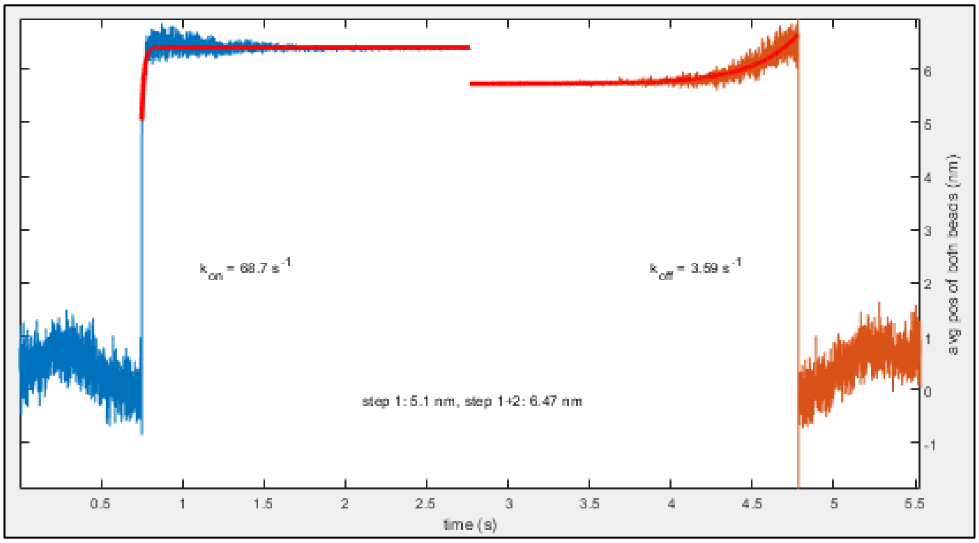

As with previous plots, when you right click this plot, a menu appears. You may click **Save as .fig** or **Save as .eps**, both described in Section 1. For this plot, as before (see Section 6), you also have the option to click **Export Data**, which will save data from the plot to an Excel file, or uncheck **Fit Averages Globally**, which will calculate fits for the two ensemble averages independently.

#### Section 14: Selectively Include Data (optional)

To exclude a file from the combined data analysis, you may uncheck the corresponding checkbox in the table. Once you do so, the plots will be updated.

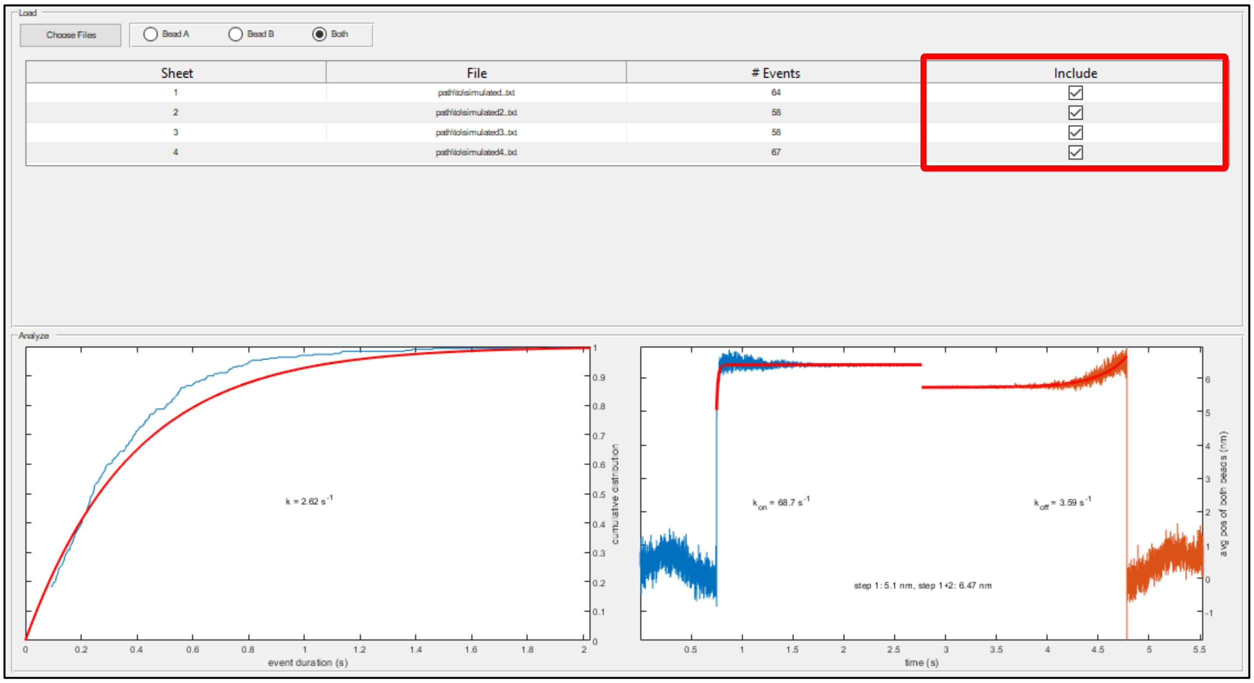

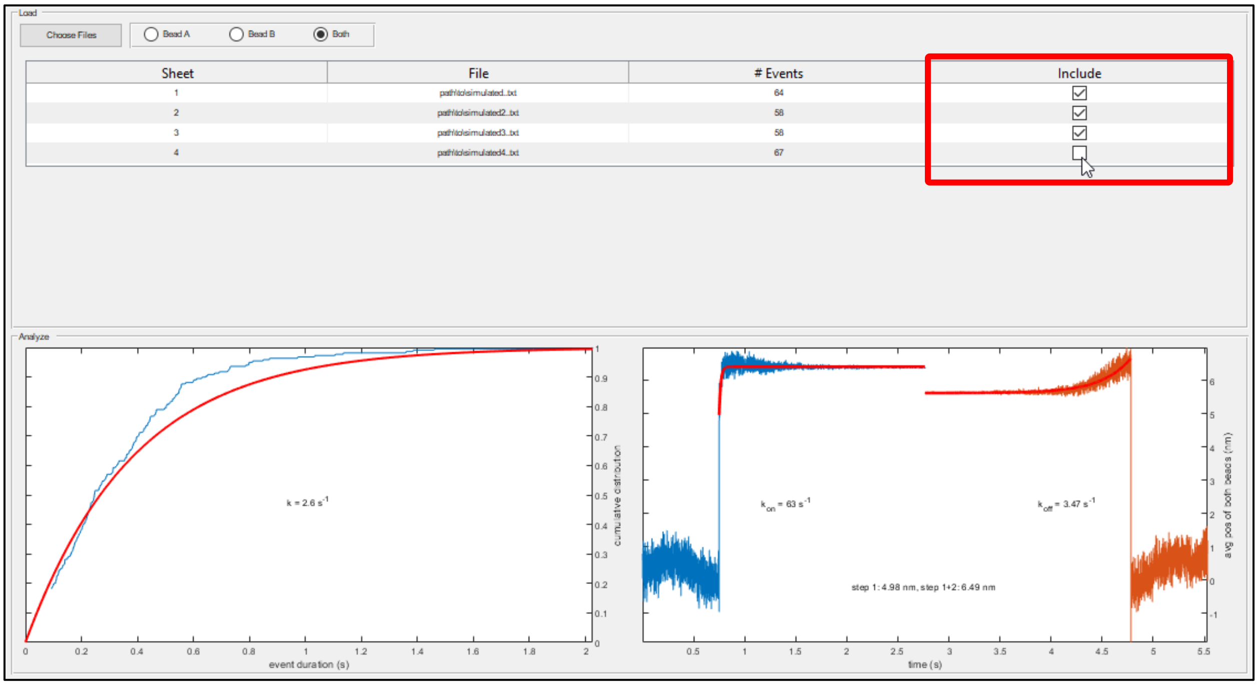

There must always be at least one file which is included.

#### Section 15: Select One Bead for Analysis (optional)

As before (see Section 3d), you may choose to analyze only one of the beads. Doing so will update the plots. By default, both beads will be analyzed, even if events were detected using just one of the beads.

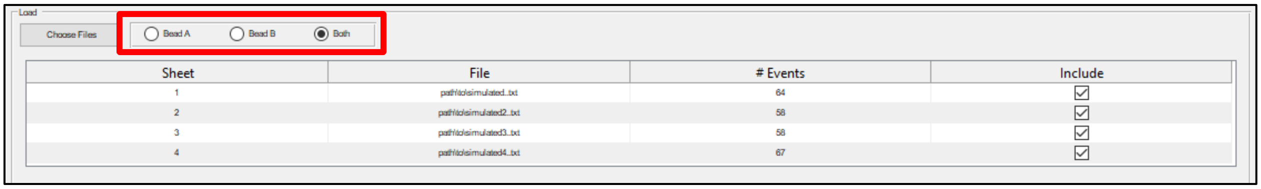

#### Appendix A: Header and Data Formatting

##### Section 1: Default Format

**SPASM** assumes that input text files have the following format:

| | | | | |
|---|---|---|---|---|
| key1 | value1 | | | header |
| ... | ... | | | |
| keyN | valueN | | | |
| colName1 | colName2 | ... | colNameM | column names |
| data(1,1) | data(1,2) | ... | data(1,M) | data |
| ... | ... | ... | ... | |
| data(L,1) | data(L,2) | ... | data(L,M) | |

Here, columns of text are tab-delimited. The following must be true:

(1) All data must be numeric text (e.g. '1.0'), and the first row of data must be the first row in the file which contains only numeric text.

(2) There must be keys labeled 'K1', 'K3', 'CAL1', 'CAL3', and 'Sample Rate', with corresponding numeric values. These values determine the trap stiffnesses for bead A (K1) and bead B (K3), in units of pN/nm, the volt-to-nanometer conversion factors for bead A (CAL1) and bead B (CAL3), in units of nm/V, and the sampling frequency (Sample Rate), in $s^{-1}$.

(3) Two of the column names must equal 'Trap1X' and 'Trap2X'. The corresponding columns will be treated as the positions of beads A and B, respectively, in units of volts.

You may refer to any of the sample files provided with **SPASM** for an example of valid formatting.

#### Section 2: Custom Input Function

The above restrictions are set in place by a function that **SPASM** uses to read text files. This function will fail if you attempt to read a text file without the proper format, and an error message will be displayed.

If you are running the uncompiled version of **SPASM**, you may supply your own function for reading text files. In this way, you have greater flexibility when it comes to supplying data. The drawback is that you will need to write this function and tailor it to the format of your data. This section aims to show how you might go about doing this.

If you choose to supply your own function for reading data, it must satisfy the following requirements:

A. It must accept one input argument. This will be a character vector containing the full path of the input text file.

B. It must return 7 output arguments. These are, in order:

   1. The sampling frequency, in $s^{-1}$.
   2. The trap stiffness of bead A, in pN/nm.
   3. The trap stiffness of bead B, in pN/nm.
   4. The volt-to-nanometer conversion factor of bead A, in nm/V.
   5. The volt-to-nanometer conversion factor of bead B, in nm/V.
   6. A MATLAB table. Two of the columns of this table must be named 'BeadAPos' and 'BeadBPos'. The data in these columns are treated as the positions of beads A and B, respectively, in units of volts. The number of rows in this table equals the number of data points, and the rows are ordered chronologically.
   7. A $7^{th}$ variable which does not affect data analysis. This variable only affects how your data will be saved to a text file, should you choose to save it (see User Guide Section 2, Save), and it therefore allows the input function to directly communicate with the output function. For more information, see the next section, **Custom Output Function**.

Consider the following text file.

```
10000.000       2000
A y     A x     B y     B x     Feedback
0.002   0.026   0.009   0.717   -0.001
0.014   0.029   0.002   0.711   0.001
-0.001  0.023   0.004   0.713   0.000
```

The one-line header in this file contains the total number of rows of data (10,000) and the sampling frequency, in $s^{-1}$ (2000). This file is not compatible with **SPASM**, as it does not satisfy any of the three requirements laid out in Section 1 of this appendix. The following MATLAB function, **myCustomInputFunction**, was written to successfully read this file:

```
function [fs, KA, KB, CALA, CALB, tData, header] = myCustomInputFunction(file)
    fileID = fopen(file); % Open the file.
    header = textscan(fileID, '%s%s', 1, 'Delimiter', '\t'); % Read the one-line
      header into a cell array. header{1}{1} contains the total number of rows of
      data, as a character vector. header{2}{1} contains the sampling frequency,
      as a character vector.
    fclose(fileID); % Close the file.

    % Set the K and CAL values manually, as the file does not contain these values.
    KA = 0.05;
    KB = 0.05;
    CALA = 40;
    CALB = 40;

    % Set the sampling frequency based on the header. fs must be numeric, so
      convert from a character vector to a double with str2double( ).
    fs = str2double(header{2}{1});

    % Read the rest of the file into a MATLAB table. Tell readtable( ) that there
      is only 1 header line.
    tData = readtable(file, 'HeaderLines', 1);

    % Rename the 2nd and 4th columns to 'BeadAPos' and 'BeadBPos'.
    tData.Properties.VariableNames{2} = 'BeadAPos';
    tData.Properties.VariableNames{4} = 'BeadBPos';
end
```

You may now supply this function as an argument when starting **SPASM**. There are two ways to do this. First, you can create a function handle which references your function, by prepending the @ symbol to your function's name.

```
Command Window
fx >> SPASM(@myCustomInputFunction)
```

MATLAB must be able to find your custom function (i.e., it must be in your Current Folder or on your MATLAB path). Alternatively, you can provide the full path of your function as a character vector.

```
Command Window
fx >> SPASM('path/to/myCustomInputFunction.m')
```

Here is another custom function, **myCustomInputWhichUsesUI**, which uses MATLAB functions *uiimport( )* and *inputdlg( )* to display user interfaces that let you interactively tell MATLAB how to read the input file.

```
function [fs, KA, KB, CALA, CALB, tData, header] = myCustomInputWhichUsesUI(file)
    % 1st Import Tool.
    H = uiimport(file); % Open the Import Wizard. Store the output struct into H.
    Hfields = fieldnames(H); % Get the names of H's fields (there should be 1).
    header = H.(Hfields{1}); % Store the data contained in the first field of H.
    fs = header(2); % Grab the sampling frequency from the second value.

    % Input dialog box.
    A = inputdlg({'KA','KB','CALA','CALB'}); % Create the input dialog box.
    KA = str2double(A{1}); % Convert inputs from character vectors to doubles.
    KB = str2double(A{2});
    CALA = str2double(A{3});
    CALB = str2double(A{4});

    % 2nd Import Tool.
    D = uiimport(file); % Open the Import Wizard again. D should have 2 fields.
    Dfields = fieldnames(D);
    for i = 1:length(Dfields)
        field = D.(Dfields{i});
        if size(field,1) > 2 % The data which we want has more than 2 rows.
            tData = array2table(field);
        end
    end

    % Rename the 2nd and 4th columns to 'BeadAPos' and 'BeadBPos'.
    tData.Properties.VariableNames{2} = 'BeadAPos';
    tData.Properties.VariableNames{4} = 'BeadBPos';
end
```

If we supply this function to **SPASM** and try to read the above text file, we will first see MATLAB's **Import Tool**. If MATLAB thinks this file contains no header lines, it will read the first line as data and then stop at the second line, as the second line is formatted differently.

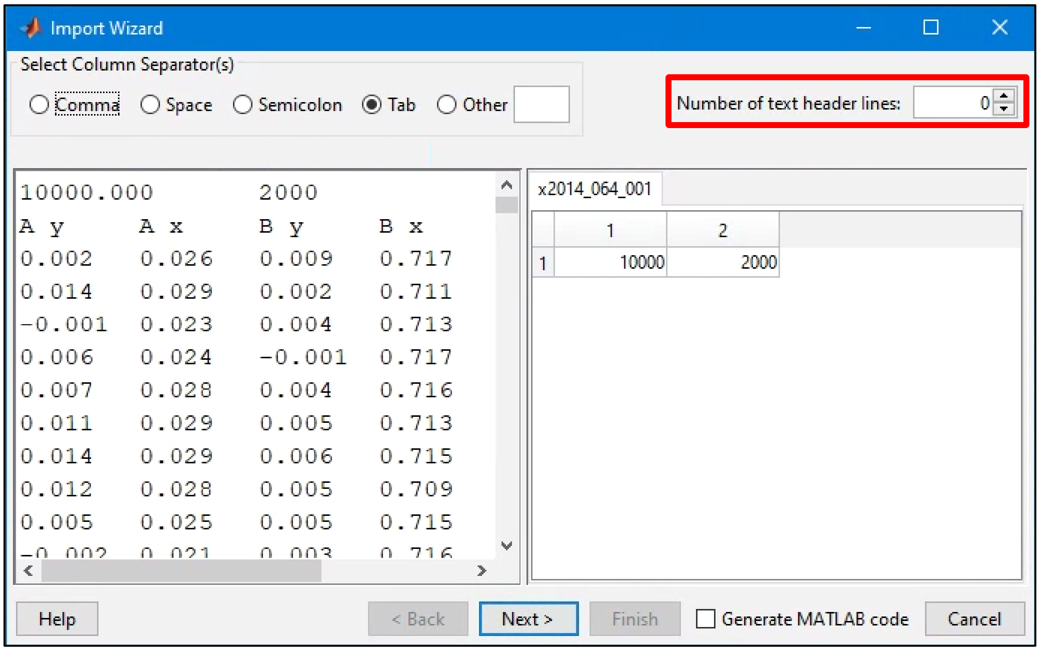

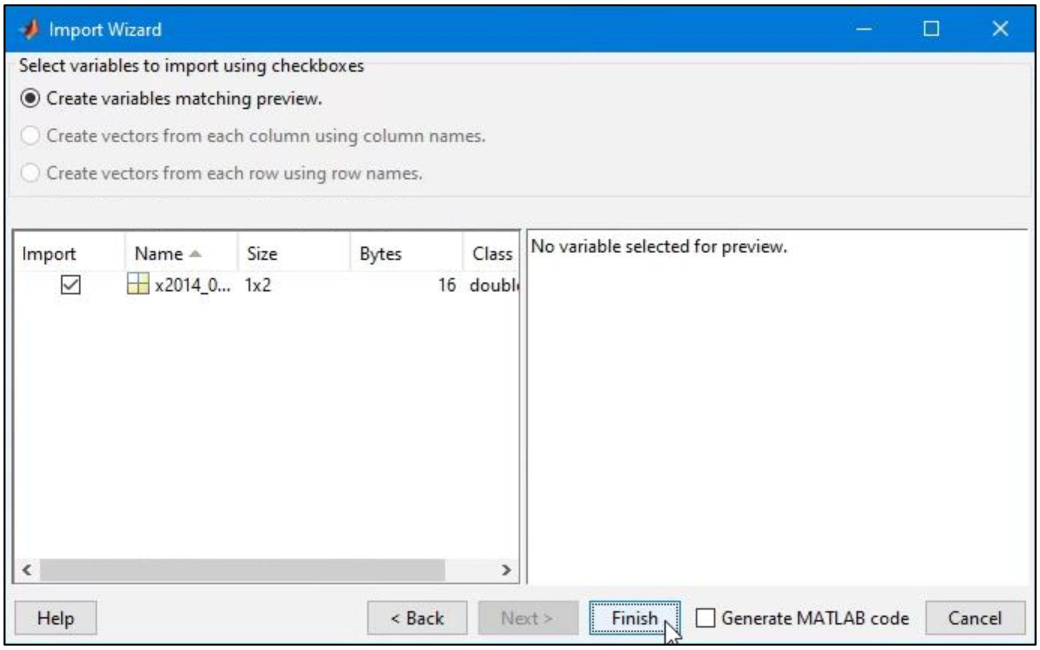

Next, an input dialog box will appear, prompting us to input values for the stiffnesses and volt-to-nanometer conversion factors.

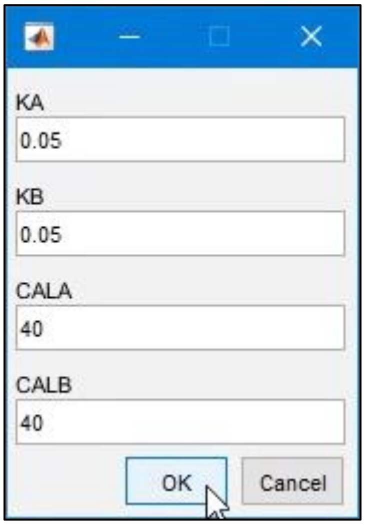

Finally, the **Import Tool** will again show. If we now say that this file contains two header lines, the **Import Tool** will start at the third line and read the entire file.

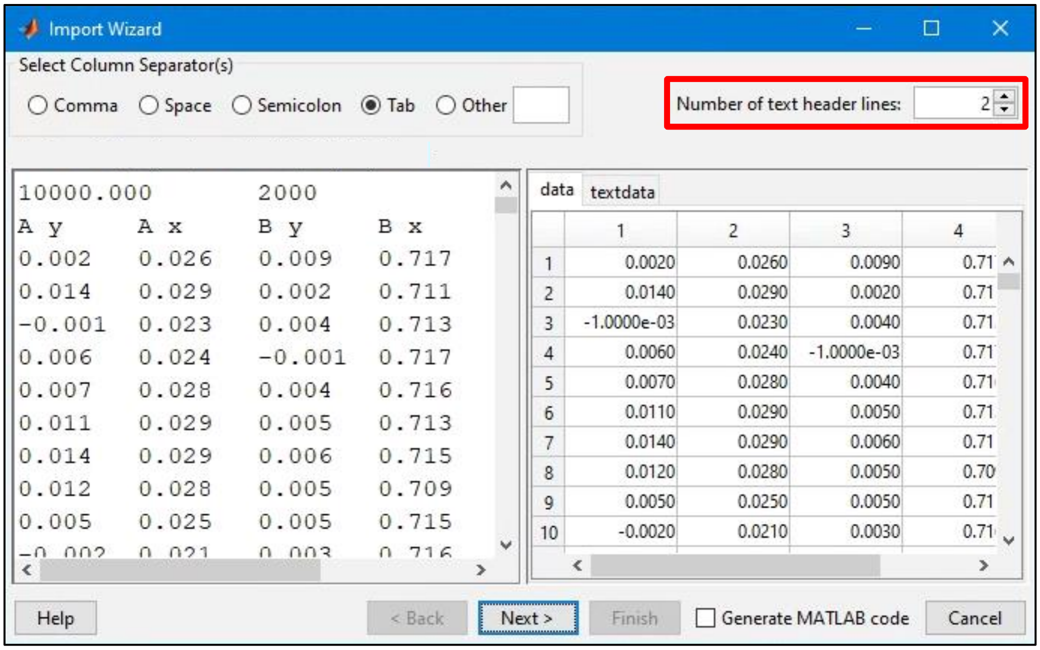

These two functions work with the example text file shown previously. They will not necessarily work with your data, but hopefully they will guide you in writing a function which does work. The user needs to make sure that the function used to read data is compatible with the format used to store the data.

#### Section 3: Custom Output Function

In Section 2 of the User Guide, it was explained that you can remove some of your data and, if desired, save the remaining data to a new text file. By default, **SPASM** will save this data in the format laid out in Section 1 of this appendix. However, you may want to save this data in a different format.

In the previous section, it was shown how you can define a custom MATLAB function for reading your data. This section shows how you can also define a custom MATLAB function for writing your data. As before, this function must satisfy certain requirements:

A. It must accept 7 input arguments. The 7 variables supplied to the function will be the 7 variables returned by **SPASM**'s input function **exactly as they were returned by that function**, in the same order. As a reminder, these variables are:
   1. The sampling frequency, in $s^{-1}$.
   2. The trap stiffness of bead A, in pN/nm.
   3. The trap stiffness of bead B, in pN/nm.
   4. The volt-to-nanometer conversion factor of bead A, in nm/V.
   5. The volt-to-nanometer conversion factor of bead B, in nm/V.
   6. A MATLAB table. Two of the columns of this table will be named 'BeadAPos' and 'BeadBPos'. The data in these columns give the positions of beads A and B, respectively, in units of volts. The number of rows in this table equals the number of data points, and the rows are ordered chronologically.
   7. The 7th output from the function's input function (see the previous section, **Custom Input Function**).

B. It must return one output argument, which will be printed directly to the top of your output text file. This variable can be anything, but it is recommended that this variable be a string or character vector containing your desired header and column names in the proper format, with tabs ('\t') separating columns of text and newline characters ('\n') separating rows of text.

Thus, you can use whatever header format and column names you would like for the newly generated text file. Consider again the following file.

```
10000.000       2000
A y     A x     B y     B x     Feedback
0.002   0.026   0.009   0.717   -0.001
0.014   0.029   0.002   0.711   0.001
-0.001  0.023   0.004   0.713   0.000
```

Had **SPASM** read this file with **myCustomInputFunction**, the first custom input function from the previous section, then our custom output function might look something like this:

```
function sHeader = myCustomOutputFunction(~, ~, ~, ~, ~, ~, header)
    colNames = {'A y', 'A x', 'B y', 'B x', 'Feedback'};
    sHeader = [sprintf('%s\t%s\n', header{1}{1}, header{2}{1})...
        sprintf('%s\t', colNames{1:end-1})...
        sprintf('%s\n', colNames{end})];
end
```

We create the character vector `sHeader` using the desired column names and the variable `header` which was returned by **myCustomInputFunction**. `header`, as a reminder, was a 1x2 cell array, containing the number of rows of data in `header{1}{1}` and the sampling frequency in `header{2}{1}`, both as character vectors. We place tabs ('\t') between columns and newline characters ('\n') between rows. Note that the first six inputs to the function are not needed and are thus replaced by '~'. This function will create a text file which looks *almost identical* to our input text file, except some rows of data will be missing, corresponding to the removed data.

Had **SPASM** instead read our file with **myCustomInputWhichUsesUI**, the second custom input function from the previous section, our custom output function needs to be edited slightly:

```
function sHeader = myCustomOutputWhichUsesUI(~, ~, ~, ~, ~, ~, header)
    colNames = {'A y', 'A x', 'B y', 'B x', 'Feedback'};
    sHeader = [sprintf('%.3f\t%d\n', header)... <- note the difference in this line
        sprintf('%s\t', colNames{1:end-1})...
        sprintf('%s\n', colNames{end})];
end
```

Whereas before `header` was a cell array which contained character vectors, here it is a double array (which contains doubles). *sprintf( )* can thus work directly with `header`. In addition, we must explicitly tell *sprintf( )* to format the number of rows with three decimal places.

In the previous section, the custom input function was specific to the format of the input text file. Here, the custom output function is specific not only to the format of the output text file but also to the input function. As before, the above two examples will not necessarily work with your data, but hopefully they give an idea of how to write a function which does work for you.

#### Appendix B: Generating Simulated Data

Alongside **SPASM**, we have provided another file, **simulator.m**. This text-based file allows you to create, visualize, and save simulated single molecule data.

There are a number of parameters within this file that you may wish to change, divided among the first four sections.

1. General parameters:

| Parameter name | Description | Value we use |
|---|---|---|
| `numEvents` | The total number of simulated actomyosin binding events. | `100` |
| `fs` | The sampling frequency, in $s^{-1}$. | `2000` |
| `plotTF` | Whether to plot the simulated data and the covariance. | `false` |
| `saveTF` | Whether to save the simulated data to a text file which can be analyzed by **SPASM**. | `true` |

2. Parameters pertaining to the Markov chain model:

| Parameter name | Description | Value we use |
|---|---|---|
| `trans` | An NxN matrix specifying the transition rates among the states, in $s^{-1}$, where N is the number of states. The $i_{th}$ row, $j_{th}$ column within `trans` gives the rate of transitioning from the $j_{th}$ state to the $i_{th}$ state. | $\begin{pmatrix} 0 & 0 & 4 \\ 0.5 & 0 & 0 \\ 0 & 70 & 0 \end{pmatrix}$ |
| `step_sizes` | A 1xN vector containing the position of the myosin during each state, in nm. | $(0 \;\; 4.7 \;\; 6.6)$ |
| `step_size_var` | A 1xN vector containing the variability in the step sizes during each state, in nm. | $(0 \;\; 0 \;\; 0)$ |
| `high_freq_noise_state` | A 1xN vector containing the amplitude of high frequency noise during each state, in nm. | $(25 \;\; 15 \;\; 15)$ |
| `low_freq_noise_A` | The amplitude of low frequency noise, in nm, which is applied uniformly throughout the data. | `0.5` |
| `low_freq_noise_f` | The frequency of low frequency noise, in $s^{-1}$, which is applied uniformly throughout the data. | `0.01` |

The system starts in the first state.

3. Coupling parameters:

| Parameter name | Description | Value we use |
|---|---|---|
| `AArBound` | A MATLAB anonymous function which returns the proportion of bead A's noise which is NOT shared with bead B, when myosin is bound. | `@() 1` |
| `BBrBound` | A MATLAB anonymous function which returns the proportion of bead B's noise which is NOT shared with bead A, when myosin is bound. | `@() 1` |

| | | |
|---|---|---|
| AArUnbound | A MATLAB anonymous function which returns the proportion of bead A's noise which is NOT shared with bead B, when myosin is unbound. | @() rand |
| BBrUnbound | A MATLAB anonymous function which returns the proportion of bead B's noise which is NOT shared with bead A, when myosin is unbound. | @() rand |

4. Covariance parameters:

| Parameter name | Description | Value we use |
|---|---|---|
| covwindow | Moving average filter window, for calculating the covariance. | 175 |
| covsmooth | 2nd order Savitzky-Golay filter, for smoothing the covariance. | 73 |

To run **simulator.m** in MATLAB, make sure this file is contained in your Current Folder or on your MATLAB path. You may need to move **simulator.m** to your Current Folder or change your Current Folder to whatever folder contains **simulator.m**. Then, either type *simulator* in the Command Window and press Enter

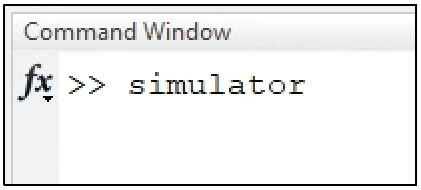

or open **simulator.m** in the Editor and click the green Run button.

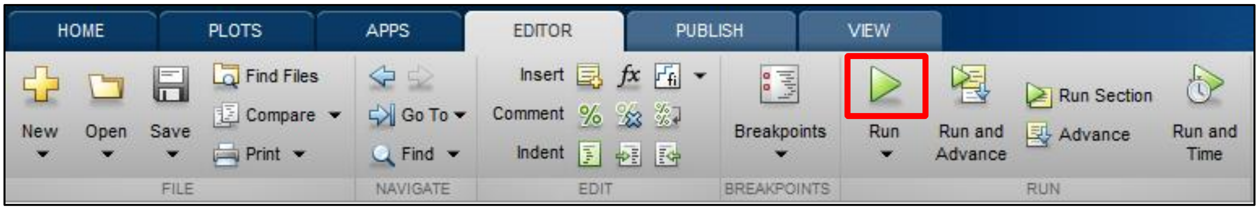

If you opted to plot the simulated data, two figures should appear. The first figure displays the bead positions over time, with the covariance plotted underneath. The second figure displays the covariance histogram. Handles to both figures are contained within a struct named `fig`.

If you opted to save the simulated data to a text file, a window should appear, prompting you to choose a filename and location for this file. The file produced by **simulator.m** will contain a column labeled **Key**. All non-zero data within this column correspond to times during which the simulated actomyosin is bound.

If you load the file into **SPASM**, data from the **Key** column is used to determine the locations of the true events. These locations are plotted on top of the beads' positions and on top of the covariance in blue, making it easy to see false positive and false negative events. It is also easy to compare the detected transition times with the true transition times.

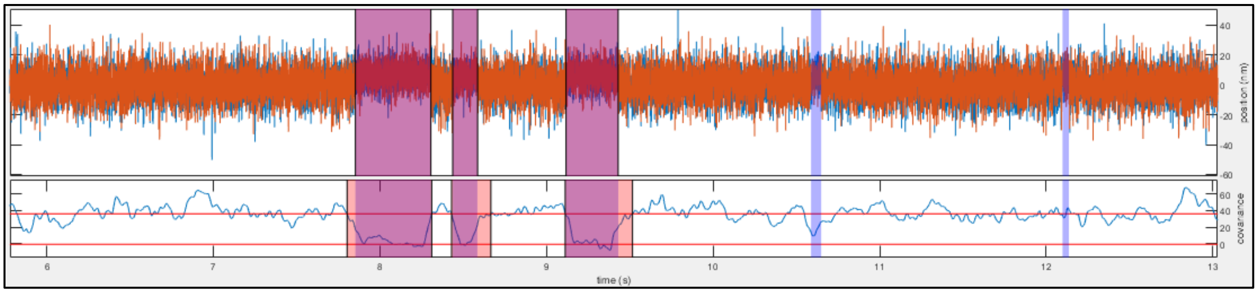

In the individual event plot, yellow vertical lines show all true transition times contained within the window.

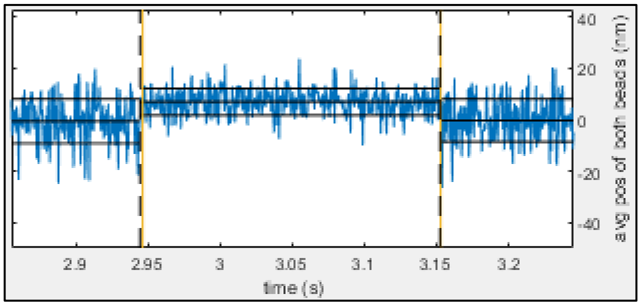

#### Appendix C: Optimizing the Window Sizes and Cutoffs

Our data analysis aims to avoid false positive and false negative events and accurately estimate the event start and end times. The change point algorithm is concerned with the latter of these. The former, on the other hand, is completely determined by the detection of events with the covariance and the exclusion of events by the minimum event duration and minimum event separation.

Thus, four parameters contribute to the number of false positive and false negative events, denoted $f_p$ and $f_n$ respectively:

1. The moving average window size, which is used to calculate the covariance.
2. The 2nd order Savitzky-Golay filter window size, which is used to smooth the covariance.
3. The minimum event duration, which is used to remove short events.
4. The minimum event separation, which is used to remove close neighboring events.

We were interested in studying cardiac myosin. Therefore, we generated a file of data to simulate this myosin. The rates of the simulated data matched the expected rates of this myosin, as determined by stopped flow experiments. It was assumed that values for the above four parameters which optimized analysis on this simulated data would also work well with experimental data obtained from this myosin.

After the simulated data was analyzed by **SPASM**, $f_p$ and $f_n$ were determined. To find false positive events, each detected event was mapped to the nearest real event. If a detected event did not overlap with any real events, it was counted as a false positive event. If multiple detected events were mapped to the same real event, all but the closest were counted as false positive events. Likewise, to find false negative events, each real event was mapped to the nearest detected event. If a real event did not overlap with any detected events, it was counted as a false negative event. If multiple real events were mapped to the same detected event, all but the closest were counted as false negative events.

The MATLAB function *simulannealbnd( )* was used to determine the values of the above four parameters which minimized the sum, $f_p+f_n$.

The same general approach can be used to estimate appropriate values for these parameters for any set of experimental data:

1. Generate a set of data which simulates the system measured by your experimental data.
2. Develop a method to detect false positive events among the detected events and to detect false negative events among the real events.
3. Use an optimization algorithm to determine values for the window sizes and cutoffs which minimize $f_p$ and $f_n$. You do not have to minimize the sum of $f_p$ and $f_n$, as we did. For example, you may care more about avoiding false positive events, in which case you might minimize a weighted average of $f_p$ and $f_n$ which heavily weights $f_p$.

If you are running the uncompiled version of **SPASM**, you can analyze your data without opening the user interface by using public methods and properties.

```
Command Window
>> methods SPASM

Methods for class SPASM:

SPASM                  getActiveDur           getBeadsToAnalyze      getTrimBeads
delete                 getActiveEvents        getForce               getTrimData
getActiveBeads         getActiveSep           getPosNStepSizes

Static methods:

calculateCov           defaultTxtInput        findPrelimEventsByMin
calculatePeaksMin      findChangepointWindows findRealEvents
changepoint            findOneChangepoint     findTwoChangepoints
checkInstall           findPrelimEvents       scoreDetectedEvents

Methods of SPASM inherited from handle.
```

```
Command Window
>> properties SPASM

Properties for class SPASM:

    allEvents
    activeEvents
    minSep
    minDur
    autop1
    autop2
    automin
```

You can use MATLAB's *help( )* function for more information about each method and property.

```
Command Window
>> help SPASM.calculatePeaksMin
  calculatePeaksMin Calculates the two peaks of a bimodal histogram, as well
  as the minimum value between the two peaks.
    [p1, p2, m] = calculatePeaksMin(edges, values) returns the lower peak p1,
    the upper peak p2, and the minimum m of the histogram defined by inputs
    edges and values.

    See also histcounts.
```

```
Command Window
>> help SPASM.autop1
 autop1 -  The lower peak of the covariance histogram, which is used in the peak-to-peak method.
```

#### Appendix D: Version of SPASM Requiring Only One Bead

If only one of the optically trapped beads is monitored, **SPASM** is unable to calculate the covariance between the two beads' positions. In this case, the variance of the monitored bead can be used in place of the covariance. We have created a version of **SPASM** which only requires one bead, using the variance instead of the covariance. This version, named **SPASM_one_bead**, can be found at https://github.com/GreenbergLab/SPASM.

Most aspects of **SPASM_one_bead** remain unchanged from **SPASM**, but there are a handful of differences. These differences are described in the following sections.

#### Section 1: Data Format

The required data format for **SPASM** is detailed in Appendix A, Section 1. The general layout for **SPASM_one_bead** is unchanged:

| | | | | |
|---|---|---|---|---|
| key1 | value1 | | | header |
| ... | ... | | | |
| keyN | valueN | | | |
| colName1 | colName2 | ... | colNameM | column names |
| data(1,1) | data(1,2) | ... | data(1,M) | data |
| ... | ... | ... | ... | |
| data(L,1) | data(L,2) | ... | data(L,M) | |

Again, columns of text are tab-delimited. As before, the following must be true:

(1) All data must be numeric text (e.g. '1.0'), and the first row of data must be the first row in the file which contains only numeric text.

However, the following requirements are different:

(2) There must be a key labeled 'K1' **or** 'K', and there must be a key labeled 'CAL1' **or** 'CAL'. There must also be a key labeled 'Sample Rate'. All three of these keys must have corresponding numeric values. These values determine the trap stiffness for the bead (K1 or K), in units of pN/nm, the volt-to-nanometer conversion factor for the bead (CAL1 or CAL), in units of nm/V, and the sampling frequency (Sample Rate), in $s^{-1}$.

(3) One of the column names must equal 'Trap1X' **or** 'TrapX'. The corresponding column will be treated as the position of the bead, in units of volts.

When data are saved to a new .txt file (see User Guide Section 2, Save), the new file will have a format compatible with **SPASM_one_bead**. The program will save the trap stiffness using the key 'K1' and the volt-to-nanometer conversion factor using the key 'CAL1'. The bead's position will be saved in a column named 'Trap1X'.

#### Section 2: Custom Input Function

As before, if you are running the uncompiled version of **SPASM_one_bead**, you may supply your own function for reading text files. The requirements for this function are slightly different:

A. It must accept one input argument. This will be a character vector containing the full path of the input text file.
B. It must return 5 output arguments. These are, in order:
    1. The sampling frequency, in $s^{-1}$.
    2. The trap stiffness of the bead, in pN/nm.
    3. The volt-to-nanometer conversion factor of the bead, in nm/V.
    4. A MATLAB table. One of the columns of this table must be named ‘BeadPos’. The data in this column are treated as the position of the bead, in units of volts. The number of rows in this table equals the number of data points, and the rows are ordered chronologically.
    5. A $5^{th}$ variable which gives the text file’s header. This variable is not used within the program and does not affect data analysis. It only affects how your data will be saved to a text file, should you choose to save it (see User Guide Section 2, Save). For more information, see the next section, **Custom Output Function**.

Refer to Appendix A, Section 2 for more information.

#### Section 3: Custom Output Function

As before, if you are running the uncompiled version of **SPASM_one_bead**, you may supply your own function for writing your data to a text file. The requirements for this function are slightly different:

A. It must accept 5 input arguments. The 5 variables supplied to the function will be the 5 variables returned by the program's input function **exactly as they were returned by that function**, in the same order. As a reminder, these variables are:
   1. The sampling frequency, in $s^{-1}$.
   2. The trap stiffness of the bead, in pN/nm.
   3. The volt-to-nanometer conversion factor of the bead, in nm/V.
   4. A MATLAB table. One of the columns of this table will be named 'BeadPos'. The data in this column give the position of the bead, in units of volts. The number of rows in this table equals the number of data points, and the rows are ordered chronologically.
   5. A $5^{th}$ variable containing the text file's header.

B. It must return one output argument, which will be printed directly to the top of your output text file. This variable can be anything, but it is recommended that this variable be a string or character vector containing your desired header and column names in the proper format, with tabs ('\t') separating columns of text and newline characters ('\n') separating rows of text.

Refer to Appendix A, Section 3 for more information.

#### Section 4: Select One Bead for Analysis

In Section 3d of the User Guide, it was shown how you can choose to analyze only one of the two trapped beads. In **SPASM**, by default, the average position between bead A and bead B is analyzed by the change point algorithm and used to generated ensemble averages. In **SPASM_one_bead**, as there is only one bead, you no longer have the option to specify whether one bead or both beads are used in analysis.

#### Section 5: Save to Excel

The process for saving data to an Excel sheet is unchanged. Refer to Section 9 of the User Guide for details. However, **SPASM_one_bead** saves slightly different values. After saving data to an Excel sheet, that sheet will include

I. general information about the analyzed data, such as the full path of the .txt file, the sampling frequency, and the number of analyzed events;

II. all of the parameters used in data analysis, such as the window sizes or the covariance histogram peak locations, so that the data analysis can be repeated; and

III. a table which contains, for each event,

- the event start and end times, in both points and seconds;
- the duration of the event, in both points and seconds;
- the amount of time separating the event from its neighboring events, in both points and seconds;
- the position of the bead just before and just after the event start, averaged over 10 ms windows;
- the position of the bead just before and just after the event end, averaged over 10 ms windows;
- the estimated substep 1, substep 2, and total step sizes based on the bead's position;
- the force on the bead during the event, averaged over the entire event;
- the force on the bead before and after the event, averaged over the entire time separating the event from its neighboring events; and
- the force on the bead just before and just after the event end, averaged over 10 ms windows.
